# Ethylene is a local modulator of jasmonate-dependent phenolamide accumulation during *Manduca sexta* herbivory in *Nicotiana attenuata*

**DOI:** 10.1101/2020.07.21.213371

**Authors:** Florent Figon, Ian T. Baldwin, Emmanuel Gaquerel

## Abstract

Rapid reconfigurations of intimately connected phytohormone signaling networks allow plants to tune their physiology to constantly varying ecological conditions. During insect herbivory, most of the induced changes in defense-related leaf specialized metabolites are controlled by jasmonate (JA) signaling, which, in the wild tobacco *Nicotiana attenuata*, recruits MYB8, a transcription factor controlling the accumulation of phenolic-polyamine conjugates (phenolamides). In this and other plant species, herbivory also locally triggers ethylene (ET) production but the outcome of the JA-ET crosstalk at the level of secondary metabolism regulation has remained superficially investigated. Here, we analyzed local and systemic herbivory-induced changes by mass spectrometry-based metabolomics in leaves of transgenic plants impaired in JA, ET and MYB8 signaling. Parsing deregulations in this factorial data-set identified a network of JA/MYB8-dependent phenolamides for which impairment of ET signaling attenuated their accumulation only in locally-damaged leaves. Further experiments revealed that ET, albeit biochemically interrelated to polyamine metabolism via the intermediate *S*-adenosylmethionine, does not impart on free polyamine levels, but instead significantly modulates phenolamide levels and marginally affected transcript levels in this pathway. This study identifies ET as a local modulator of phenolamide investments and provides a metabolomics data-platform to mine associations between herbivory-induced signaling and specialized metabolite groups in *N. attenuata*.

**Summary statement:** Herbivory-elicited ethylene acts as a local enhancer of the production of jasmonate/MYB8-dependent phenolamides during the defense response of *Nicotiana attenuata* leaves against herbivory by the lepidopteran larva *Manduca sexta*.

## Introduction

Plants, as sessile organisms, must continuously adjust their physiology to cope with fluctuating abiotic and biotic conditions. To resist insect herbivory, plants have evolved efficient constitutive as well as inducible defense strategies, the latter being thought to mitigate resource allocation costs to defense during a plant’s ontogeny (Heil & Baldwin 2002). Defense induction translates from the activation and complex interactions among phytohormone signal transduction pathways, which are triggered in attacked and distant leaves upon insect feeding (Erb, Meldau & Howe 2012).

Activation of ethylene (ET) signaling is part of the hormonal response of plants against attackers. ET biosynthesis involves the two-step transformation of *S*-adenosylmethionine (SAM) into ET by ACS (1-Aminocyclopropane-1-Carboxylate [ACC] Synthase) and ACO (ACC oxidase) (Figure 1a). The interaction between ET and the ETR1 receptor protein, leads to the activation of ET-dependent transcriptional responses (Figure 1a) (Stepanova & Alonso 2009). This signaling pathway is known to contribute to the regulation of induced defenses in the wild tobacco *Nicotiana attenuata* after attack by the specialized larva of *Manduca sexta* (von Dahl *et al*. 2007; Onkokesung *et al*. 2010b). Conversely, the performance of this herbivore is strongly improved when feeding on *N. attenuata* lines deficient in ET signaling (Onkokesung, Baldwin & Gális 2010a). In tobacco species, ET signaling is well known to antagonize the jasmonate-dependent accumulation of the defensive neurotoxin nicotine. This regulatory response is thought as a mechanism to adjust nitrogen investments into costly nicotine production and requires several ERF (Ethylene Response Factor) transcription factors part of the NIC regulatory loci positively controlling nicotine biosynthetic genes (Shoji, Kajikawa & Hashimoto 2010). *M. sexta* larvae are known to be tolerant to high nicotine levels (Onkokesung *et al*. 2010b). Hence, the important increases in *M. sexta* performance that were previously detected when feeding on ET signaling deficient plants likely translate from more profound and yet-to-explored deregulations of the host plant defense metabolism. In this respect, cross-talks between ET and other herbivory-induced hormonal sectors have been characterized at the molecular level (Wang, Li & Ecker 2002) and are likely to account for the deregulated defense response in *N. attenuata* lines rendered genetically insensitive to ET perception (von Dahl *et al*. 2007; Li, Halitschke, Baldwin & Gaquerel 2020).

**Figure 1.**
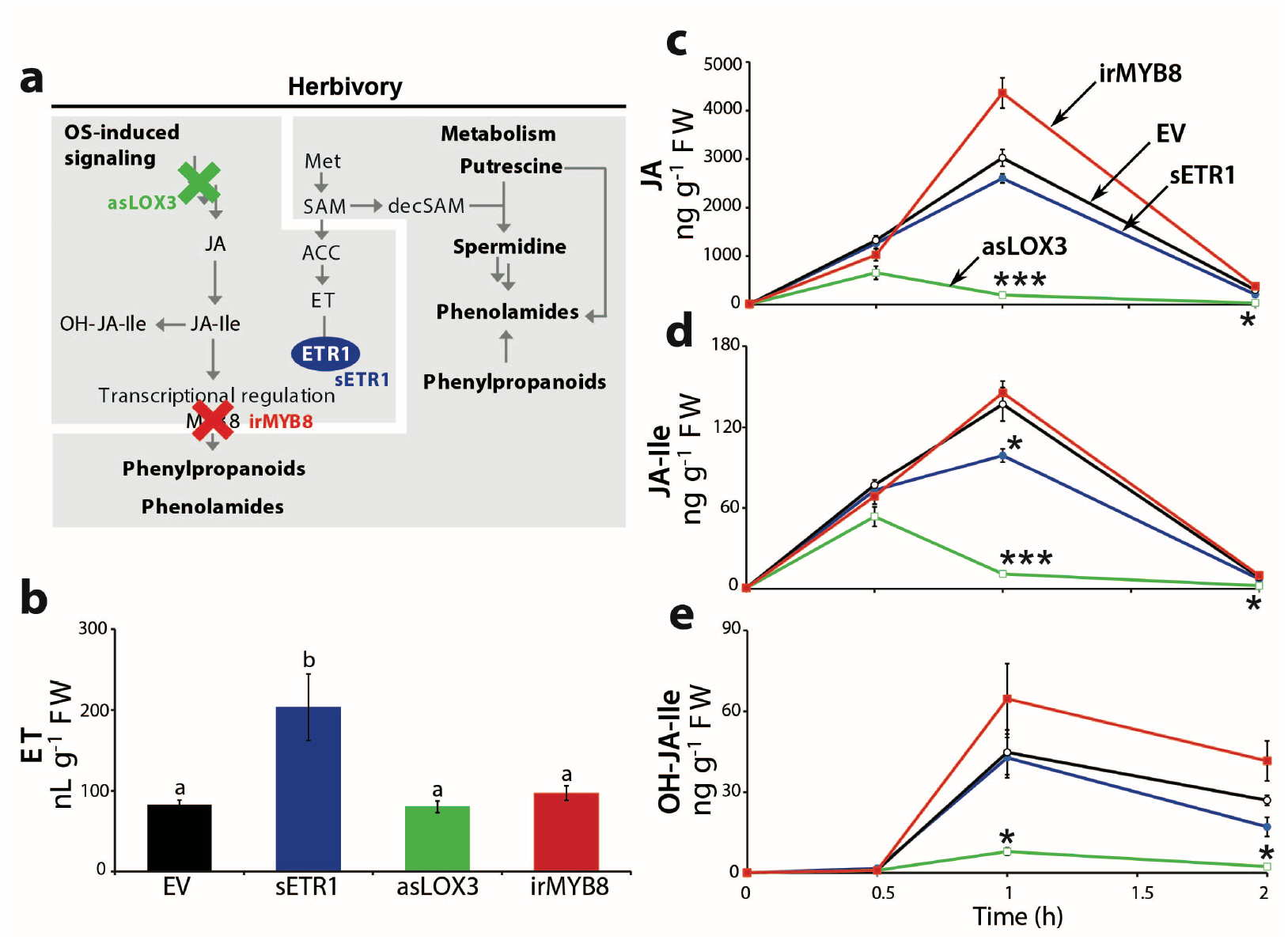
Genetic manipulations of herbivory-induced jasmonate and ethylene accumulation in leaves of *Nicotiana attenuata*. **(a)** Simulated leaf herbivory by *Manduca sexta* herbivory elicits the production of ethylene (ET) and jasmonates in *Nicotiana attenuata* and dependent signaling and metabolic reconfigurations. Three different *N. attenuata* transgenic lines were used for studying interactions between these hormonal pathways: (*i*) an asLOX3 line in which the first oxidative step for jasmonate biosynthesis is impaired, (*ii*) an sETR1 line expressing a nonfunctional mutated version of ETR1, the ET receptor and (*iii*) an irMYB8 line with silenced transcripts for MYB8, the core JA-dependent transcription factor controlling gene expression for defensive phenylpropanoid-polyamine conjugates (phenolamides) accumulation. Ethylene synthesis requires from *S*-adenosylmethionine (SAM), whose decarboxylated form (decSAM, decarboxylated-*S*-adenosylmethionine) is required within the polyamine biosynthetic pathway. **(b)** Mean levels (± SE, 5 biological replicates) of ET emission produced from leaf discs of the different genotypes accumulated over 10h after simulated herbivory by mechanical wounding and application of *M. sexta* oral secretions (W+OS). Different letters indicate significant differences (P ≤ 0.05, ANOVA followed by Tukey HSD post-hoc tests) between genotypes. **(c-e)** Mean levels (± SE, 5 biological replicates) of JA, JA-Ile and OH-JA-Ile in leaves elicited by W+OS treatment. Asterisks indicate significant differences compared to an empty vector (EV) control at specific time points (* P ≤ 0.05, *** P ≤ 0.001, one-way ANOVA followed by Tukey HSD post-hoc tests; when normality assumption was not met, Kruskal-Wallis and pairwise Wilcoxon rank sum tests were applied). ACC, 1-aminocyclopropane-1-carboxylic acid; decSAM, decarboxylated-*S*-adenosylmethionine; JA-Ile, JA-Isoleucine; OH-JA-Ile, hydroxy-JA-Ile; Met, methionine.

Jasmonate signaling exerts a central regulatory function for most of the transcriptional changes and metabolic reconfigurations contributing to induced responses to insect herbivory (Wasternack & Hause 2013). In *N. attenuata, M. sexta* feeding elicits a rapid burst of all jasmonate biosynthetic intermediates, leading to transient and herbivory-specific accumulations of both jasmonic acid and its bioactive conjugate JA-Isoleucine (JA-Ile) (Paschold, Bonaventure, Kant & Baldwin 2008; Kallenbach, Alagna, Baldwin & Bonaventure 2010). A large body of research on this ecological interaction has shown that JA-Ile signaling is critical for the efficient activation of an anti-herbivore arsenal (reviewed in Wang & Wu 2013), which includes the above mentioned nicotine, trypsin proteinase inhibitors acting as anti-digestive proteins (Kang, Wang, Giri & Baldwin 2006), as well as a broad set of phenolamides whose contribution to anti-herbivore defenses was more recently uncovered (Onkokesung *et al*. 2011). Several studies have reported convergent evidence for a key role played by the interplay between JA and ET signaling pathways in the adjustment of appropriate defense investments to different caterpillar species (reviewed in Erb *et al*. 2012). Nevertheless, besides the well-characterized function of ET in tuning the JA-induced nicotine production, its regulatory role over other classes of defense metabolites has remained poorly examined. In the case of phenolamides, a recent large-scale metabolomics study, which tested predictions of seminal hypotheses on the remodeling of specialized metabolite production during insect herbivory, has detected that ET signaling finely tunes the metabolic response of different *Nicotiana* species according to the herbivore species (Li *et al*. 2020).

Phenolamides, which result from the conjugation of hydroxycinnamic units to a polyamine skeleton (Figure 1a), are important constitutively-produced and/or stress-inducible metabolites in the vegetative tissues of many plant species, including rice and *N. attenuata* (Onkokesung *et al*. 2011; Alamgir *et al*. 2016). In *N. attenuata*, the biosynthesis of these defensive specialized metabolites is strongly induced by the attack of *M. sexta*. One of these compounds, *N*-caffeoylputrescine, has been rigorously demonstrated to negatively affect *M. sexta* performance (Kaur, Heinzel, Schottner, Baldwin & Galis 2010). Similar to the well-characterized nicotine regulation, elevations in phenolamide levels also impart significant trade-offs for nitrogen allocation during insect herbivory and hence necessitates fine-tuning mechanisms, which remain largely unknown (Ullmann-Zeunert *et al*. 2013). From a biosynthetic standpoint, conjugation of CoA-activated hydroxycinnamic acid units onto polyamine skeletons translates from the enzymatic activities of a diverse array of BAHD *N*-acyltransferases, with AT1, DH29 and CV86 being the three BAHD controlling most of the phenolamide production in *N. attenuata* (Onkokesung *et al*. 2011). Briefly, AT1 has been shown to control the acylation of putrescine with coumaroyl-CoA or caffeoyl-CoA. DH29 was demonstrated to catalyze the first acylation step on spermidine, while CV86 is involved in the production of certain diacylated spermidine isomers from DH29-dependent mono-acylated spermidines, suggesting that additional *N*-acyltransferases are likely required to produce the full spectrum of phenolamides found in *N. attenuata*. The expression of *AT1, DH29* and *CV86* is controlled by the transcription factor MYB8, a master regulator of phenolamide biosynthesis, as evidenced by the presence of MYB8 binding sites in the promoters of those genes (Schäfer *et al*. 2017) and by the fact that MYB8-deficient plants are virtually free of phenolamides (Kaur *et al*. 2010; Onkokesung *et al*. 2011).

The present study aimed at testing whether herbivory-induced ET signaling modulates the jasmonate- and MYB8-dependent production of defensive phenolamides in *N. attenuata* leaves attacked by *M. sexta*. To comprehensively examine interplays between JA and ET signaling sectors over herbivory-induced metabolic reconfigurations, we analyzed local and systemic herbivory-induced changes in leaves of transgenic plants impaired in JA (asLOX3), ET (sETR1) and MYB8 (irMYB8) signaling using mass spectrometry-based metabolomics. The attack from *M. sexta* was simulated by the application of oral secretions (OS) from caterpillars on the leaves. From the exploration of metabolic class-specific deregulations in this factorial data-set, we detected that ET signaling is essential for the intact production of phenolamides in locally-damaged leaves. Further, we examined whether a possible metabolic cross-talk, via the shared intermediate *S*-adenosylmethionine (SAM), between the production of ET and free polyamine metabolic homeostasis could have an impact on the channeling of putrescine and spermidine units for phenolamide production at the site of attacks. Quantitative analyses of free polyamines and main phenolamides revealed that herbivory-induced ET biosynthesis does not destabilize polyamine homeostasis, but instead marginally affected transcript levels *(MYB8* and *CV86*) in this pathway. Altogether, our results indicate that ET is a local regulator of defensive phenolamides during herbivory.

## Materials and methods

### Plant material

Selected transgenic lines with suppressed transcript levels for key genes in the herbivory-induced transduction cascade of *N. attenuata* were employed in this study. An empty vector (EV) construct was used as control genetic background in most experiments. The phenotypic and molecular characterization of each of these lines has been published previously. Briefly, sense *ETR1 (ETHYLENE RECEPTOR1*, sETR1) plants ectopically express a non-functional version of ETR1 (von Dahl *et al*. 2007). These plants are ethylene-insensitive and the negative feedback exerted by ethylene perception over its own production is disrupted, resulting in higher ethylene emissions after attack from *M. sexta* in this transgenic line (Figure 1b). Antisense *LOX3 (LIPOXYGENASE3*, asLOX3) plants constitutively express a fragment of the *LOX3* gene in an antisense orientation, which leads to the silencing of endogenous *LOX3* transcripts by RNAi (Halitschke & Baldwin 2003). asLOX3 plants are impaired in the first oxidation step within the JA biosynthetic pathway, thereby leading to a strong depletion of the herbivory-elicited jasmonate bursts. Inverted repeat *MYB8* (irMYB8) plants constitutively express inverted repeat fragments of the *MYB8* gene, which results in the silencing of endogenous *MYB8* by RNAi (Kaur *et al*. 2010). Phenolamide biosynthesis, which is transcriptionally coordinated by the action of MYB8, is almost completely abolished in this transgenic line. The irACO transgenic line used in the experiment presented in Figure S3 exhibits constitutively repressed transcript levels for the ethylene biosynthesis gene *ACO* (*1-AMINOCYCLOPROPANE-1-CARBOXYLIC ACID OXIDASE*), thereby resulting in a strong impairment of ethylene production.

All seeds were first sterilized and incubated with diluted smoke and 0.1 M GA3, as previously described (Krügel, Lim, Gase, Halitschke & Baldwin 2002), and then germinated on plates containing Gamborg B5 medium. Ten-day-old seedlings were transferred to small pots (TEKU JP 3050 104 pots; Pöppelmann) with Klasmann plug soil (Klasmann-Deilmann), and after 10 d, seedlings were transferred to 1-L pots. Plants were grown in the glasshouse with a day/night cycle of 16 h (26°C – 28°C)/8 h (22°C – 24°C) under supplemental light from Master Sun-T PIA Agro 400 or Master Sun-TPIA Plus 600-W sodium lights (Philips Sun-T Agro). All experiments were conducted with 40 to 45-day old rosette-stage plants.

### Simulated herbivory treatment and experimental design

Oral secretions (OS) from 3^rd^ instar *M. sexta* larvae were collected with a pipette, pooled and kept under argon at −20°C until further use. At the start of the elicitation procedure, one transition leaf (youngest fully-expanded leaf, referred as leaf position 0) from each plant was rapidly harvested, flash-frozen in liquid nitrogen and later used as pre-treatment controls. For each plant, two rosette leaves referred to as local leaves (leaf positions +1 and +2 in Figure 2a), were mechanically wounded with a fabric pattern wheel and 20 μL of 1/8^th^ water-diluted OS were directly applied on the wounds. This procedure hereafter referred to as W+OS, has been shown in previous studies to recapitulate a large proportion of the reconfigurations detected at the level of early signaling events, transcriptome and metabolome dynamics during direct *M. sexta* herbivory (Halitschke, Schittko, Pohnert, Boland & Baldwin 2001). Two younger sink leaves (leaf positions − and −2 in Figure 2a) growing above locally treated leaves and with a minimal angular distance were collected as untreated systemic leaves and pooled prior metabolite extraction.

**Figure 2.**
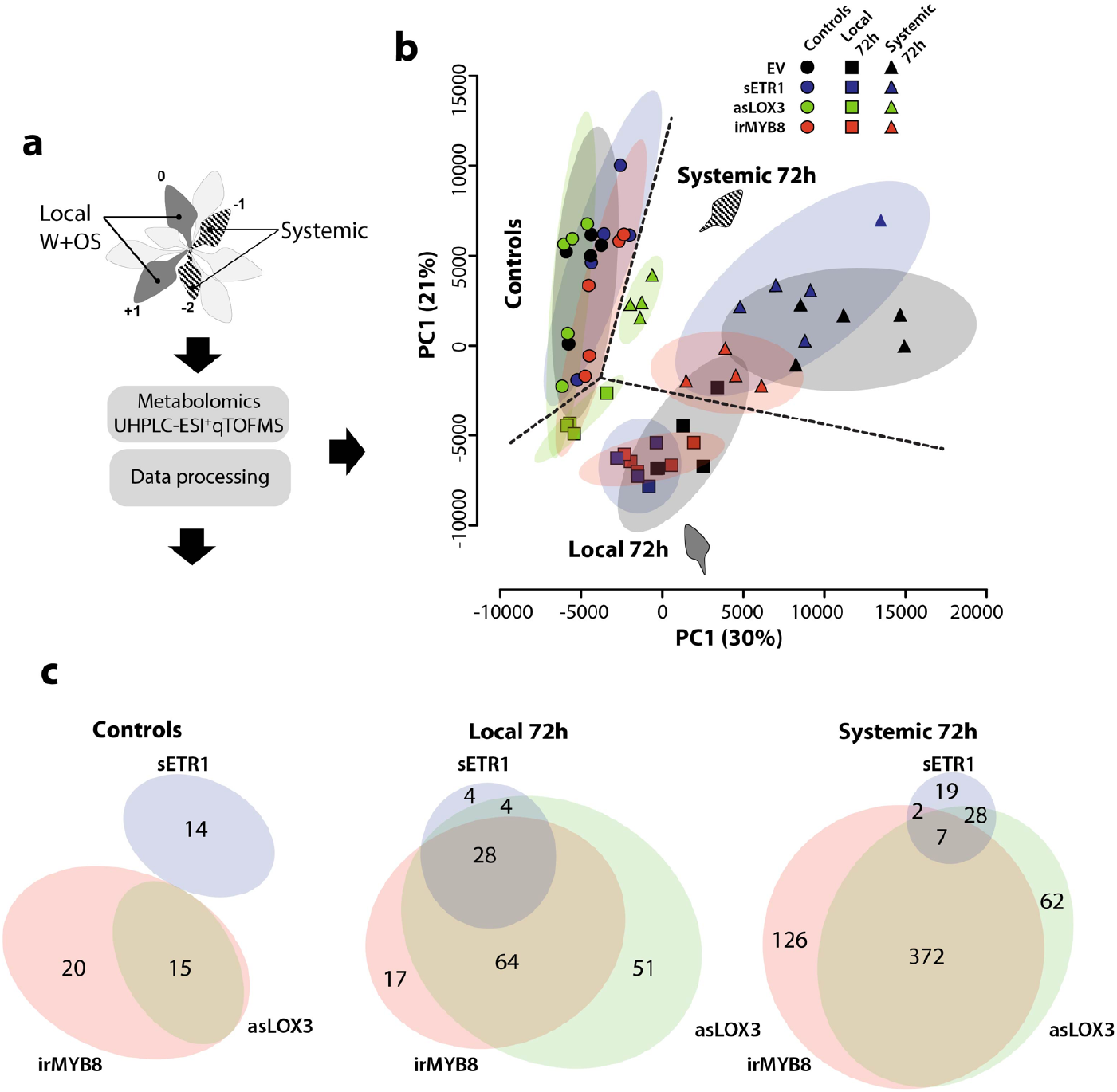
Untargeted MS-based metabolomics detects distinct herbivory-induced reconfigurations in local and systemic tissues and a major overlap between ethylene-, jasmonic acid- and MYB8-dependent responses in locally-elicited leaves. **(a)** Elicitation scheme and metabolomics workflow to parse local and systemic secondary metabolism to simulated insect herbivory. Two fully expanded rosette leaves were mechanically wounded and treated with diluted oral secretions (W+OS) from *Manduca sexta*. Untreated systemic leaves growing, with minimal angular distance, above elicited leaves were also collected in the four different genotypes (empty vector [EV], asLOX3, sETR1 and irMYB8) at specific time points after simulated herbivory. Leaf methanolic extracts were subjected to UHPLC-qTOF-MS analysis. MS data were post-processed XCMS and CAMERA R packages to generate a consistent *m/z* feature matrix analyzed by univariate and multivariate statistics. **(b)** Score plot generated for the two first principal components (PCs) derived from a principal component analysis using a autoscaled matrix for *m/z* feature peak intensities extracted for measurements of control (time point 0) and local/systemic W+OS elicitation (72 h) samples. Dashed ellipses represent 95% Hostelling confidence intervals. **(c)** Venn diagram visualization of overlapping and non-overlapping significantly deregulated *m/z* features (sampling/tissue-type-specific one-way ANOVA followed by Tukey HSD post-hoc tests, P ≤ 0.01) in sample measurements of control, local and systemic leaves of each of the transgenic line compared to in the corresponding EVs. Ellipses’ size and overlapping areas are respectively proportional to the total number of transgenic line-level deregulated *m/z* features and numbers of overlapping *m/z* features between transgenic lines.

In all experiments, 5 independent plants for each sampling time-points were used resulting in 5 biological replicates. Briefly, samples for specialized metabolite profiling were harvested for both local and systemic leaf positions at 0 (control untreated transition leaf, see above), 24, 48 and 72 h post-elicitation. The same experimental design was employed for the harvest of local and systemic leaf samples in order to monitor dynamics of free polyamines, with the exception that additional sampling times (0.5, 1, 2, 4 and 6 h) were considered to detect early changes to the W+OS treatment. Finally, to investigate changes in peaking levels of phytohomones across the different transgenic lines, only local leaves harvested 0, 0.5, 1, 1.5 and 2 h after starting the elicitation were considered for analysis.

### Quantitative real-time PCR analysis

Total RNA from 100 mg of ground tissues was extracted as described by (Linke *et al*. 2002). RNA was then treated with DNase using the DNA-free kit from Ambion following the manufacturer’s protocol. cDNA synthesis was performed using the RevertAid H Minus Reverse Transcriptase from ThermoScientific and a poly-T primer according to the manufacturer’s protocol. RT-qPCR was performed with a Stratagene MX3005P instrument. Transcript levels were calculated with a calibration curve of cDNA. For normalization, transcript levels of the housekeeping gene *ELF1a* were used (*EF1-α*, Acc. No. D63396). Untreated leaf controls harvested at the start of the elicitation procedure were used to examine basal transcript levels. To assess cross-genotype variations in W+OS-induced transcript levels, local leaves harvested 2 h after elicitation were considered since a previous transcriptomics study has shown that the focal transcripts reached maximum levels for this time-point. Primers are summarized in Table S1.

### Quantification of herbivory-induced ethylene emissions

Leaf discs were obtained from untreated plants for ethylene quantification. Directly after collection, these leaf discs were wounded with a pattern wheel, treated with 2 μl of 1/8^th^ water-diluted OS, and incubated during 10 h into 4 mL sealed glass-vials. The ethylene accumulated in the glass vials was quantified with a photoacoustic spectrophotometer (http://www.sensor-xsense.nl) operating at the ET-specific excitation frequency of 1640 Hz in the sample mode.

### Quantification of jasmonates, ABA and SA by HPLC coupled to triple-quadrupole mass spectrometry

Phytohormones were extracted following the protocol described in (Wu, Hettenhausen, Meldau & Baldwin 2007) with small changes. 200 mg of finely ground tissues were added to 1 mL ethyl acetate spiked with 200 ng of D2-JA and 40 ng of D6-ABA, D4-SA and JA-^13^C6-Ile as internal standards and metal balls were added. The samples were homogenized in a ball mill (Genogrinder 2000; SPEX CertiPrep) for 45 s at 250 strokes/min. Homogenized samples were centrifuged at 16,000 g, 4°C for 30 min. Each pellet was re-extracted with 0.5 mL of ethyl acetate and centrifuged. Supernatants were combined and dried in a SpeedVac, and finally resuspended in 0.5 mL of 70% methanol before centrifugation. Phytohormones were detected by high pressure liquid chromatography coupled to a triple quadrupole mass spectrometer as described in (Stitz *et al*. 2011). Chromatography was performed on an Agilent 1200 HPLC system (Agilent Technologies) using a Zorbax Eclipse XDB-C18 column (50 x 3 x 4.6 mm, 1.8 μm; Agilent). Formic acid (0.05%) in water and acetonitrile were employed as mobile phases A and B, respectively. The elution gradient was based on a binary solvent [time/concentration (min/%) for B]: 0 min/5%, 0.5 min/5%, 9.5 min/42%, 9.51 min/100%, 12 min/100% and 15 min/5%. The solvent A consisted in deionized water with 0.05% formic acid. The solvent B consisted in acetonitrile. The flow rate was constant at 1.1 mL/min and the column temperature was set at 25°C. An API 3200 tandem mass spectrometer (Applied Biosystems) equipped with a Turbospray ion source was operated in the negative ionization mode. Parent-to-product ion transitions specific to each hormone were monitored: *m/z* 136.9 to 93.0 for SA, *m/z* 140.9 to 97.0 for D4-SA, *m/z* 209.1 to 59.0 for JA, *m/z* 213.1 to 56.0 for D2-JA, *m/z* 263.0 to 153.2 for ABA, *m/z* 269.0 to 159.2 for D6-ABA, *m/z* 322.2 to 130.1 for JA-Ile and OH-JA-Ile, *m/z* 328.2 to 136.1 for JA-^13^C6-Ile. Quantification was achieved by integrating peak areas of natural phytohormones and comparing them to those of their respective internal standards.

### Polyamines and tyramine quantification by HPLC-FLD

Free polyamines of ground leaf tissues were quantified using a method described in (Docimo *et al*. 2012). Approximately 50 mg of ground tissues were extracted with 0.1 N HCl and metal balls were added. The samples were homogenized in the ball mill (Genogrinder 2000; SPEX CertiPrep) for 45 s at 250 strokes/min. Homogenized samples were centrifuged at 16,000 g, 4°C for 30 min. Supernatants were re-centrifuged as before. A supernatant aliquot was added to a borate buffer solution (pH 11). 50 μL of this mix was first derivatized with 50 μL of OPA (*ortho*-phthalaldehyde) for 1 min, and then a second derivatization with FMOC (fluorenylmethyloxycarbonyl chloride) was applied for 2 min. Polyamines were detected by the fluorescence of OPA molecules bound to primary amines (excitation = 340 nm, emission = 445 nm). The elution gradient was based on a binary solvent [time/concentration (min/%) for B]: 10 min/100%, 13 min/100%, 13.1 min/50% and 14.6 min/50%. Solvent A was deionized water with 0.05% formic acid. Solvent B was pure acetonitrile. The flow rate was constant at 0.8 mL/min and the column temperature was set at 25°C. Quantification of putrescine, spermidine and tyramine was achieved using external calibration curves built from the measurement of authentic standards.

### Phenolamide quantification by HPLC-PDA

Phenolamides extracted from ground leaf tissues were quantified with a HPLC-photodiode array (PDA) detector as previously described in (Onkokesung *et al*. 2011). Approximately 100 mg of frozen leaf tissue powder were extracted with 1 mL of acidified (0.5% acetic acid) 40% methanol. Small metal balls were added to the samples and those were then homogenized with a ball mill (Genogrinder 2000; SPEX CertiPrep) for 45 s at 250 strokes/min. Homogenized samples were centrifuged at 16,000 g, 4°C for 30 min. 400 μL of each supernatant were transferred to 2-mL glass vials for analysis on an Agilent HPLC 1100 series device (http://www.chem.agilent.com). 1 μL of each sample was injected into a Chromolith FastGradient RP 18-e column (50 x 3 x 2 mm; monolithic silica with bimodal pore structure, macropores with 1.6mm diameter; Merck) protected by a precolumn (Gemini NX RP18, 2 x 3 x 4.6 mm, 3 mm). The elution gradient was based on a binary solvent system. Solvent A consisted in 0.1% formic acid and 0.1% ammonium water, pH 3.5. Solvent B was pure methanol. The following conditions were applied for the gradient [time/concentration (min/%) for B]: 0.0 min/0%, 0.5 min/0%, 6.5 min/80%, 9.5 min/80%, and reconditioning for 5 min to 0% B. The flow rate was constant at 0.8 mL/min and the column temperature was set to 40°C. Phenolamides were detected by operating the PDA at a wavelength of 320 nm. Quantification was performed by building external calibration curves for authentic standards of caffeoylputrescine and chlorogenic acid (CGA). *N’,N*”-dicaffeoylspermidine (DCS) isomeric peaks were pooled and reported as total DCS. Since no standard for DCS was available, its content was expressed as CGA equivalents.

### Untargeted UHPLC-TOF-MS measurements

To achieve a comprehensive view on the secondary metabolite profile of the leaves of the different plant genotypes, an untargeted approach was undertaken by running extracted leaf samples on an Ultra High Pressure Liquid Chromatography (UHPLC) coupled with a Time-of-Flight (TOF) Mass Spectrometer (MS) as described in (Onkokesung *et al*. 2011). Two microliters of leaf extracts prepared as above were separated using a Dionex Rapid Separation LC system equipped with a Dionex Acclaim 2.2-mm 120A, 2.1 x 150 mm column. The elution gradient was based on a binary solvent. Solvent A consisted in deionized water with 0.1% acetonitrile and 0.05% formic acid. Solvent B consisted in acetonitrile with 0.05% formic acid. The following conditions were applied for the gradient: 0 to 5 min, isocratic 95% A, 5% B; 5 to 20 min, linear gradient to 32% B; 20 to 22 min, linear gradient to 80% B; isocratic for 6 min. The flow rate was constant at 0.300 mL/min. Eluted compounds were detected by a MicroToF mass spectrometer (Bruker Daltonics) equipped with an ESI source operated in positive ionization mode. Typical instrument settings were as follows: capillary voltage, 4,500 V; capillary exit, 130 V; dry gas temperature, 200°C; dry gas flow of 8 L/min. Ions were detected from *m/z* 200 to 1,400 at a repetition rate of 1 Hz. Mass calibration was performed using sodium formate clusters (10 mM solution of NaOH in 50%:50% isopropanol:water containing 0.2% formic acid).

Raw chromatograms were exported to netCDF format using the export function of the Data Analysis version 4.0 software (Bruker Daltonics) and processed using the XCMS package in R (Smith, Want, O’Maille, Abagyan & Siuzdak 2006). Peak detection was performed using the centwave algorithm with the following parameter settings: ppm = 20, snthresh = 10, peakwidth = c(5,18). Retention time correction was accomplished using the XCMS retcor function with the following parameter settings: mzwid = 0.01, minfrac = 0.5, bw = 3. Areas of missing features were estimated using the fillPeaks method. Annotation of compound spectra derived from insource fragmentation during ionization and corresponding ion species was performed with the BioConductor package CAMERA (Kuhl, Tautenhahn, Böttcher, Larson & Neumann 2012). Compound spectra were built with CAMERA according to the retention time similarity, the presence of detected isotopic patterns, and the strength of the correlation values among extracted ion currents of coeluting *m/z* features. CAMERA grouping and correlation methods were used with default parameters. Clustered features were annotated based on the match (± 5 ppm) of calculated *m/z* differences versus an ion species and neutral loss transitions rule set. The R script used for XCMS and CAMERA post-processing of the data is provided as Supplemental File S1.

### Statistical Analysis

For the targeted quantitative analysis of polyamine, phenolamide, phytohormone and transcript levels, results were analyzed by univariate statistics using native functions in the R environment. Relevant statistical tests applied to the above data-sets are mentioned in figures captions. If normality assumption, or homoscedasticity, was not met, non-parametrical tests, respectively log2 transformation, were applied. Corresponding statistical results are provided in Supplemental File S4.

In the case of MS metabolomics data, a combination of univariate and multivariate statistical approaches was employed for data exploration. The complete 75^th^ percentile-normalized XCMS/CAMERA matrix used as input for the below statistical analyses is available as Supplemental File S2. To diminish redundancy within the data matrix, multi-charged *m/z* features detected by the CAMERA analysis were excluded. *m/z* features consistently detected in at least 75% of the biological replicates of one factorial group were considered for further analysis. Zero values, which remained after application of the “filling in” function in XCMS, were replaced by one-half of the minimum positive value of the ion across all time points and conditions in the original data. Raw intensity values were 75^th^ percentile normalized before statistical analysis. Metabolite fragmentation patterns were annotated as described in Supplemental File S1.To analyze data structure, PCA analysis was performed using the prcomp package for R via the MetaboAnalyst interface (http://www.metaboanalyst.ca). During this analysis, the data matrix was first scaled using the Pareto scaling procedure and 40% of the *m/z* signals detected as constant based on the examination of their interquantile range, were removed as these variables are unlikely to be used in modeling main trends in the data. Additionally, univariate statistical analyses statistical were performed from the metabolomics data matrix to identify genotype-specific and W+OS-specific effects on individual *m/z* features. Briefly, one-way ANOVAs followed by Tukey’s Honestly Significant Difference (Tukey’s HSD) post-hoc tests were carried out on a single time-point basis, after log2-transformation of the original data. The results are visualized as part of the Venn diagrams of Figure 2c and Figure S2 summarizing overlapping and non-overlapping deregulated *m/z* features in each of the tested genotypes. Corresponding statistical results are reported in Supplemental File S3. Heat-maps in Figure 3 summarized statistical effects and fold-change regulation for individual *m/z* features as analyzed using *t*-tests. In Figure 3, color-coded log2-transformed fold-changes correspond to those derived from a genotype comparison for which a statistically significant effect (P < 0.05) was detected. In addition to pairwise counter-genotype statistical comparison, W+OS-inducibility of single *m/z* features in the EV background was also analyzed by statistically comparing peak areas in the W+OS treated EVs compared to their corresponding untreated controls. Finally, a non-parametric Wilcoxon rank test was used for pairwise comparison between each of the transgenic lines and the EV of the distribution of log2-transformed fold changes (W+OS *vs* untreated controls) obtained for features within the *m/z* cluster 1. Corresponding statistical results are reported in Supplemental File S5.

**Figure 3.**
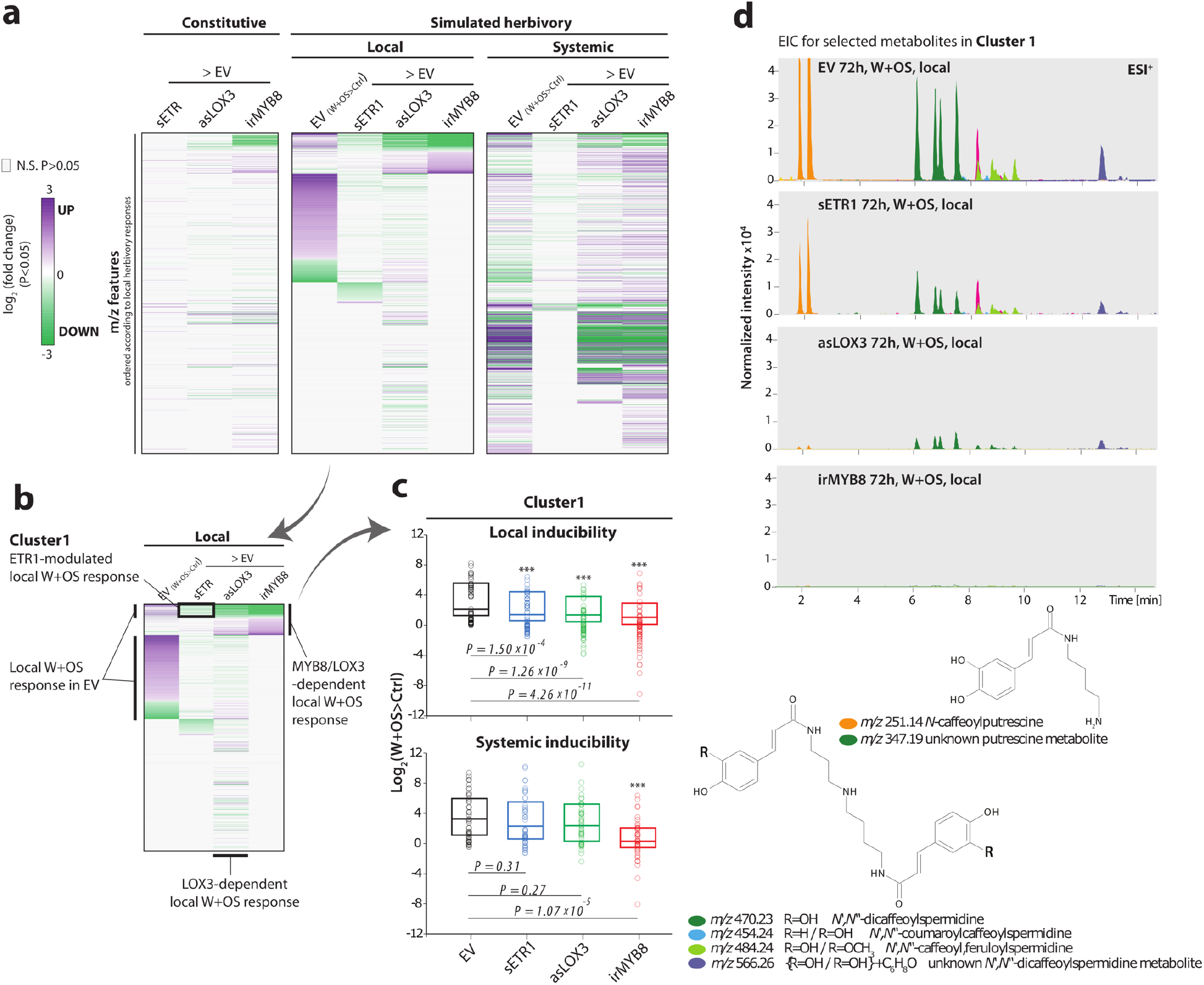
Statistical processing of MS-based metabolomics data pinpoints on a cluster of JA/MYB8-dependent phenolamides whose local W+OS induction is attenuated in sETR1. **(a)** Transgenic line-specific deregulations in metabolite signals (*m/z* features): color-coded cells (green to violet) in the heat-maps denote, unlike grey cells, features for which a statistically significant effect was detected for a given comparison. The first heat-map visualizes constitutive (untreated leaves) deregulation in the three reported transgenic lines compared to in the corresponding empty vector plants (EV). The next two heat-maps focus on local and systemic deregulations at 72 h after W+OS (simulated herbivory) in sETR1, asLOX3 and irMYB8 compared to the same conditions for the EV. For these two heat-maps the first column depicts *m/z* feature herbivory-regulation in the EV by comparing the level of intensities after W+OS of the concerned features compared to in controls. All pairwise comparisons involved *t*-tests (P = 0.05 as threshold for statistical significance). **(b)** Annotation of patterns visualized in the “local” heat-map after ordering of log2-transformed fold change values according to in the EV, next for the irMYB8 and finally for the sETR1 background. Cluster 1 comprises many *m/z* features induced locally in the EV but whose local induction is strongly impaired in irMYB8/asLOX3 and attenuated in sETR1. **(c)** Comparison of log2-transformed fold changes after W+OS only for features of cluster 1. Distributions were statistically compared to in EV by Wilcoxon rank sum tests **(d)** Uniformly-scaled representative extracted ion currents (electrospray positive, ESI+) in the different genotypes for *m/z* values corresponding to precursors ([M+H]^+^ adducts) of phenolamides enriched in cluster 1. Putrescine- and spermidine-based phenolamide annotations are provided.

## Results

### *Effect of genetic manipulations of ethylene and jasmonate signaling on simulated leaf herbivory in* N. attenuata

In the present study, we made use of previously generated transgenic lines that are specifically manipulated in the perception of ET (sETR1), the biosynthesis of jasmonates (asLOX3) and in the expression of MYB8 (irMYB8), a pivotal jasmonate-dependent transcription factor regulating herbivory-induced phenolic metabolism (Figure 1a). Preliminary to the exploration of metabolomics dynamics in the different transgenic lines and an empty vector (EV) control line, we extended on previous analyses of jasmonate bursts in the sETR1 line. In agreement with previous studies by von Dahl *et al*. and Onkokesung *et al*. on the ET perception-mediated feedback inhibition of ET biosynthesis in sETR1 (von Dahl *et al*. 2007; Onkokesung *et al*. 2010b), leaves of the sETR1 line used in the present study emitted twice more ET over a 10 h period than those of the EV (Figure 1b). Total ET emissions by asLOX3 and irMYB8 were unaltered compared to that in EV (Figure 1b). Peaking levels of JA were identical in locally-elicited leaves of sETR1 plants to those of EV, albeit a 28% reduction in the levels of JA-Ile was detected (Figure 1c, d). Jasmonate levels had not been previously characterized by Kaur *et al*. (Kaur *et al*. 2010) as part of their phenotyping study on irMYB8 transgenic plants. Here, while locally-induced peaking JA levels at 1 h in leaves of irMYB8 were on average greater than those of EV, no statistically significant differences were detected when comparing jasmonate dynamics in this line and the EV background. Salicylic acid (SA) and abscisic acid (ABA) were also measured in the context of these analyses. Levels of both hormones increased after the W+OS (Figure S1), but no alterations specific to the manipulation of ET signaling were detected.

### Transgenic lines exhibit overlapping and non-overlapping deregulations of herbivory-induced metabolic responses

To explore the control exerted by phytohormonal cross-talks over herbivory-induced metabolic adjustments, local and systemic leaves of the aforementioned transgenic lines were collected 0, 24 h, 48 h and 72 h following simulation of insect herbivory. They were analyzed by untargeted MS-based metabolomics using an optimized UHPLC-ESI^+^-qTOF-MS method that we previously optimized to profile polar-to-semi-polar *N. attenuata* leaf metabolites (Gaquerel, Heiling, Schoettner, Zurek & Baldwin 2010). The overall procedure makes use of the power of non-targeted MS metabolomics coupled with statistical analyses to infer overlapping and non-overlapping sets of differentially-regulated metabolites across transgenic manipulations of hormonal signaling pathways. A concatenated peak (defined as a mass-to-charge ratio feature, *m/z*, at a single retention time) matrix was exported from the UHPLC-ESI^+^-qTOF-MS factorial data-set. This matrix was then subjected to a principal component analysis (PCA). As expected, the resulting PCA score plot showed a clear demarcation between local and systemic leaf-derived metabolic profiles, as well as between metabolic profiles corresponding leaves before and after simulated herbivory (Figure 2b). In line with the central role of JA signaling in the activation of herbivory-induced metabolic changes, separation was not as strong in asLOX3 plants as in the other genotypes. *m/z* features exhibiting greatest loading values for sample clustering according to the two principal components (PC; Figure 2b) corresponded to nicotine ([M+H]^+^: C_10_H_15_N_2_^+^ *m/z:* 163.12, loading PC1: 0.44) and several phenolamides, such as CP ([M+H]^+^: C_13_H_19_N_2_O_3_^+^ *m/z*: 251.14, loading PC1: 0.33) and DCS ([M+H]^+^: C_25_H_32_N_3_O_6_^+^ *m/z*: 470.23, loading PC2: 0.15). To compare metabolic responses in the different test lines, the overlap of significantly deregulated *m/z* features in asLOX3, irMYB8 and sETR1 plants (compared to in EV) was investigated by means of Venn diagrams (Fig 2c and Figure S2). The latter analysis revealed that the identity of deregulated *m/z* features in asLOX3 and irMYB8 plants largely overlapped in the two leaf positions collected, while deregulated features in sETR1 plants only partly overlap for locally-elicited leaves with those inferred in asLOX3 and irMYB8 genetic backgrounds.

### ET-insensitivity differentially alters local and systemic metabolic responses to simulated herbivory

To further parse the most important metabolic deregulations activated in each of the different transgenic lines, we then conducted univariate statistical analyses using the *m/z* feature concatenated matrix and visualized statistical results as heat-maps (Figure 3). Briefly, the first heat-map of Figure 3 visualizes constitutive (untreated leaves) deregulation in the three reported transgenic lines compared to the corresponding empty vector plants (EV). The next two heat-maps focus on local and systemic deregulations at 72 h after W+OS treatment in sETR1, asLOX3 and irMYB8 compared to the same conditions for EV. We further ranked *m/z* features according to most important statistical changes in locally-induced EV in order to visualize how the corresponding core-set of metabolites was regulated in the different transgenic lines. This data exploration approach confirmed that, as expected, the majority of locally-induced *m/z* features in EV was significantly down-regulated in asLOX3 plants. Those down-regulations were even more pronounced in systemically-induced leaves. Similarly, the impact of silencing MYB8, as materialized by the down-regulation of many herbivory-induced *m/z* features, was most clearly apparent in systemically-induced leaves. From the examination of heat-maps for local metabolic responses, one cluster, hereafter referred to as cluster 1, raised our attention. Cluster 1 comprises herbivory-induced *m/z* features severely down-regulated in asLOX3 and irMYB8 and with less pronounced statistical effect of down-regulation in sETR1. Overall, the inducibility of cluster 1 was significantly reduced in in locally-induced leaves of the three transgenic lines, while a statistically significant reduction of cluster inducibility was only detected for the irMYB8 line when considering systemic leaf metabolic profiles (Figure 3b and c). Extracted ion chromatograms for some of the most abundant *m/z* features in cluster 1, which corresponded to protonated adducts for previously characterized MYB8-dependent phenolamides, confirmed down-regulations in the production of these compounds in leaves of sETR1 plants locally-elicited by the W+OS treatment.

Table 1 summarizes fold change deregulations in the intensity of *m/z* signals characteristics for those and other phenolamides previously structurally identified or annotated based on diagnostic fragments (Gaquerel *et al*. 2010; Onkokesung *et al*. 2011). As indicated in previous studies (Onkokesung *et al*. 2011), production of the complete detectable phenolamide chemotype was abolished in irMYB8 plants, while deficiency in JA signaling (asLOX3) led to a weaker suppression of these phenolic derivatives. In asLOX3 systemic leaves, several di-acylated spermidine isomers were up-regulated compared to in EV, suggesting that jasmonate-signaling may also more subtly contribute in balancing the production of the different spermidine-containing phenolamides. Within cluster 1, *N*-caffeoylputrescine, which was previously detected as one of the most strongly induced phenolamides in *N. attenuata* upon insect herbivory (Kaur *et al*. 2010; Onkokesung *et al*. 2011), appeared as the one most significantly one affected by the insensitivity to ET (Table 1, Figure 4, Figure 5). Consistent with a tuning effect of ET over the phenolamide chemotype rather than a complete pathway-level control, subtle remodelings of phenolamide levels were detected in locally-elicited sETR1 leaves. In particular, only a sub-part of the profile of spermidine-based phenolamides (*N*’,*N*” dicaffeoylspermidine and *N*’,*N*”-caffeoyl,feruloyspermidine) exhibited a significant reduction in herbivory-induction in sETR1 plants, albeit never to the extent of the severe down-regulation effects detected in irMYB8 and asLOX3 (Figure 4). We additionally conducted a smaller-scale metabolomics profiling of a previously characterized irACO transgenic line which is silenced for the ACC OXIDASE and hence biosynthetically repressed for ET production (Figure S3). In contrast with the sETR1 line, which did not exhibit major vegetative growth defects, the irACO line, under our experimental conditions, exhibited a delay in rosette growth compared with EV and stress symptoms (notably numerous necrotic spots detected onto leaves), which hampered its initial use in the larger metabolomics exploratory approach. Similarly to sETR1, locally-induced levels of several phenolamides were reduced in irACO compared with EV (Figure S3).

**Figure 4.**
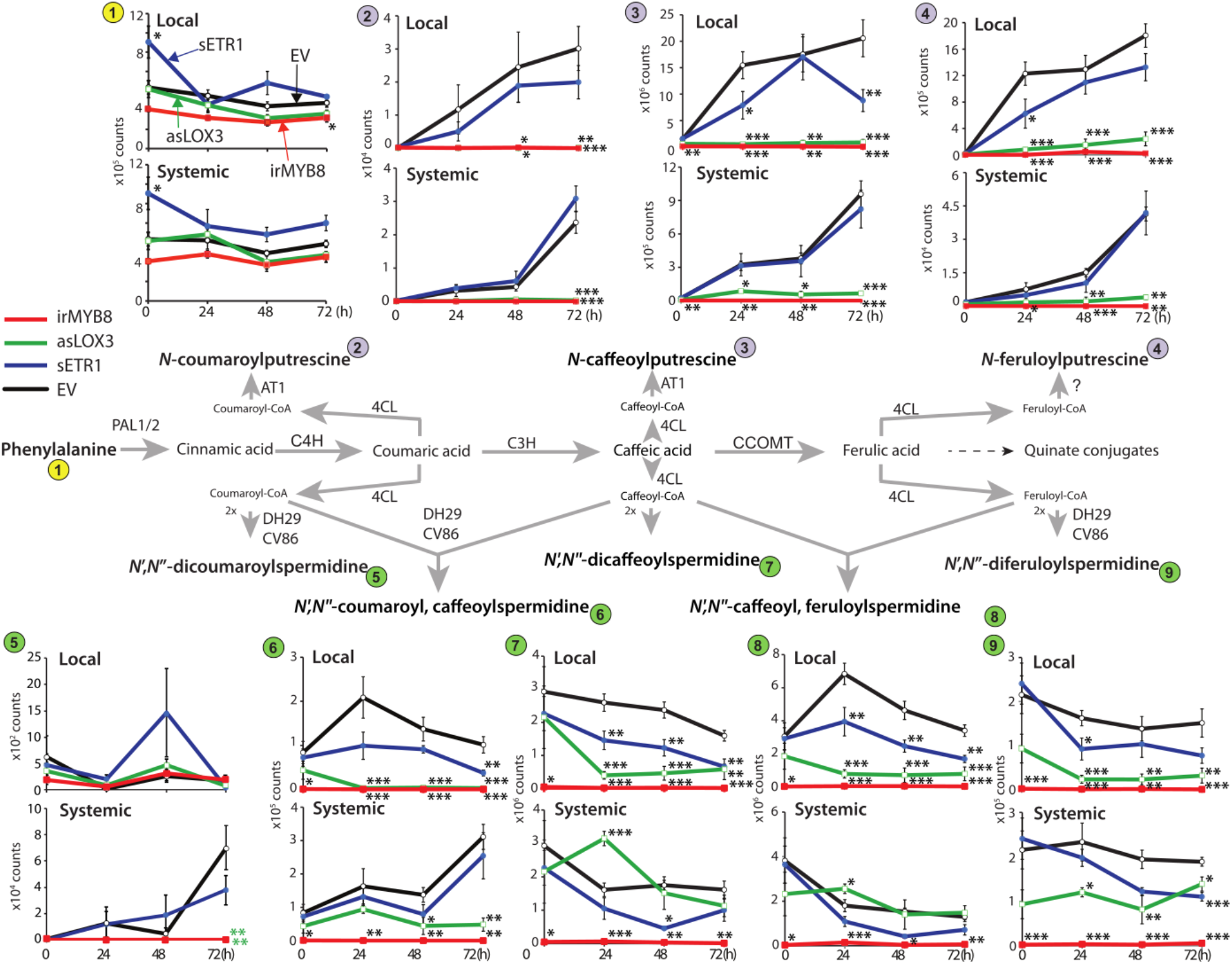
Local accumulation dynamics of JA/MYB8-dependent phenolamides in response to W+OS elicitation are modulated by ET in *Nicotiana attenuata*. Mean values (± SE, 4 to 5 biological replicates) of phenolamides and phenylalanine main *m/z* features in local and systemic leaves during simulated herbivory. Asterisks indicate significant differences with EV at respective time points (* P < 0.05, ** P < 0.01, *** P < 0.001, ANOVA followed by Tukey HSD post-hoc tests). Leaves from EV, sETR1, asLOX3 and irMYB8 plants were harvested at different time points before and after W+OS treatment. Their metabolites were extracted with acidified methanol and subjected to mass spectrometry analysis using UHPLC-qToF-MS operating in positive mode. Only *m/z* features of phenylalanine and known phenolamides were kept for further analysis. The biosynthetic pathway of phenolamides from phenylalanine and phenylpropanoids is depicted. Enzymes are indicated above and below reaction arrows.

**Figure 5.**
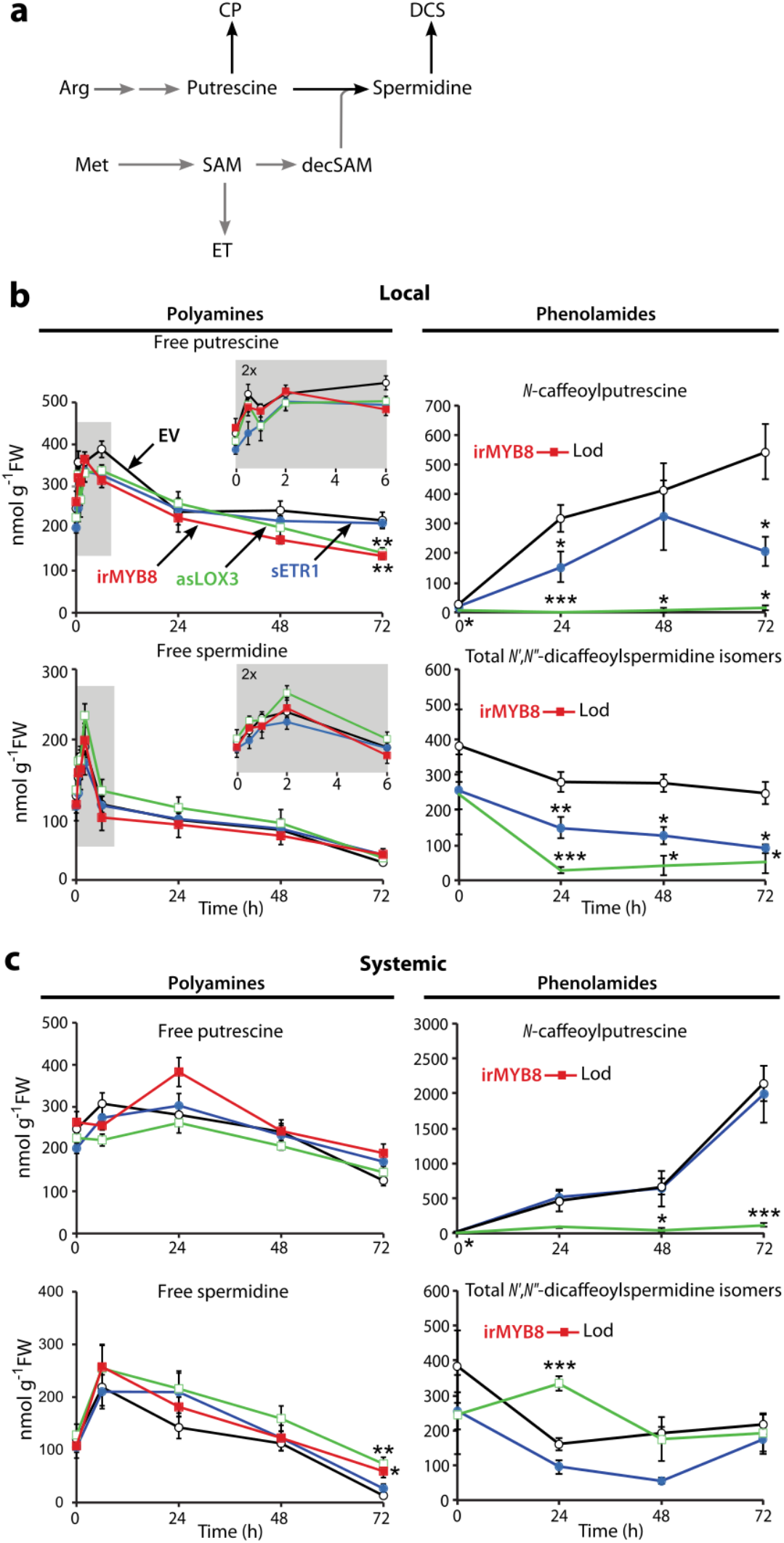
Targeted HPLC-UV analysis of free polyamines and phenolamides indicate that the modulatinethylene on phenolamide accumulation after herbivory, without any influence on polyamines pathway. **(a)** Schematic pathway depicting biochemical connections between ethylene and polyamine synthesis and acylation of polyamines into *N. attenuata* main phenolamides (in black). Arg, arginine. CP, caffeoylputrescine. DCS, dicaffeoylspermidine. decSAM, decarboxylated-*S*-adenosylmethionine. ET, ethylene. Met, methionine. SAM, S-adenosylmethionine. **(b)** and **(c)** Mean values (± SE, 5 biological replicates) of free polyamines and phenolamides in local and systemic leaves, respectively, during simulated herbivory. Leaves from EV, sETR1, asLOX3 and irMYB8 plants were harvested at different time points before and after W+OS treatment. “Systemic” refers to undamaged young leaves. Free polyamines levels were measured by HPLC-UV in leaves extracts subjected to derivatization and quantified with appropriate standards. Insets show details on the dynamic induction of free polyamines at early time points of the same experiment. Phenolamides levels were measured from leaves extracts by HPLC-PDA and quantified with appropriate standards when available. DCS are thus reported as CGA equivalent. DCS levels are a pool of the different DCS isomers that were detectable. Phenolamides were undetectable in irMYB8 plants (Lod). ANOVA tests followed by Tukey HSD post-hoc tests were applied to free polyamines and phenolamides levels at each time point. When normality assumption was not met, Kruskal-Wallis and pairwise Wilcoxon rank sum tests were applied. Only significant differences between EV plants and sETR1, asLOX3 or irMYB8 plants are reported. * P < 0.05, ** P < 0.01, *** P < 0.001.

**Table 1.**
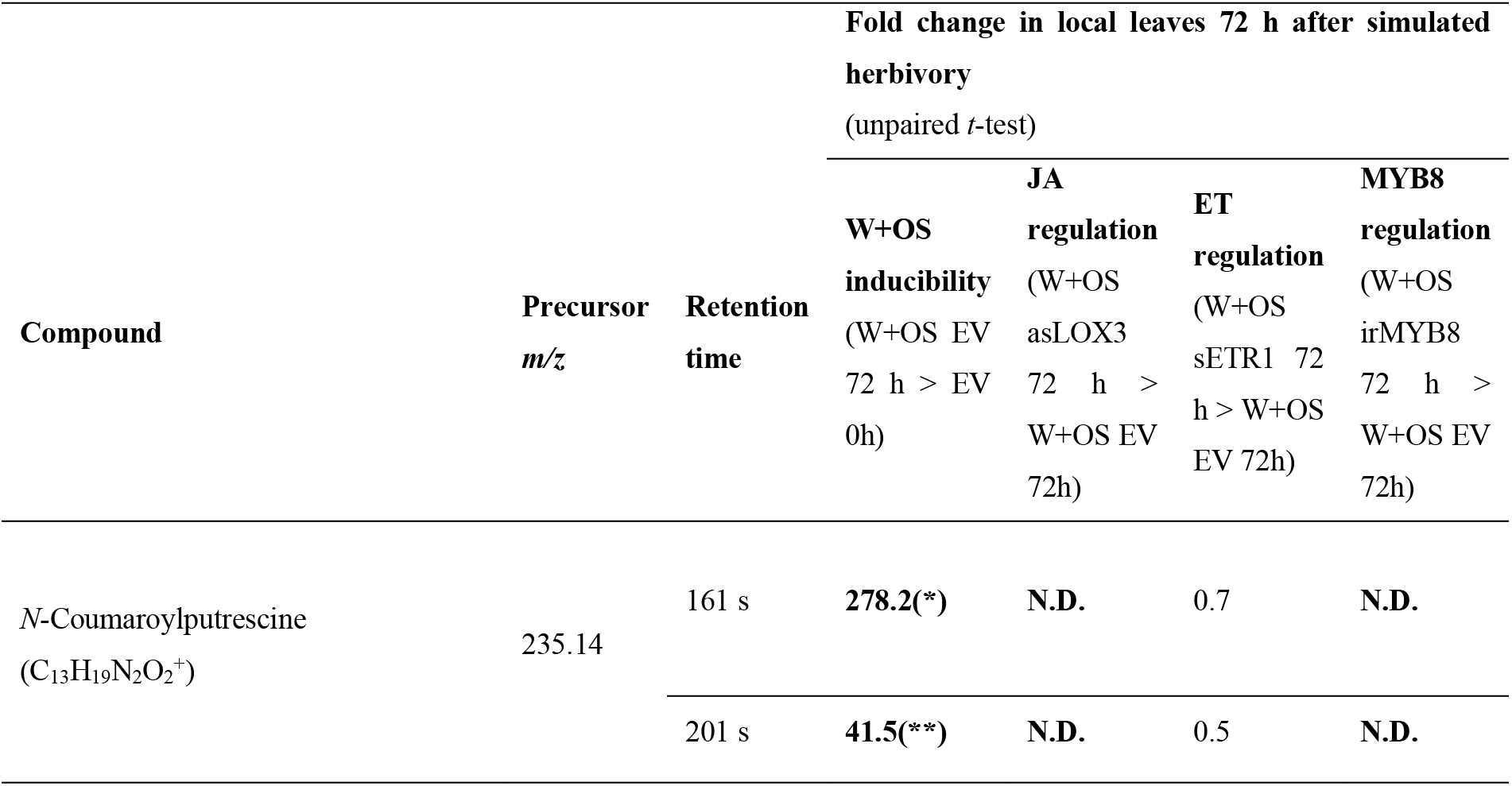
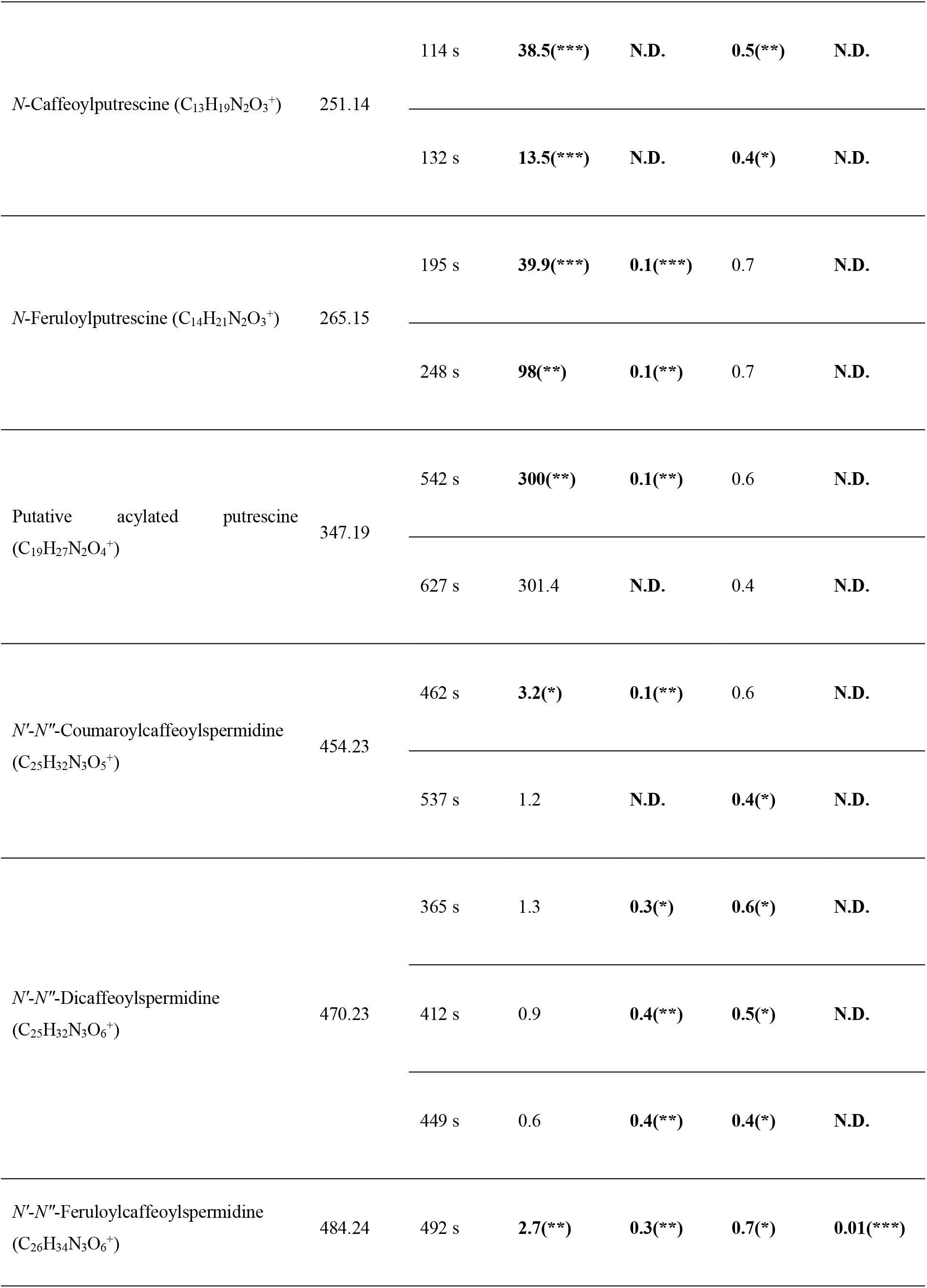
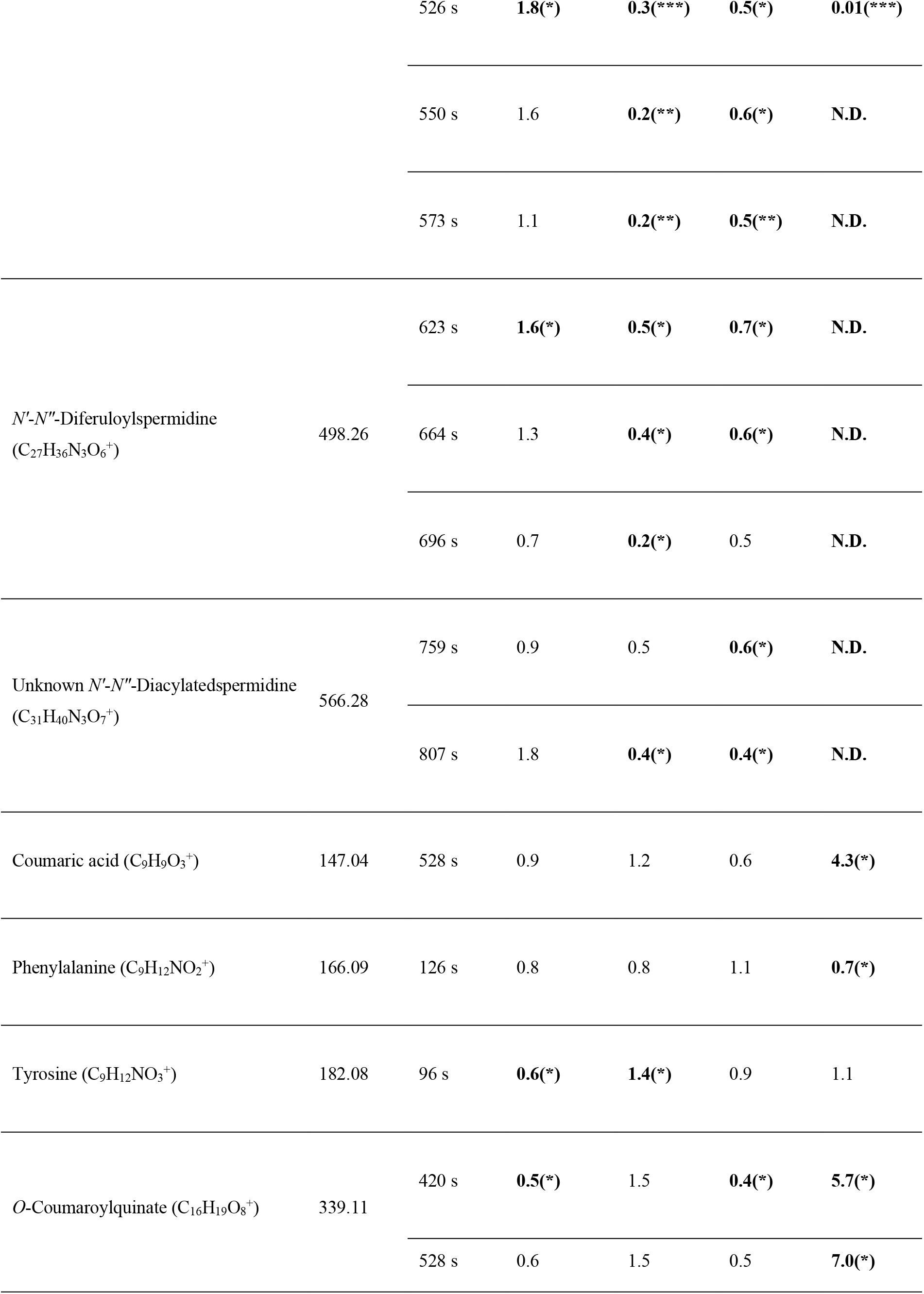
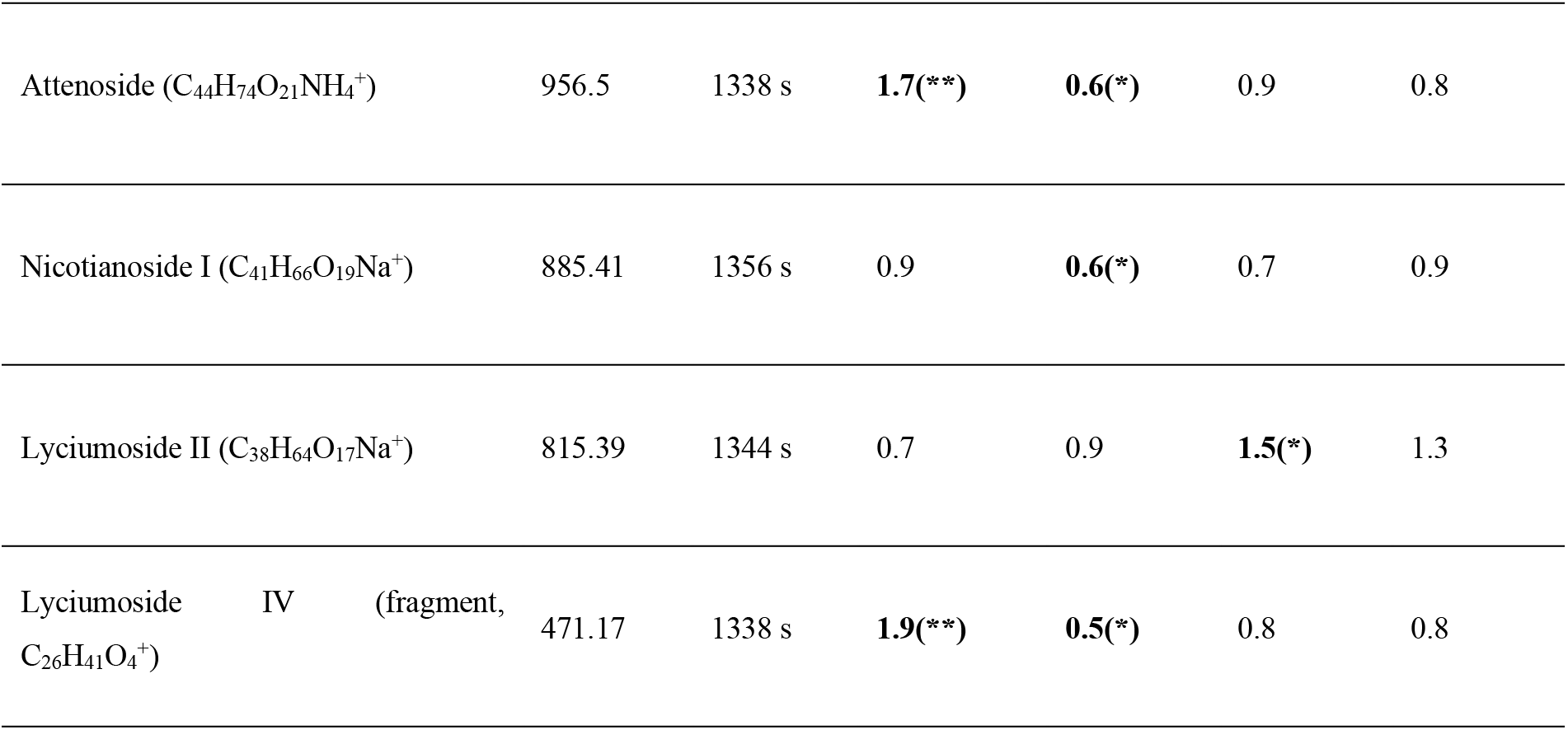
Known metabolites measured by UHPLC-TOF-MS in local leaves after simulated herbivory, which are differentially regulated by the jamonate, ethylene and MYB8 pathways. Rosette leaves of wild-type, asLOX3, sETR1 and irMYB8 plants were wounded and treated with diluted oral secretion of caterpillars (W+OS treatment). Leaves before and 72 hours after the treatment were harvested, and their metabolites were extracted with acidified methanol. Extracts were subjected to mass spectrometry analysis using UHPLC-qToF-MS. *m/z* features were recorded for different elution times and analysed using Metaboanalyst. According to previous studies, some *m/z* features could have been assigned to known compounds (Gaquerel *et al*., 2010, Onkokesung *et al*., 2011). Only known compounds that were differentially regulated in at least one genotype are presented. Fold change were calculated either by comparing treated leaves with control leaves in wild-type (local inducibility) or by comparing treated leaves in the different genotypes with treated leaves in wild-type. Unpaired *t*-tests were calculated from Metaboanalyst-analysed data and sample weights were taking into account to normalize peak area. * P < 0.05, ** P < 0.01. *** P < 0.001. Significant up- and down-regulations are highlighted in grey and in bold font. These regulations are due to W+OS elicitation (local inducibility) or RNAi silencing (asLOX3, sETR1 or irMYB8). N.D., Not Detected.

Previously, we evidenced that the metabolic tension in coumarate allocation between polyamine- and quinate-hydroxycinnamate conjugates is exacerbated during herbivory (Gaquerel, Kotkar, Onkokesung, Galis & Baldwin 2013). In the present study, free coumarate and its derivative *O*-coumaroylquinate over-accumulated in local and systemic leaves of irMYB8 plants during simulated herbivory (Figure S4). The latter result indicates that irMYB8 plants, which barely produced phenolamides, likely channeled coumarate units not conjugated to polyamines toward the production of quinate derivatives.

### ET perception is required for intact local herbivory-induced levels of the two most abundant phenolamides

Next, we conducted targeted quantitative analyses for the two most abundant phenolamides, namely CP and DCS. Their relative levels appeared, in our metabolomics data-set, to be modulated locally by the ET signaling sector (Figure 5). Targeted analyses are instrumental to confirm trends detected from the untargeted metabolomics approach. As we previously reported in the untargeted approach, CP levels dramatically increased in response to W+OS treatment. This effect was stronger in undamaged (systemic) leaves than in local leaves: CP accumulated 4 times more in systemic leaves at 72 h. Our quantitative analyses confirmed that CP and DCS were down-regulated in asLOX3 and sETR1 plants compared to EV plants (Figure 5). Levels of these two compounds in sETR1 plants were intermediary to those detected at corresponding sampling times in EV and asLOX3 plants. Impairing ET signaling in the sETR1 line significantly reduced the production of CP and DCS only in local leaves (by 62% and 64% at 72 h, respectively; Figure 5b), but not in systemic leaves (Figure 5c). Finally, in contrast with previous studies (Kaur *et al*. 2010; Onkokesung *et al*. 2011), we did not detect in our analyses an induction of total DCS levels in response to the W+OS treatment (Figure 5b, c). Total DCS levels were nearly constant, or slightly decreased, during the time-course experiment in response to the elicitation.

### ET signaling does not affect levels of free polyamines during simulated Herbivory

We then explored whether local herbivory-induced ET production metabolically imparts on polyamine homeostasis for phenolamide production because of the shared SAM precursor (Figure 1a). To this end, we quantified the time-course changes in levels of free putrescine, spermidine and spermine in leaf tissues after simulated herbivory. The extent to which the pools of theses amines are reconfigured during insect herbivory had remained, to our knowledge, largely unexplored. Constitutive amine levels were not statistically different among transgenic lines and the EV (Figure 5b, c). Upon W+OS treatment, spermidine levels decreased consistently in all lines in both local and systemic tissues, resulting in up to 10 times less free spermidine detected in EV plants at 72 h compared to prior treatment (Figure 5b). While strongly decreased because of the W+OS treatment, levels of spermidine in systemic leaves of asLOX3 and irMYB8 collected at 72 h were significantly higher than those in EV. In locally-treated leaves, spermidine and putrescine exhibited transient elevations, albeit weaker in the case of putrescine, consecutive to the simulated herbivory (Figure 5b). Putrescine levels measured at 72 h in local leaves were significantly reduced by 60% compared to those in EV. Importantly, putrescine and spermidine levels in sETR1 did not differ from those in EV (Figure 5).

### Free tyramine levels strongly increase in locally-elicited leaves

We additionally quantified tyramine, an arylamine that can be acylated with hydroxycinnamic units to form an additional group of herbivory-induced phenolamides, not detected in our metabolomics screen (Bassard, Ullmann, Bernier & Werck-Reichhart 2010; Kim, Yon, Gaquerel, Gulati & Baldwin 2011). Of all amine analysed in this study, tyramine exhibited the strongest response to the simulated herbivory treatment with locally-induced levels, reaching up to 15 times those prior elicitation in EV plants (Figure S5). By clear contrast, asLOX3 and MYB8 plants, but not sETR1 plants, accumulated less tyramine in local leaves 72 h after simulated herbivory (Figure S5). Intrigued by the strong local accumulation of tyramine and its possible direct role as a defense compound *per se*, we conducted *M. sexta* larval performance assays on artificial diet supplemented with different doses of tyramine. No significant effect of tyramine on *M. sexta* weight gain was detected (Figure S5).

### Silencing ethylene perception does not significantly alter the transcription of MYB8, PAL2, AT1, DH29 and CV86

Since the effect of ET on the jasmonate-/MYB8-dependent induction of phenolamide production did not appear to result from modulations in the homeostasis of free polyamines, we finally measured the transcriptional regulation of genes involved in the biosynthesis of phenolamides. As expected, transcript levels of *MYB8, PAL2, AT1, DH29* and *CV86* strongly increased after simulated herbivory compared with corresponding controls (Figure 6). As previously shown (Gális *et al*. 2006; Kaur *et al*. 2010; Onkokesung *et al*. 2011), expression of the selected genes was compromised in asLOX3 and irMYB8. Only marginally significant reductions in transcript levels of *MYB8* and *CV86* were detected in sETR1 compared to EV (Wilcoxon rank sum tests; P = 0.095 and 0.063, respectively).

**Figure 6.**
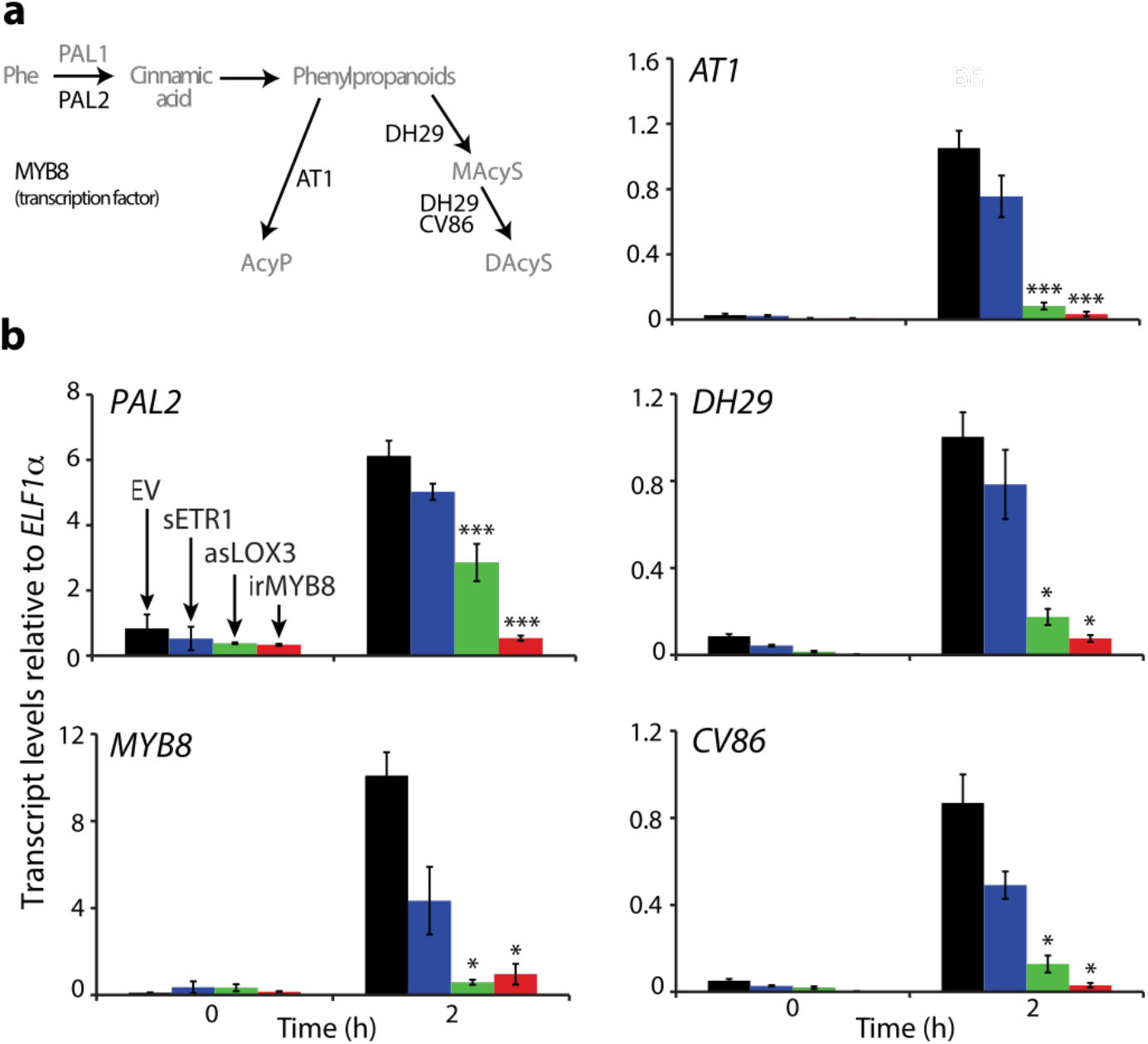
Ethylene insensitivity in sETR1 marginally affects the transcription of phenylpropanoid-phenolamide-related genes. **(a)** Involvement of four enzymes and one transcription factor in phenolamides production: phenylalanine ammonia lyase (PAL), MYB8 (transcription factor), AT1, DH29 and CV86. Enzymes whose gene transcriptions have been measured are in black font. Phe, phenylalanine. AcyP, acylated putrescine. MAcyS, mono-acylated spermidine. DAcyS, di-acylated spermidine. **(b)** Mean values (± SE, 4 to 5 biological replicates) of relative transcript of *PAL2, MYB8, AT1, DH29* and *CV86* in control (time point 0) and local leaves 2 h after simulated herbivory. * P < 0.05, *** P < 0.001.

## Discussion

Multiple phytohormone signaling sectors are activated in damaged tissues when plants are engaged into interactions with phytophagous. However, the mechanistic basis of this interplay between these phytohormones and the breath of defense traits under its control are not fully understood (Song, Qi, Wasternack & Xie 2014; Broekgaarden, Caarls, Vos, Pieterse & Van Wees 2015). In the present study, we combined exploratory MS-based metabolomics and analytical approaches targeted to phytohormones, phenolamides, polyamines, as well as relative quantifications of specific transcripts to gain new insights on ET-, JA- and MYB8 pathways in the response to *M. sexta*. Altogether, our results demonstrate that ET acts as an accessory signal for local JA-dependent reconfigurations of the phenolamide chemotype, provide new insights on the coupling between polyamine and phenolamide metabolisms and reinforce a role for ET in adjusting nitrogen investments into defense molecules.

Phenolamides have been characterized as herbivory- and JA signaling-induced metabolites against herbivores in *N. attenuata* (Kaur *et al*. 2010) and in other Solanaceae (Tebayashi *et al*. 2007), as well as in rice (Alamgir *et al*. 2016) and in maize (Marti *et al*. 2013),. Virtually any signaling nodes influencing jasmonate pools is therefore likely to alter induced phenolamide levels. In the present study, our metabolomics data exploration strategy identified that intact ET signaling is necessary to sustain full phenolamide production in locally-elicited. The importance of ET in this response, as detected in the sETR1 deaf for ET, was further observed in a smaller scale-profiling approach in the irACO line, which is biosynthetically-repressed for ET production. Other studies have recently uncovered a role of ET signaling for phenolamide metabolism. For instance, in *Arabidopsis thaliana*, the transcription of phenolamide-related genes was found to be triggered to higher levels of expression when both the JA and the ET pathways were simultaneously activated (Li *et al*. 2018). In rice, JA and ET signaling are both required for the elevated production of phenolamides in response to the infestation of the white-backed planthopper *Sogatella furcifera* (Wang *et al*. 2020).

In a recent large-scale metabolomics study, the degree of plasticity and diversity of the specialized metabolic profile of different *Nicotiana* species was investigated in response to herbivory by a “specialist” (*M. sexta*) and a “generalist” (*Spodoptera littoralis*, the cotton leaf worm) chewing insects (Li *et al*. 2020). It was identified that, while JA is the central signal shaping the amplitude of convergent metabolic responses to both herbivories, ET fine-tunes the specificity of this metabolic response according to the herbivore diet breadth. The latter study, as in the present one, detected that phenolamides are the foliar metabolites whose production is most tightly dependent on the fine-tuning signaling function of ET (Li *et al*. 2020). This convergence further reinforces a role of ET in mitigating the production of these metabolites according to insect herbivory type and echoes ecological implications suggested for the well-known function of ET in antagonizing the JA-dependent accumulation of nicotine when tobacco plants are attacked by nicotine-tolerant *M. sexta* larvae. It would be interesting to test whether deregulations in phenolamide levels in sETR1 compared to EV are also attenuated during herbivory by *S. littoralis* or *S. exigua*, whose OS do not elicit ET in *N. attenuata* compared to *M. sexta* (Diezel, von Dahl, Gaquerel & Baldwin 2009; Li *et al*. 2020). Finally, our study confirms that ET oppositely affects JA-regulated nicotine and phenolamide levels.

The fact that deregulations in phenolamide levels in sETR1 were only observed in locally-elicited leaves suggests a convergence of JA- and ET-dependent signaling in damaged cells. In this respect, previous work has shown that the interplay of JA and ET signaling at sites of mechanical damage during a W+OS treatment in *N. attenuata* mitigates cell division and tissue regrowth (Onkokesung *et al*. 2010b), but the regulatory bases to this response has remained elusive. Other phytohormonal cross-talks are known to affect induced phenolamide levels such as that between the JA and cytokinin pathways (Schäfer, Meza-Canales, Brütting, Baldwin & Meldau 2015). The ET pathway could act on several levels for the JA-dependent production of phenolamides. A possible layer of signaling regulation would be associated with an effect of herbivory-elicited ET on jasmonate levels. However, it is very unlikely that the moderate decrease of JA-Ile levels detected in this study solely account for the pattern of phenolamide accumulation in sETR1 plants. Additionally, deregulations in OS-elicited metabolic profiles extracted from sETR1 and asLOX3 locally-elicited leaves relatively poorly overlapped. Contrastingly, the strong overlapping between MYB8- and JA-dependent metabolic regulation supports the idea that MYB8 transcription factor activity more broadly impact defense regulation than only phenolamide accumulation (Schäfer *et al*. 2017).

Biochemical bases to the effect of ET on phenolamide levels were also examined. Indeed, a large body of reports on the parallel occurrence of ET and polyamine biosynthesis in certain tissues has long questioned whether SAM utilized for both metabolisms is rate limiting for either pathway (Bassard *et al*. 2010; Wuddineh, Minocha & Minocha 2018). This hypothesis of a metabolic tension between these two pathways has been tested in the context of several developmental responses including seedling development and fruit ripening. For the later development context, flux experiments on tomato fruits concluded on the fact that ET and polyamine can be synthesized simultaneously without exacerbated competition for SAM (Lasanajak *et al*. 2014). Such flux studies have been more rarely undertaken in the context of plant responses to biotic stresses. Transgenic tomato lines overexpressing a yeast spermidine synthase involved in polyamine (PA) biosynthesis resulting in a down-regulation of ET biosynthesis and signaling were identified as more susceptible to infection with *Botrytis cinerea* than WT plants (Nambeesan *et al*. 2012). Such deregulation of ET signaling and a consecutive increase in susceptibility to the biotic stress inflicted was not observed when the latter tomato lines were challenged by *M. sexta* herbivory (Nambeesan *et al*. 2012). The latter results suggest that genetically-engineering a strong increase in the polyamine biosynthetic flux does not alter the likely key signaling function of the ET signaling sector for defense induction against *M. sexta* in the above tomato lines.

In our study, levels of free polyamines in leaf tissues prior and in the first 72 h after simulated herbivory did not differ between EV and sETR1 plants, though the amplitude of the ET burst was twice more important in the latter line. Our data further indicate that the turnover of polyamines likely exceeds the production rate of the two key phenolamides measured, CP and DCS. Thus, the greater production of ET in sETR1 plants may not be sufficient to drain the pool of SAM engaged into polyamine biosynthesis. These results are hence in line with the above mentioned study which did not detect alterations in ^14^C incorporation from SAM to ET in transgenic lines producing high levels of spermidine during tomato fruit ripening (Lasanajak *et al*. 2014). Finally, free putrescine levels, whose production does not depend on SAM, did not accumulate in our study to higher levels in local leaves after simulated herbivory, as would normally be expected if less putrescine molecules were converted into spermidines from a diversion of SAM for ET synthesis.

Tyramine exhibited, from all analyzed amines, the most pronounced response to the simulated herbivory treatment. While earlier metabolomics studies conjointly detected herbivory-induced tyramine-containing phenolamides in leaves in sand-grown *N. attenuata* (Kim *et al*. 2011), the present and a recent metabolomics analyses (Gaquerel *et al*. 2010; Li *et al*. 2020) did not detect the presence of such metabolites in herbivory-induced profiles. It is unclear whether this metabolic shift for tyramine-containing phenolamides is related to the differential nitrogen supplies between these two soils and allocation in the plant (Lou & Baldwin 2004). We observed that putrescine and spermidine exhibit transient local responses to simulated herbivory, which precedes the biosynthesis of most abundant phenolamides such as CP and DCS. Compared with the tight regulatory control of JA- and MYB8-dependent signaling pathways over phenolamide production, which translate into massive alterations of the production of these compounds in the corresponding transgenic lines, levels of putrescine and spermidine levels were much less altered in asLOX3 and irMYB8. Such observation reinforces the idea that the control exerted by these signaling pathways for *de novo* phenolamides mostly occur at the level of the phenylpropanoid pathway-dependent hydroxycinnamate flux and its branching onto polyamine metabolism.

The fact that higher constitutive phenylalanine levels were detected in sETR1 plants but not in other transgenic lines points towards a possible effect of the ET on the phenylpropanoid pathway. Additional metabolic analyses would however be needed to examine possible alterations in the phenylpropanoid flux in sETR1 plants. Such a regulatory function of ET has already been suggested for the metabolism of fruits and vegetables. For instance, the ET pathway enhanced transcriptional levels of *PAL*, as well as the activity of the corresponding enzyme, during wound responses (Heredia & Cisneros-Zevallos 2009), ripening (Villarreal, Bustamante, Civello & Marti-nez 2010) and defense responses against the fungal pathogen *Rhizopus nigricans* (Pan, Fu, Zhu, Lu & Luo 2013). In our study, transcript levels of *PAL2* in sETR1 did not allow to firmly conclude on an effect of ET over the associated metabolic branch. More generally, even though marginal reductions in elicited levels for these transcripts were observed in sETR1 compared to EV controls, those trends did not appear to be statistically significant. Of note, a previous integrative study combining transcriptomics and metabolomics analyses has shown that remodeling of BAHD gene expression and *de novo* phenolamide production are not strictly collinear (Woldemariam, Oh, Gaquerel, Baldwin & Galis 2013), suggesting that biochemical conversions among pre-existing and *de novo* produced phenolamides are important layers of regulation in the herbivory-elicited response.

In conclusion, this study sheds light on the local interplay of JA-ET signaling sectors for defensive phenolamide production. Interestingly, a recent genetic repression of the circadian clock evening loop component *TOC1* in *N. attenuata* led to increase herbivory-induced levels of nicotine concomitant with lower ET signaling and strong decreases in phenolamide investments (Valim *et al*. 2020). Future integrative studies combining metabolome and transcriptome analyses during different types of herbivory could hence provide a clearer picture on the role of ET as one of the key orchestrators of nitrogen allocation to defense metabolite production.

## Acknowledgments

We are grateful to Matthias Schoettner, Michael Reichelt, Michael Stitz and Sven Heiling for technical assistance. We thank Jérôme Casas, Heidi Dalton and Dapeng Li for their valuable comments on an early version of this manuscript. The work was supported by funding of the Max Planck Society. ENS de Lyon is thanked for its financial support to F. Figon. E. Gaquerel’s research in Strasbourg is financially supported by the CNRS and the University of Strasbourg.

## Supplemental material

### Supplemental tables

**Table S1.**
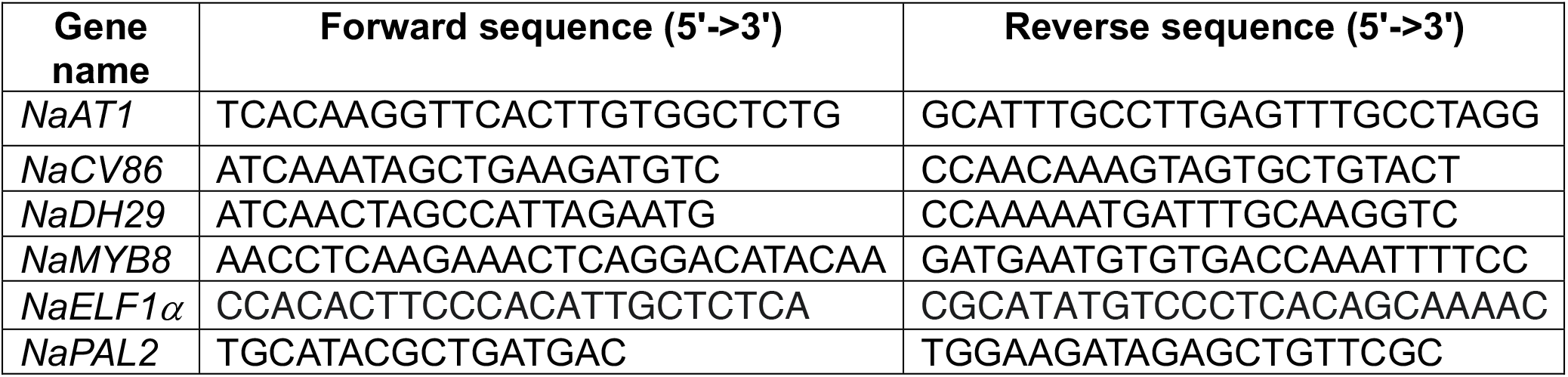
Primers used for qRT-PCR analysis.

### Supplemental figures

**Figure S1.**
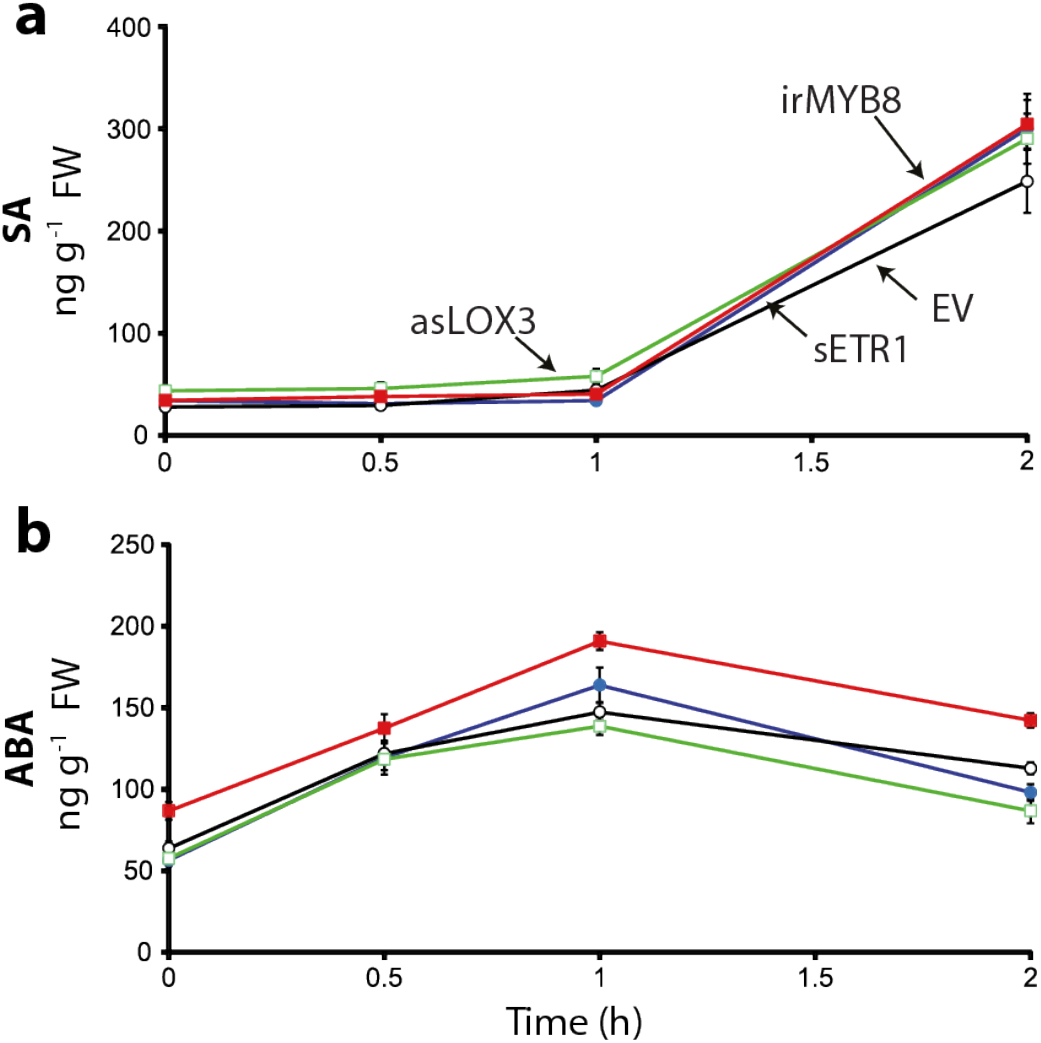
Accumulation dynamics of abscisic and salicylic acids after simulated herbivory are not affected by impairing either the jasmonate, MYB8 or ethylene pathways. Mean leaf levels (± SE, 5 biological replicates) of salicylic (SA, **a**) and abscisic (ABA, **b**) acids after simulated herbivory in wild-type (EV), sETR1, asLOX3 and irMYB8 transgenic plants. No significant differences in SA and ABA dynamics between EV and the other genotypes were observed.

**Figure S2.**
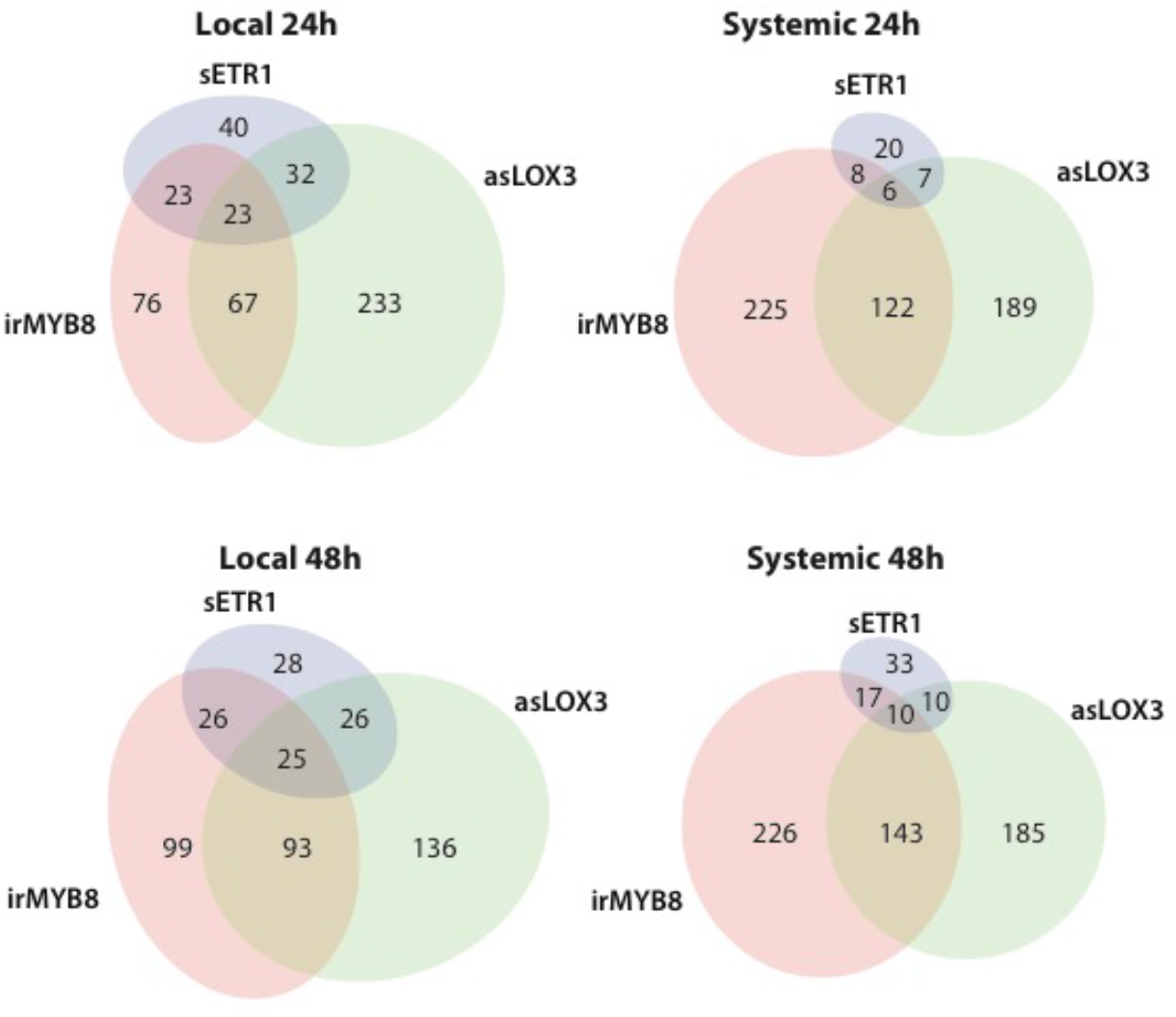
Venn diagram visualization of overlapping and non-overlapping significantly deregulated *m/z* features in each transgenic line backgrounds. Sampling/tissue-type-specific one-way ANOVA followed by Tukey HSD post-hoc tests (P ≤ 0.05) were conducted in sample measurements of local and systemic elicited leaves of each of the transgenic line compared to in the corresponding empty vector controls (EVs). Ellipse size and overlapping areas are respectively proportional to the total number of transgenic line-level deregulated *m/z* features and numbers of overlapping *m/z* features between transgenic lines.

**Figure S3.**
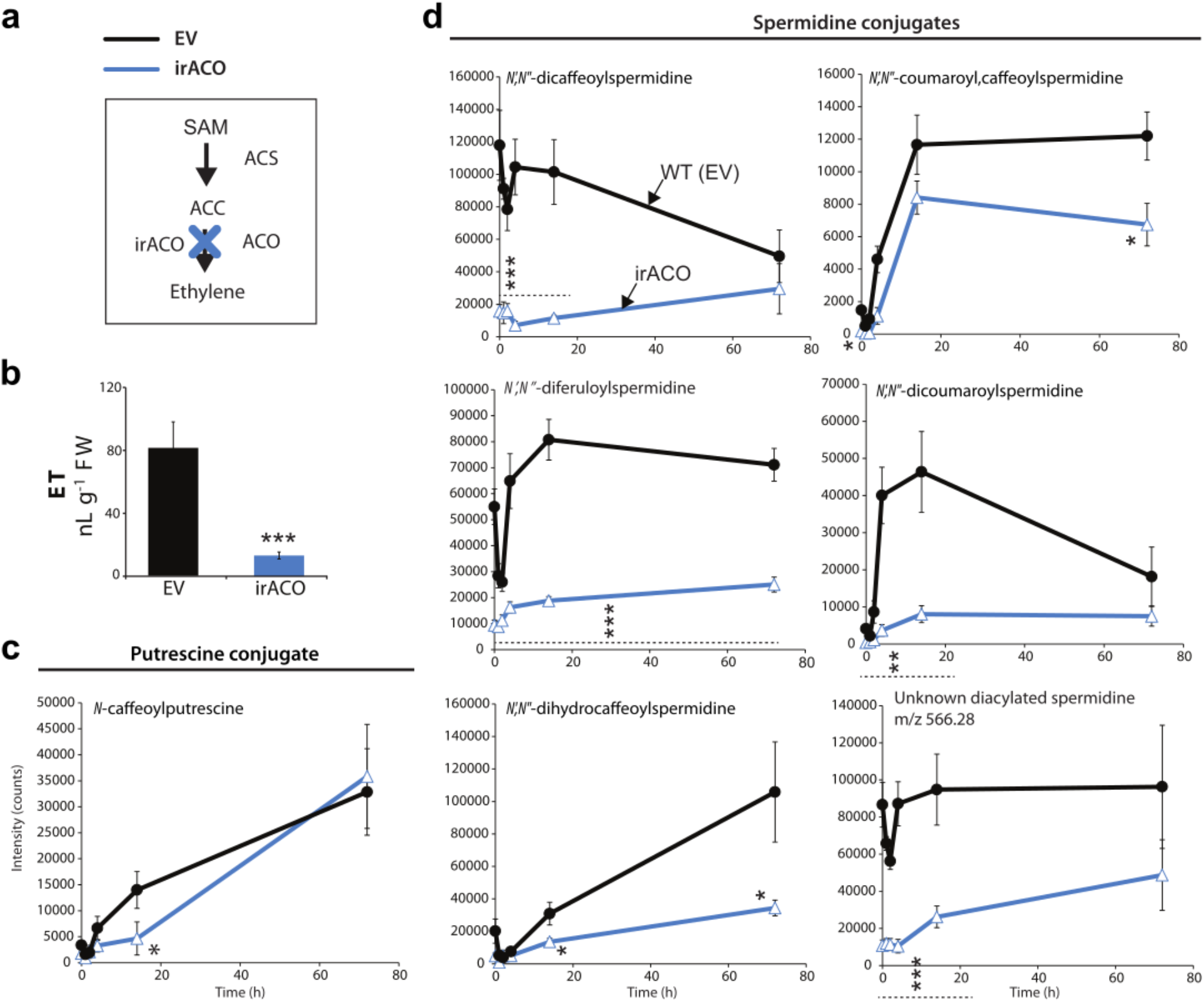
Impaired ethylene biosynthesis in the irACO transgenic line impairs constitutive and locally-induced leaf levels of phenolamides after simulated herbivory. **(a)** Biosynthesis pathway of ethylene (ET) from *S*-adenosylmethionine (SAM) catalyzed by the 1-aminocyclopropane-1-carboxylic acid (ACC) synthase (ACS) and oxidase (ACO). irACO transgenic plants are impaired in the production of ethylene from ACC. **(b)** Mean levels (± SE, 5 biological replicates, *** P < 0.001, Student *t*-test) of ET emission in EV and irACO transgenic leaf discs over 10 hours after simulated herbivory. **(c-d)** Normalized mean intensity (in counts, ± SE, 6 biological replicates) for *m/z* features corresponding to [M+H]^+^ adducts of dominant phenolamides detected in local and systemic leaves during simulated herbivory. Asterisks indicate significant differences with the empty vector (EV) at specific time points (* P < 0.05, ** P < 0.01, *** P < 0.001, ANOVA followed by Tukey post-hoc tests).

**Figure S4.**
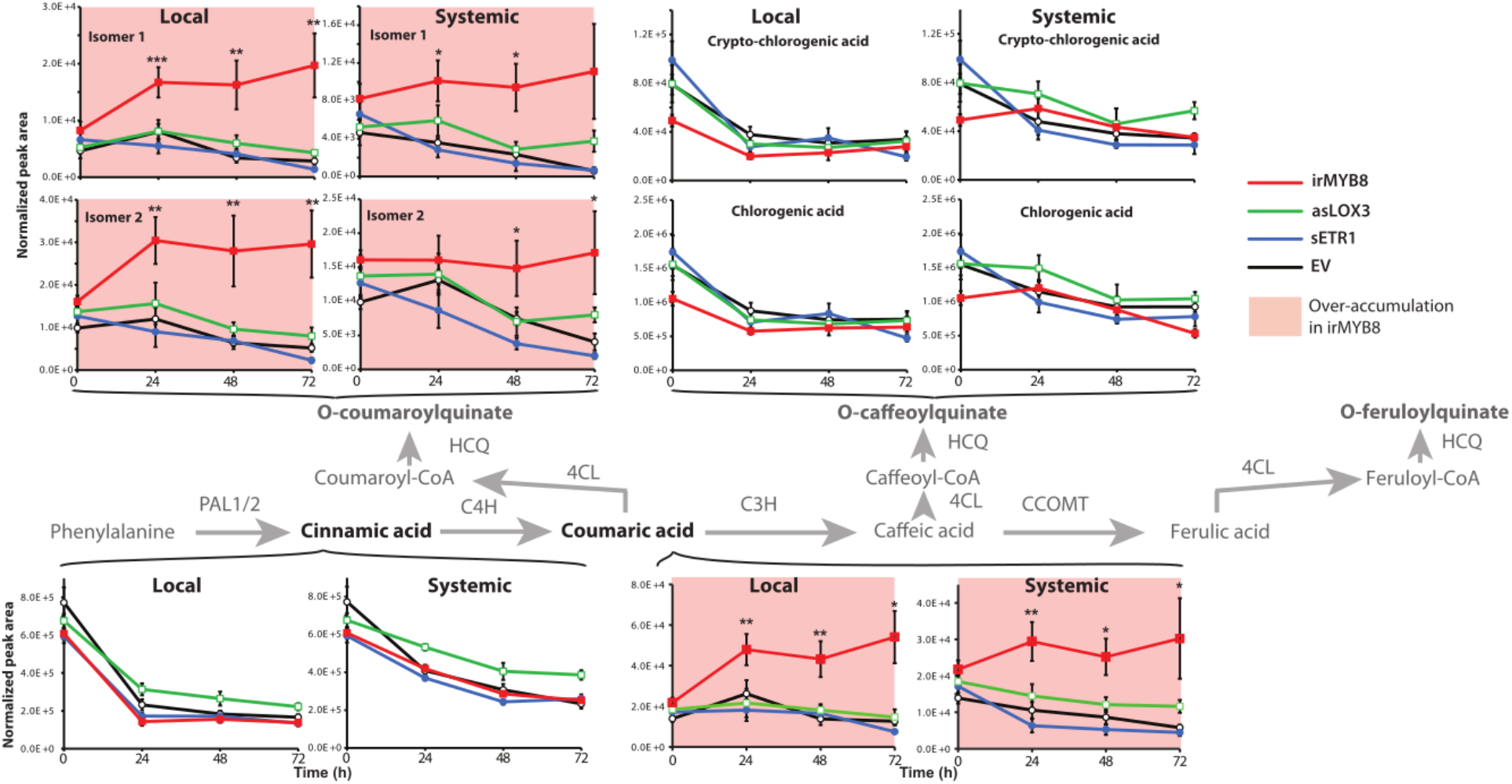
Silencing *MYB8* disrupts the incorporation of activated coumaric acid incorporation into phenolamide metabolism and creates an ectopic flux towards activated coumaric acid-quinate conjugation. Mean values (± SE, 4 to 5 biological replicates) of cinnamic acids and quinate conjugates main *m/z* features in local and systemic leaves during simulated herbivory. Asterisks indicate significant differences with EV at respective time points (* P < 0.05, ** P < 0.01, *** P < 0.001, ANOVA followed by Tukey HSD post-hoc tests). Leaves from EV, sETR1, asLOX3 and irMYB8 plants were harvested at different time points before and after W+OS treatment. Their metabolites were extracted with acidified methanol and subjected to mass spectrometry analysis using UHPLC-qToF-MS operating in positive mode. Only *m/z* features of phenylpropanoids and their quinate conjugates were kept for further analysis. “Systemic” refers to undamaged young leaves. The biosynthetic pathway of quinate conjugates from phenylalanine and phenylpropanoids is highlighted. Enzymes are indicated above and below reaction arrows.

**Figure S5.**
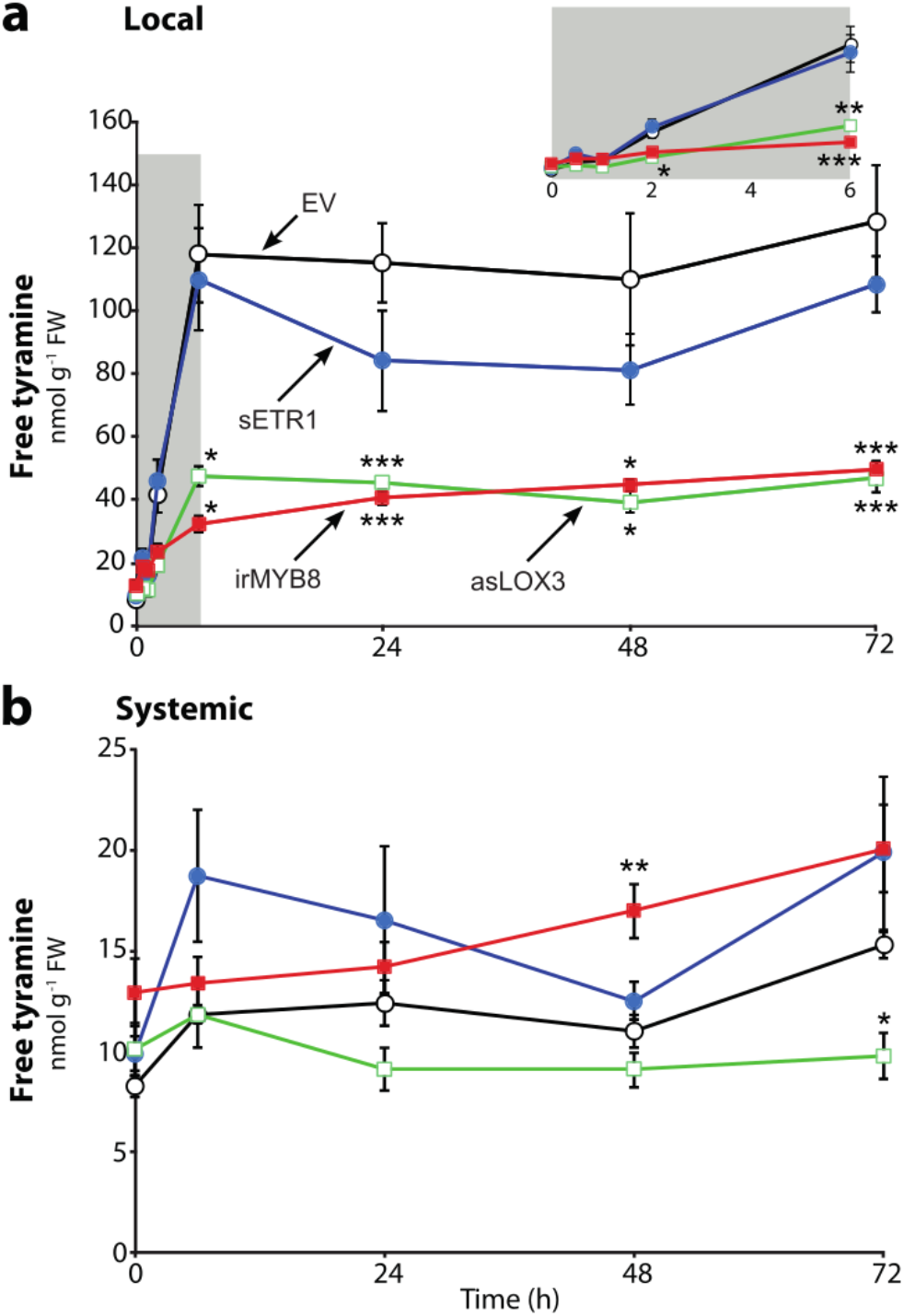
Local increases in tyramine levels upon simulated herbivory depend on jasmonate and MYB8-based signaling but not on ethylene signaling. Mean levels (± SE, 5 biological replicates) of free tyramine in local **(a)** and systemic **(b)** leaves of the empty vector (EV), sETR1, asLOX3 and irMYB8 transgenic plants during simulated herbivory. Grey insets represent close-up views on 0-to-6h accumulation dynamics in locally-treated leaves. Tyramine was quantified in acidified leaf extracts by HPLC-UV after derivatization with o-phthalaldehyde (OPDA) and fluorenylmethyloxycarbonyl chloride. Asterisks indicate significant differences with EV at respective time points (* P < 0.05, ** P < 0.01, *** P < 0.001, ANOVA followed by Tukey HSD post-hoc tests; when normality assumption was not met, Kruskal-Wallis and pairwise Wilcoxon rank sum tests were applied).

**Figure S6.**
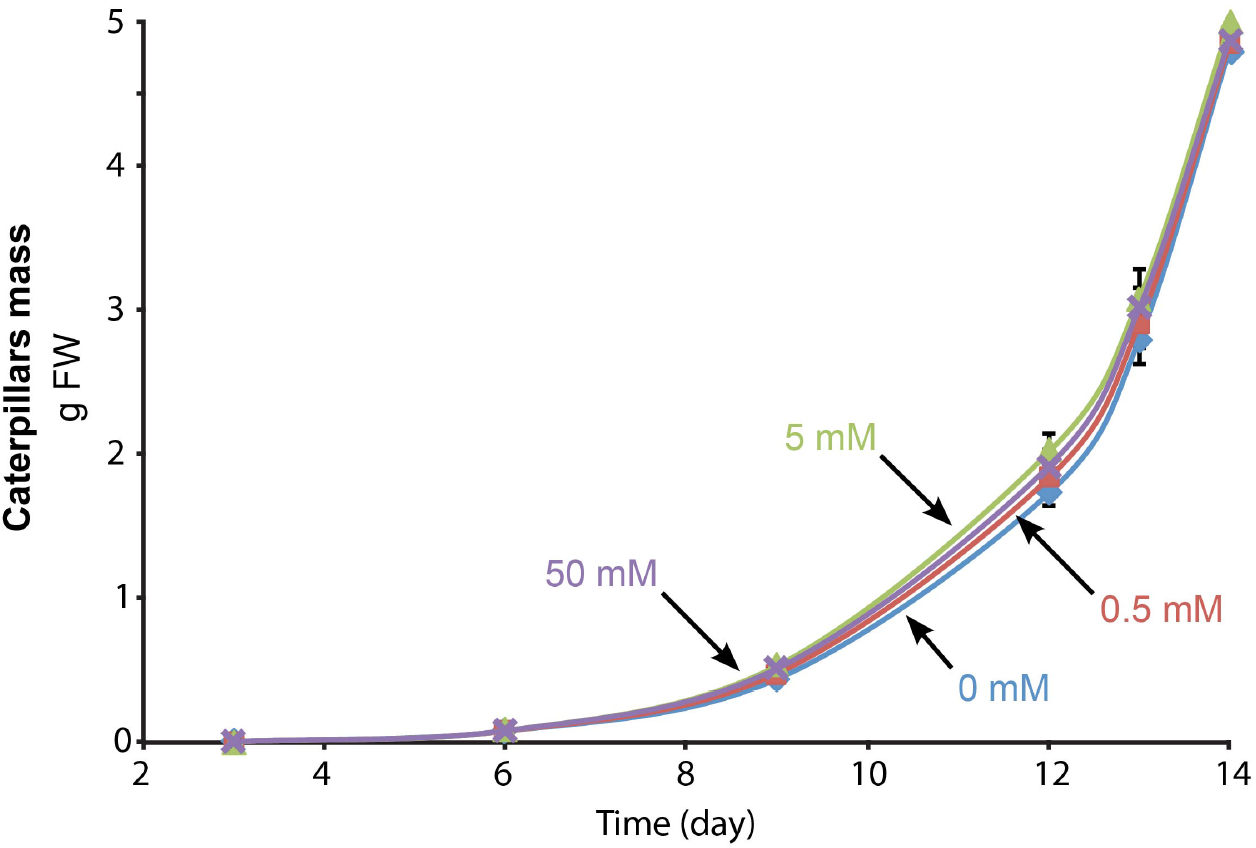
Addition of tyramine in artificial diet does not alter *Manduca sexta* larvae mass gain. Mean mass (± SE, 6 to 10 biological replicates) of *Manduca sexta* caterpillars fed on an artificial diet supplemented with different concentrations of tyramine.

