## Supplemental File S4 for "Ethylene is a local modulator of jasmonate-dependent phenolamide accumulation during *Manduca sexta* herbivory in *Nicotiana attenuata*"

### Statistical analysis of phytohormone levels

```
data = read.table("R_data.csv", header=T, sep=",", dec=".")

res_0 = subset(data, data$Time==0.0)
res_0.5 = subset(data, data$Time==0.5)
res_1 = subset(data, data$Time==1.0)
res_2 = subset(data, data$Time==2.0)

### Normality analysis

shapiro.test(subset(res_0$JA, res_0$Genotype=="EV")) # OK
##
##  Shapiro-Wilk normality test
##
## data:  subset(res_0$JA, res_0$Genotype == "EV")
## W = 0.92689, p-value = 0.5753
shapiro.test(subset(res_0$JA, res_0$Genotype=="ETR1")) # OK
##
##  Shapiro-Wilk normality test
##
## data:  subset(res_0$JA, res_0$Genotype == "ETR1")
## W = 0.92422, p-value = 0.5609
shapiro.test(subset(res_0$JA, res_0$Genotype=="LOX3")) # OK
##
##  Shapiro-Wilk normality test
##
## data:  subset(res_0$JA, res_0$Genotype == "LOX3")
## W = 0.89148, p-value = 0.3646
shapiro.test(subset(res_0$JA, res_0$Genotype=="MYB8")) # OK
##
##  Shapiro-Wilk normality test
##
## data:  subset(res_0$JA, res_0$Genotype == "MYB8")
## W = 0.9471, p-value = 0.7164
shapiro.test(subset(res_0.5$JA, res_0.5$Genotype=="EV")) # OK
##
##  Shapiro-Wilk normality test
##
## data:  subset(res_0.5$JA, res_0.5$Genotype == "EV")
## W = 0.97952, p-value = 0.932
shapiro.test(subset(res_0.5$JA, res_0.5$Genotype=="ETR1")) # OK
##
##  Shapiro-Wilk normality test
##
## data:  subset(res_0.5$JA, res_0.5$Genotype == "ETR1")
## W = 0.88292, p-value = 0.3513
shapiro.test(subset(res_0.5$JA, res_0.5$Genotype=="LOX3")) # NO
##
##  Shapiro-Wilk normality test
##
## data:  subset(res_0.5$JA, res_0.5$Genotype == "LOX3")
## W = 0.71159, p-value = 0.01261
ks.test(subset(res_0.5$JA, res_0.5$Genotype=="LOX3"), "pnorm") # NO
##
##  One-sample Kolmogorov-Smirnov test
##
## data:  subset(res_0.5$JA, res_0.5$Genotype == "LOX3")
## D = 1, p-value < 2.2e-16
## alternative hypothesis: two-sided
shapiro.test(subset(res_0.5$JA, res_0.5$Genotype=="MYB8"))# OK
##
##  Shapiro-Wilk normality test
##
## data:  subset(res_0.5$JA, res_0.5$Genotype == "MYB8")
## W = 0.83556, p-value = 0.153
shapiro.test(subset(res_1$JA, res_1$Genotype=="EV")) # OK
##
##  Shapiro-Wilk normality test
##
## data:  subset(res_1$JA, res_1$Genotype == "EV")
## W = 0.81807, p-value = 0.1128
shapiro.test(subset(res_1$JA, res_1$Genotype=="ETR1")) # OK
##
##  Shapiro-Wilk normality test
##
## data:  subset(res_1$JA, res_1$Genotype == "ETR1")
## W = 0.87758, p-value = 0.3284
shapiro.test(subset(res_1$JA, res_1$Genotype=="LOX3")) # NO
##
##  Shapiro-Wilk normality test
##
## data:  subset(res_1$JA, res_1$Genotype == "LOX3")
## W = 0.70375, p-value = 0.01048
shapiro.test(subset(res_1$JA, res_1$Genotype=="MYB8")) # OK
##
##  Shapiro-Wilk normality test
##
## data:  subset(res_1$JA, res_1$Genotype == "MYB8")
## W = 0.96912, p-value = 0.8696
shapiro.test(subset(res_2$JA, res_2$Genotype=="EV")) # OK
##
##  Shapiro-Wilk normality test
##
## data:  subset(res_2$JA, res_2$Genotype == "EV")
## W = 0.90879, p-value = 0.4604
shapiro.test(subset(res_2$JA, res_2$Genotype=="ETR1")) # OK
##
##  Shapiro-Wilk normality test
##
## data:  subset(res_2$JA, res_2$Genotype == "ETR1")
## W = 0.94757, p-value = 0.7009
shapiro.test(subset(res_2$JA, res_2$Genotype=="LOX3")) # OK
##
##  Shapiro-Wilk normality test
##
## data:  subset(res_2$JA, res_2$Genotype == "LOX3")
## W = 0.86422, p-value = 0.2438
shapiro.test(subset(res_2$JA, res_2$Genotype=="MYB8")) # OK
##
##  Shapiro-Wilk normality test
##
## data:  subset(res_2$JA, res_2$Genotype == "MYB8")
## W = 0.91751, p-value = 0.514
shapiro.test(subset(res_0$JA.Ile, res_0$Genotype=="EV")) # OK
##
##  Shapiro-Wilk normality test
##
## data:  subset(res_0$JA.Ile, res_0$Genotype == "EV")
## W = 0.93905, p-value = 0.6592
shapiro.test(subset(res_0$JA.Ile, res_0$Genotype=="ETR1")) # OK
##
##  Shapiro-Wilk normality test
##
## data:  subset(res_0$JA.Ile, res_0$Genotype == "ETR1")
## W = 0.81623, p-value = 0.1347
shapiro.test(subset(res_0$JA.Ile, res_0$Genotype=="LOX3")) # OK
##
##  Shapiro-Wilk normality test
##
## data:  subset(res_0$JA.Ile, res_0$Genotype == "LOX3")
## W = 0.84921, p-value = 0.192
shapiro.test(subset(res_0$JA.Ile, res_0$Genotype=="MYB8")) # OK
##
##  Shapiro-Wilk normality test
##
## data:  subset(res_0$JA.Ile, res_0$Genotype == "MYB8")
## W = 0.83641, p-value = 0.1552
shapiro.test(subset(res_0.5$JA.Ile, res_0.5$Genotype=="EV")) # OK
##
##  Shapiro-Wilk normality test
##
## data:  subset(res_0.5$JA.Ile, res_0.5$Genotype == "EV")
## W = 0.80685, p-value = 0.09204
shapiro.test(subset(res_0.5$JA.Ile, res_0.5$Genotype=="ETR1")) # OK
##
##  Shapiro-Wilk normality test
##
## data:  subset(res_0.5$JA.Ile, res_0.5$Genotype == "ETR1")
## W = 0.94816, p-value = 0.7046
shapiro.test(subset(res_0.5$JA.Ile, res_0.5$Genotype=="LOX3")) # OK
##
##  Shapiro-Wilk normality test
##
## data:  subset(res_0.5$JA.Ile, res_0.5$Genotype == "LOX3")
## W = 0.77871, p-value = 0.05373
shapiro.test(subset(res_0.5$JA.Ile, res_0.5$Genotype=="MYB8")) # NO
##
##  Shapiro-Wilk normality test
##
## data:  subset(res_0.5$JA.Ile, res_0.5$Genotype == "MYB8")
## W = 0.67779, p-value = 0.005528
shapiro.test(subset(res_1$JA.Ile, res_1$Genotype=="EV")) # OK
##
##  Shapiro-Wilk normality test
##
## data:  subset(res_1$JA.Ile, res_1$Genotype == "EV")
## W = 0.84215, p-value = 0.1709
shapiro.test(subset(res_1$JA.Ile, res_1$Genotype=="ETR1")) # OK
##
##  Shapiro-Wilk normality test
##
## data:  subset(res_1$JA.Ile, res_1$Genotype == "ETR1")
## W = 0.91637, p-value = 0.5168
shapiro.test(subset(res_1$JA.Ile, res_1$Genotype=="LOX3")) # OK
##
##  Shapiro-Wilk normality test
##
## data:  subset(res_1$JA.Ile, res_1$Genotype == "LOX3")
## W = 0.9776, p-value = 0.9214
shapiro.test(subset(res_1$JA.Ile, res_1$Genotype=="MYB8")) # OK
##
##  Shapiro-Wilk normality test
##
## data:  subset(res_1$JA.Ile, res_1$Genotype == "MYB8")
## W = 0.94781, p-value = 0.7216
shapiro.test(subset(res_2$JA.Ile, res_2$Genotype=="EV")) # OK
##
##  Shapiro-Wilk normality test
##
## data:  subset(res_2$JA.Ile, res_2$Genotype == "EV")
## W = 0.8497, p-value = 0.1936
shapiro.test(subset(res_2$JA.Ile, res_2$Genotype=="ETR1")) # OK
##
##  Shapiro-Wilk normality test
##
## data:  subset(res_2$JA.Ile, res_2$Genotype == "ETR1")
## W = 0.99595, p-value = 0.9856
shapiro.test(subset(res_2$JA.Ile, res_2$Genotype=="LOX3")) # OK
##
##  Shapiro-Wilk normality test
##
## data:  subset(res_2$JA.Ile, res_2$Genotype == "LOX3")
## W = 0.80886, p-value = 0.09551
shapiro.test(subset(res_2$JA.Ile, res_2$Genotype=="MYB8")) # OK
##
##  Shapiro-Wilk normality test
##
## data:  subset(res_2$JA.Ile, res_2$Genotype == "MYB8")
## W = 0.89565, p-value = 0.3863
shapiro.test(subset(res_0$OH.JA.Ile, res_0$Genotype=="EV")) # NO
##
##  Shapiro-Wilk normality test
##
## data:  subset(res_0$OH.JA.Ile, res_0$Genotype == "EV")
## W = 0.55218, p-value = 0.000131
shapiro.test(subset(res_0$OH.JA.Ile, res_0$Genotype=="ETR1")) # NO
##
##  Shapiro-Wilk normality test
##
## data:  subset(res_0$OH.JA.Ile, res_0$Genotype == "ETR1")
## W = 0.62978, p-value = 0.001241
shapiro.test(subset(res_0$OH.JA.Ile, res_0$Genotype=="MYB8")) # NO
##
##  Shapiro-Wilk normality test
##
## data:  subset(res_0$OH.JA.Ile, res_0$Genotype == "MYB8")
## W = 0.73489, p-value = 0.02144
shapiro.test(subset(res_0.5$OH.JA.Ile, res_0.5$Genotype=="EV")) # NO
##
##  Shapiro-Wilk normality test
##
## data:  subset(res_0.5$OH.JA.Ile, res_0.5$Genotype == "EV")
## W = 0.7728, p-value = 0.04774
shapiro.test(subset(res_0.5$OH.JA.Ile, res_0.5$Genotype=="ETR1")) # OK
##
##  Shapiro-Wilk normality test
##
## data:  subset(res_0.5$OH.JA.Ile, res_0.5$Genotype == "ETR1")
## W = 0.91379, p-value = 0.5027
shapiro.test(subset(res_0.5$OH.JA.Ile, res_0.5$Genotype=="LOX3")) # OK
##
##  Shapiro-Wilk normality test
##
## data:  subset(res_0.5$OH.JA.Ile, res_0.5$Genotype == "LOX3")
## W = 0.91009, p-value = 0.4682
shapiro.test(subset(res_0.5$OH.JA.Ile, res_0.5$Genotype=="MYB8")) # OK
##
##  Shapiro-Wilk normality test
##
## data:  subset(res_0.5$OH.JA.Ile, res_0.5$Genotype == "MYB8")
## W = 0.89084, p-value = 0.3613
shapiro.test(subset(res_1$OH.JA.Ile, res_1$Genotype=="EV")) # OK
##
##  Shapiro-Wilk normality test
##
## data:  subset(res_1$OH.JA.Ile, res_1$Genotype == "EV")
## W = 0.85369, p-value = 0.2065
shapiro.test(subset(res_1$OH.JA.Ile, res_1$Genotype=="ETR1")) # OK
##
##  Shapiro-Wilk normality test
##
## data:  subset(res_1$OH.JA.Ile, res_1$Genotype == "ETR1")
## W = 0.90889, p-value = 0.4765
shapiro.test(subset(res_1$OH.JA.Ile, res_1$Genotype=="LOX3")) # OK
##
##  Shapiro-Wilk normality test
##
## data:  subset(res_1$OH.JA.Ile, res_1$Genotype == "LOX3")
## W = 0.90192, p-value = 0.4206
shapiro.test(subset(res_1$OH.JA.Ile, res_1$Genotype=="MYB8")) # OK
##
##  Shapiro-Wilk normality test
##
## data:  subset(res_1$OH.JA.Ile, res_1$Genotype == "MYB8")
## W = 0.90794, p-value = 0.4553
shapiro.test(subset(res_2$OH.JA.Ile, res_2$Genotype=="EV")) # OK
##
##  Shapiro-Wilk normality test
##
## data:  subset(res_2$OH.JA.Ile, res_2$Genotype == "EV")
## W = 0.89688, p-value = 0.3929
shapiro.test(subset(res_2$OH.JA.Ile, res_2$Genotype=="ETR1")) # OK
##
##  Shapiro-Wilk normality test
##
## data:  subset(res_2$OH.JA.Ile, res_2$Genotype == "ETR1")
## W = 0.95527, p-value = 0.7491
shapiro.test(subset(res_2$OH.JA.Ile, res_2$Genotype=="LOX3")) # OK
##
##  Shapiro-Wilk normality test
##
## data:  subset(res_2$OH.JA.Ile, res_2$Genotype == "LOX3")
## W = 0.96809, p-value = 0.8629
shapiro.test(subset(res_2$OH.JA.Ile, res_2$Genotype=="MYB8")) # OK
##
##  Shapiro-Wilk normality test
##
## data:  subset(res_2$OH.JA.Ile, res_2$Genotype == "MYB8")
## W = 0.87888, p-value = 0.3043
shapiro.test(subset(res_0$ABA, res_0$Genotype=="EV")) # OK
##
##  Shapiro-Wilk normality test
##
## data:  subset(res_0$ABA, res_0$Genotype == "EV")
## W = 0.84539, p-value = 0.1804
shapiro.test(subset(res_0$ABA, res_0$Genotype=="ETR1")) # NO
##
##  Shapiro-Wilk normality test
##
## data:  subset(res_0$ABA, res_0$Genotype == "ETR1")
## W = 0.76149, p-value = 0.04919
shapiro.test(subset(res_0$ABA, res_0$Genotype=="LOX3")) # OK
##
##  Shapiro-Wilk normality test
##
## data:  subset(res_0$ABA, res_0$Genotype == "LOX3")
## W = 0.83979, p-value = 0.1643
shapiro.test(subset(res_0$ABA, res_0$Genotype=="MYB8")) # NO
##
##  Shapiro-Wilk normality test
##
## data:  subset(res_0$ABA, res_0$Genotype == "MYB8")
## W = 0.6885, p-value = 0.007232
shapiro.test(subset(res_0.5$ABA, res_0.5$Genotype=="EV")) # OK
##
##  Shapiro-Wilk normality test
##
## data:  subset(res_0.5$ABA, res_0.5$Genotype == "EV")
## W = 0.91628, p-value = 0.5062
shapiro.test(subset(res_0.5$ABA, res_0.5$Genotype=="ETR1")) # OK
##
##  Shapiro-Wilk normality test
##
## data:  subset(res_0.5$ABA, res_0.5$Genotype == "ETR1")
## W = 0.94879, p-value = 0.7086
shapiro.test(subset(res_0.5$ABA, res_0.5$Genotype=="LOX3")) # OK
##
##  Shapiro-Wilk normality test
##
## data:  subset(res_0.5$ABA, res_0.5$Genotype == "LOX3")
## W = 0.94637, p-value = 0.7113
shapiro.test(subset(res_0.5$ABA, res_0.5$Genotype=="MYB8")) # OK
##
##  Shapiro-Wilk normality test
##
## data:  subset(res_0.5$ABA, res_0.5$Genotype == "MYB8")
## W = 0.96929, p-value = 0.8707
shapiro.test(subset(res_1$ABA, res_1$Genotype=="EV")) # OK
##
##  Shapiro-Wilk normality test
##
## data:  subset(res_1$ABA, res_1$Genotype == "EV")
## W = 0.91955, p-value = 0.527
shapiro.test(subset(res_1$ABA, res_1$Genotype=="ETR1")) # OK
##
##  Shapiro-Wilk normality test
##
## data:  subset(res_1$ABA, res_1$Genotype == "ETR1")
## W = 0.86327, p-value = 0.2721
shapiro.test(subset(res_1$ABA, res_1$Genotype=="LOX3")) # OK
##
##  Shapiro-Wilk normality test
##
## data:  subset(res_1$ABA, res_1$Genotype == "LOX3")
## W = 0.87396, p-value = 0.2828
shapiro.test(subset(res_1$ABA, res_1$Genotype=="MYB8"))# OK
##
##  Shapiro-Wilk normality test
##
## data:  subset(res_1$ABA, res_1$Genotype == "MYB8")
## W = 0.93235, p-value = 0.6125
shapiro.test(subset(res_2$ABA, res_2$Genotype=="EV")) # NO
##
##  Shapiro-Wilk normality test
##
## data:  subset(res_2$ABA, res_2$Genotype == "EV")
## W = 0.77477, p-value = 0.04967
shapiro.test(subset(res_2$ABA, res_2$Genotype=="ETR1")) # OK
##
##  Shapiro-Wilk normality test
##
## data:  subset(res_2$ABA, res_2$Genotype == "ETR1")
## W = 0.88872, p-value = 0.3772
shapiro.test(subset(res_2$ABA, res_2$Genotype=="LOX3")) # OK
##
##  Shapiro-Wilk normality test
##
## data:  subset(res_2$ABA, res_2$Genotype == "LOX3")
## W = 0.88449, p-value = 0.3302
shapiro.test(subset(res_2$ABA, res_2$Genotype=="MYB8")) # OK
##
##  Shapiro-Wilk normality test
##
## data:  subset(res_2$ABA, res_2$Genotype == "MYB8")
## W = 0.95129, p-value = 0.7464
shapiro.test(subset(res_0$SA, res_0$Genotype=="EV")) # OK
##
##  Shapiro-Wilk normality test
##
## data:  subset(res_0$SA, res_0$Genotype == "EV")
## W = 0.971, p-value = 0.8817
shapiro.test(subset(res_0$SA, res_0$Genotype=="ETR1")) # OK
##
##  Shapiro-Wilk normality test
##
## data:  subset(res_0$SA, res_0$Genotype == "ETR1")
## W = 0.9435, p-value = 0.6758
shapiro.test(subset(res_0$SA, res_0$Genotype=="LOX3")) # OK
##
##  Shapiro-Wilk normality test
##
## data:  subset(res_0$SA, res_0$Genotype == "LOX3")
## W = 0.89539, p-value = 0.3849
shapiro.test(subset(res_0$SA, res_0$Genotype=="MYB8")) # OK
##
##  Shapiro-Wilk normality test
##
## data:  subset(res_0$SA, res_0$Genotype == "MYB8")
## W = 0.91827, p-value = 0.5188
shapiro.test(subset(res_0.5$SA, res_0.5$Genotype=="EV")) # OK
##
##  Shapiro-Wilk normality test
##
## data:  subset(res_0.5$SA, res_0.5$Genotype == "EV")
## W = 0.95516, p-value = 0.7739
shapiro.test(subset(res_0.5$SA, res_0.5$Genotype=="ETR1")) # OK
##
##  Shapiro-Wilk normality test
##
## data:  subset(res_0.5$SA, res_0.5$Genotype == "ETR1")
## W = 0.98253, p-value = 0.9167
shapiro.test(subset(res_0.5$SA, res_0.5$Genotype=="LOX3")) # OK
##
##  Shapiro-Wilk normality test
##
## data:  subset(res_0.5$SA, res_0.5$Genotype == "LOX3")
## W = 0.91798, p-value = 0.517
shapiro.test(subset(res_0.5$SA, res_0.5$Genotype=="MYB8")) # OK
##
##  Shapiro-Wilk normality test
##
## data:  subset(res_0.5$SA, res_0.5$Genotype == "MYB8")
## W = 0.88876, p-value = 0.3509
shapiro.test(subset(res_1$SA, res_1$Genotype=="EV")) # OK
##
##  Shapiro-Wilk normality test
##
## data:  subset(res_1$SA, res_1$Genotype == "EV")
## W = 0.91825, p-value = 0.5187
shapiro.test(subset(res_1$SA, res_1$Genotype=="ETR1")) # OK
##
##  Shapiro-Wilk normality test
##
## data:  subset(res_1$SA, res_1$Genotype == "ETR1")
## W = 0.95, p-value = 0.7161
shapiro.test(subset(res_1$SA, res_1$Genotype=="LOX3")) # OK
##
##  Shapiro-Wilk normality test
##
## data:  subset(res_1$SA, res_1$Genotype == "LOX3")
## W = 0.86602, p-value = 0.2507
shapiro.test(subset(res_1$SA, res_1$Genotype=="MYB8")) # OK
##
##  Shapiro-Wilk normality test
##
## data:  subset(res_1$SA, res_1$Genotype == "MYB8")
## W = 0.86379, p-value = 0.2422
shapiro.test(subset(res_2$SA, res_2$Genotype=="EV")) # OK
##
##  Shapiro-Wilk normality test
##
## data:  subset(res_2$SA, res_2$Genotype == "EV")
## W = 0.84804, p-value = 0.1884
shapiro.test(subset(res_2$SA, res_2$Genotype=="ETR1")) # OK
##
##  Shapiro-Wilk normality test
##
## data:  subset(res_2$SA, res_2$Genotype == "ETR1")
## W = 0.98393, p-value = 0.9247
shapiro.test(subset(res_2$SA, res_2$Genotype=="LOX3")) # OK
##
##  Shapiro-Wilk normality test
##
## data:  subset(res_2$SA, res_2$Genotype == "LOX3")
## W = 0.92301, p-value = 0.5496
shapiro.test(subset(res_2$SA, res_2$Genotype=="MYB8")) # OK
##
##  Shapiro-Wilk normality test
##
## data:  subset(res_2$SA, res_2$Genotype == "MYB8")
## W = 0.94713, p-value = 0.7167
### Variance analysis

JA_0 = formula(res_0$JA~res_0$Genotype)
JA.Ile_0 = formula(res_0$JA.Ile~res_0$Genotype)
OH.JA.Ile_0 = formula(res_0$OH.JA.Ile~res_0$Genotype)
ABA_0 = formula(res_0$ABA~res_0$Genotype)
SA_0 = formula(res_0$SA~res_0$Genotype)
JA_0.5 = formula(res_0.5$JA~res_0.5$Genotype)
JA.Ile_0.5 = formula(res_0.5$JA.Ile~res_0.5$Genotype)
OH.JA.Ile_0.5 = formula(res_0.5$OH.JA.Ile~res_0.5$Genotype)
ABA_0.5 = formula(res_0.5$ABA~res_0.5$Genotype)
SA_0.5 = formula(res_0.5$SA~res_0.5$Genotype)
JA_1 = formula(res_1$JA~res_1$Genotype)
JA.Ile_1 = formula(res_1$JA.Ile~res_1$Genotype)
OH.JA.Ile_1 = formula(res_1$OH.JA.Ile~res_1$Genotype)
ABA_1 = formula(res_1$ABA~res_1$Genotype)
SA_1 = formula(res_1$SA~res_1$Genotype)
JA_2 = formula(res_2$JA~res_2$Genotype)
JA.Ile_2 = formula(res_2$JA.Ile~res_2$Genotype)
OH.JA.Ile_2 = formula(res_2$OH.JA.Ile~res_2$Genotype)
ABA_2 = formula(res_2$ABA~res_2$Genotype)
SA_2 = formula(res_2$SA~res_2$Genotype)


fligner.test(JA_0) # OK
##
##  Fligner-Killeen test of homogeneity of variances
##
## data:  res_0$JA by res_0$Genotype
## Fligner-Killeen:med chi-squared = 4.1477, df = 3, p-value = 0.2459
fligner.test(JA.Ile_0) # OK
##
##  Fligner-Killeen test of homogeneity of variances
##
## data:  res_0$JA.Ile by res_0$Genotype
## Fligner-Killeen:med chi-squared = 0.50845, df = 3, p-value = 0.917
fligner.test(OH.JA.Ile_0) # OK
##
##  Fligner-Killeen test of homogeneity of variances
##
## data:  res_0$OH.JA.Ile by res_0$Genotype
## Fligner-Killeen:med chi-squared = 3.279, df = 3, p-value = 0.3506
fligner.test(ABA_0) # OK
##
##  Fligner-Killeen test of homogeneity of variances
##
## data:  res_0$ABA by res_0$Genotype
## Fligner-Killeen:med chi-squared = 0.85436, df = 3, p-value =
## 0.8364
fligner.test(SA_0) # OK
##
##  Fligner-Killeen test of homogeneity of variances
##
## data:  res_0$SA by res_0$Genotype
## Fligner-Killeen:med chi-squared = 3.321, df = 3, p-value = 0.3447
fligner.test(JA_0.5) # OK
##
##  Fligner-Killeen test of homogeneity of variances
##
## data:  res_0.5$JA by res_0.5$Genotype
## Fligner-Killeen:med chi-squared = 0.064607, df = 3, p-value =
## 0.9957
fligner.test(JA.Ile_0.5) # OK
##
##  Fligner-Killeen test of homogeneity of variances
##
## data:  res_0.5$JA.Ile by res_0.5$Genotype
## Fligner-Killeen:med chi-squared = 0.56792, df = 3, p-value =
## 0.9037
fligner.test(OH.JA.Ile_0.5) # OK
##
##  Fligner-Killeen test of homogeneity of variances
##
## data:  res_0.5$OH.JA.Ile by res_0.5$Genotype
## Fligner-Killeen:med chi-squared = 2.4158, df = 3, p-value = 0.4907
fligner.test(ABA_0.5) # OK
##
##  Fligner-Killeen test of homogeneity of variances
##
## data:  res_0.5$ABA by res_0.5$Genotype
## Fligner-Killeen:med chi-squared = 1.744, df = 3, p-value = 0.6272
fligner.test(SA_0.5) # OK
##
##  Fligner-Killeen test of homogeneity of variances
##
## data:  res_0.5$SA by res_0.5$Genotype
## Fligner-Killeen:med chi-squared = 1.3014, df = 3, p-value = 0.7288
fligner.test(JA_1) # OK
##
##  Fligner-Killeen test of homogeneity of variances
##
## data:  res_1$JA by res_1$Genotype
## Fligner-Killeen:med chi-squared = 5.8955, df = 3, p-value = 0.1168
fligner.test(JA.Ile_1) # OK
##
##  Fligner-Killeen test of homogeneity of variances
##
## data:  res_1$JA.Ile by res_1$Genotype
## Fligner-Killeen:med chi-squared = 3.7594, df = 3, p-value = 0.2886
fligner.test(OH.JA.Ile_1) # OK
##
##  Fligner-Killeen test of homogeneity of variances
##
## data:  res_1$OH.JA.Ile by res_1$Genotype
## Fligner-Killeen:med chi-squared = 4.2952, df = 3, p-value = 0.2313
fligner.test(ABA_1) # OK
##
##  Fligner-Killeen test of homogeneity of variances
##
## data:  res_1$ABA by res_1$Genotype
## Fligner-Killeen:med chi-squared = 0.59787, df = 3, p-value =
## 0.8969
fligner.test(SA_1) # OK
##
##  Fligner-Killeen test of homogeneity of variances
##
## data:  res_1$SA by res_1$Genotype
## Fligner-Killeen:med chi-squared = 2.9788, df = 3, p-value = 0.3949
fligner.test(JA_2) # OK
##
##  Fligner-Killeen test of homogeneity of variances
##
## data:  res_2$JA by res_2$Genotype
## Fligner-Killeen:med chi-squared = 6.7703, df = 3, p-value =
## 0.07959
fligner.test(JA.Ile_2) # OK
##
##  Fligner-Killeen test of homogeneity of variances
##
## data:  res_2$JA.Ile by res_2$Genotype
## Fligner-Killeen:med chi-squared = 4.4596, df = 3, p-value = 0.2159
fligner.test(OH.JA.Ile_2) # OK
##
##  Fligner-Killeen test of homogeneity of variances
##
## data:  res_2$OH.JA.Ile by res_2$Genotype
## Fligner-Killeen:med chi-squared = 4.6469, df = 3, p-value = 0.1996
fligner.test(ABA_2) # OK
##
##  Fligner-Killeen test of homogeneity of variances
##
## data:  res_2$ABA by res_2$Genotype
## Fligner-Killeen:med chi-squared = 3.4088, df = 3, p-value = 0.3328
fligner.test(SA_2) # OK
##
##  Fligner-Killeen test of homogeneity of variances
##
## data:  res_2$SA by res_2$Genotype
## Fligner-Killeen:med chi-squared = 0.67665, df = 3, p-value =
## 0.8787
### ANOVAs

summary(aov(JA_0))
##                Df Sum Sq Mean Sq F value Pr(>F)
## res_0$Genotype  3  94.29   31.43   2.697  0.083 .
## Residuals      15 174.78   11.65
## ---
## Signif. codes:  0 '***' 0.001 '**' 0.01 '*' 0.05 '.' 0.1 ' ' 1
summary(aov(JA.Ile_0))
##                Df Sum Sq Mean Sq F value Pr(>F)
## res_0$Genotype  3 0.0430 0.01434   0.416  0.744
## Residuals      15 0.5169 0.03446
kruskal.test(OH.JA.Ile_0)
##
##  Kruskal-Wallis rank sum test
##
## data:  res_0$OH.JA.Ile by res_0$Genotype
## Kruskal-Wallis chi-squared = 2.8086, df = 3, p-value = 0.4221
kruskal.test(ABA_0)
##
##  Kruskal-Wallis rank sum test
##
## data:  res_0$ABA by res_0$Genotype
## Kruskal-Wallis chi-squared = 11.28, df = 3, p-value = 0.0103
pairwise.wilcox.test(res_0$ABA, res_0$Genotype)
##
##  Pairwise comparisons using Wilcoxon rank sum test
##
## data:  res_0$ABA and res_0$Genotype
##
##      ETR1  EV    LOX3
## EV   0.571 -     -
## LOX3 1.000 1.000 -
## MYB8 0.063 0.048 0.048
##
## P value adjustment method: holm
summary(aov(SA_0))
##                Df Sum Sq Mean Sq F value Pr(>F)
## res_0$Genotype  3  642.2  214.07     2.3  0.119
## Residuals      15 1396.2   93.08
kruskal.test(JA_0.5)
##
##  Kruskal-Wallis rank sum test
##
## data:  res_0.5$JA by res_0.5$Genotype
## Kruskal-Wallis chi-squared = 9.0316, df = 3, p-value = 0.02887
pairwise.wilcox.test(res_0.5$JA, res_0.5$Genotype)
##
##  Pairwise comparisons using Wilcoxon rank sum test
##
## data:  res_0.5$JA and res_0.5$Genotype
##
##      ETR1  EV    LOX3
## EV   0.603 -     -
## LOX3 0.159 0.095 -
## MYB8 0.603 0.603 0.603
##
## P value adjustment method: holm
kruskal.test(JA.Ile_0.5)
##
##  Kruskal-Wallis rank sum test
##
## data:  res_0.5$JA.Ile by res_0.5$Genotype
## Kruskal-Wallis chi-squared = 4.2568, df = 3, p-value = 0.235
kruskal.test(OH.JA.Ile_0.5)
##
##  Kruskal-Wallis rank sum test
##
## data:  res_0.5$OH.JA.Ile by res_0.5$Genotype
## Kruskal-Wallis chi-squared = 4.4605, df = 3, p-value = 0.2158
summary(aov(ABA_0.5))
##                  Df Sum Sq Mean Sq F value Pr(>F)
## res_0.5$Genotype  3   1165   388.4   1.312  0.307
## Residuals        15   4440   296.0
summary(aov(SA_0.5))
##                  Df Sum Sq Mean Sq F value Pr(>F)
## res_0.5$Genotype  3  834.4   278.1   2.254  0.124
## Residuals        15 1851.4   123.4
kruskal.test(JA_1)
##
##  Kruskal-Wallis rank sum test
##
## data:  res_1$JA by res_1$Genotype
## Kruskal-Wallis chi-squared = 15.829, df = 3, p-value = 0.001229
pairwise.wilcox.test(res_1$JA, res_1$Genotype)
##
##  Pairwise comparisons using Wilcoxon rank sum test
##
## data:  res_1$JA and res_1$Genotype
##
##      ETR1  EV    LOX3
## EV   0.111 -     -
## LOX3 0.063 0.048 -
## MYB8 0.063 0.063 0.048
##
## P value adjustment method: holm
summary(aov(JA.Ile_1))
##                Df Sum Sq Mean Sq F value   Pr(>F)
## res_1$Genotype  3  56708   18903   56.76 2.03e-08 ***
## Residuals      15   4995     333
## ---
## Signif. codes:  0 '***' 0.001 '**' 0.01 '*' 0.05 '.' 0.1 ' ' 1
TukeyHSD(aov(JA.Ile_1))
##   Tukey multiple comparisons of means
##     95% family-wise confidence level
##
## Fit: aov(formula = JA.Ile_1)
##
## $`res_1$Genotype`
##                  diff         lwr       upr     p adj
## EV-ETR1     37.960322    2.678004  73.24264 0.0330086
## LOX3-ETR1  -88.023449 -123.305768 -52.74113 0.0000168
## MYB8-ETR1   46.432554   11.150236  81.71487 0.0085528
## LOX3-EV   -125.983771 -159.248260 -92.71928 0.0000001
## MYB8-EV      8.472232  -24.792256  41.73672 0.8818918
## MYB8-LOX3  134.456004  101.191515 167.72049 0.0000000
summary(aov(OH.JA.Ile_1))
##                Df Sum Sq Mean Sq F value  Pr(>F)
## res_1$Genotype  3   8317  2772.3   7.544 0.00263 **
## Residuals      15   5512   367.5
## ---
## Signif. codes:  0 '***' 0.001 '**' 0.01 '*' 0.05 '.' 0.1 ' ' 1
TukeyHSD(aov(OH.JA.Ile_1))
##   Tukey multiple comparisons of means
##     95% family-wise confidence level
##
## Fit: aov(formula = OH.JA.Ile_1)
##
## $`res_1$Genotype`
##                 diff       lwr       upr     p adj
## EV-ETR1     2.007294 -35.05453 39.069119 0.9985824
## LOX3-ETR1 -34.821598 -71.88342  2.240226 0.0690960
## MYB8-ETR1  21.870489 -15.19134 58.932313 0.3572606
## LOX3-EV   -36.828892 -71.77112 -1.886669 0.0372445
## MYB8-EV    19.863194 -15.07903 54.805417 0.3881690
## MYB8-LOX3  56.692086  21.74986 91.634309 0.0015176
summary(aov(ABA_1))
##                Df Sum Sq Mean Sq F value   Pr(>F)
## res_1$Genotype  3   7898  2632.8   11.92 0.000298 ***
## Residuals      15   3313   220.9
## ---
## Signif. codes:  0 '***' 0.001 '**' 0.01 '*' 0.05 '.' 0.1 ' ' 1
TukeyHSD(aov(ABA_1))
##   Tukey multiple comparisons of means
##     95% family-wise confidence level
##
## Fit: aov(formula = ABA_1)
##
## $`res_1$Genotype`
##                 diff        lwr       upr     p adj
## EV-ETR1   -16.522927 -45.257132 12.211277 0.3786340
## LOX3-ETR1 -25.185830 -53.920034  3.548375 0.0959722
## MYB8-ETR1  27.004099  -1.730106 55.738303 0.0690101
## LOX3-EV    -8.662902 -35.753770 18.427965 0.7938493
## MYB8-EV    43.527026  16.436159 70.617894 0.0016569
## MYB8-LOX3  52.189928  25.099061 79.280796 0.0002897
summary(aov(SA_1))
##                Df Sum Sq Mean Sq F value Pr(>F)
## res_1$Genotype  3   1426   475.4   2.784  0.077 .
## Residuals      15   2561   170.8
## ---
## Signif. codes:  0 '***' 0.001 '**' 0.01 '*' 0.05 '.' 0.1 ' ' 1
summary(aov(JA_2))
##                Df Sum Sq Mean Sq F value   Pr(>F)
## res_2$Genotype  3 328976  109659   11.97 0.000292 ***
## Residuals      15 137418    9161
## ---
## Signif. codes:  0 '***' 0.001 '**' 0.01 '*' 0.05 '.' 0.1 ' ' 1
TukeyHSD(aov(JA_2))
##   Tukey multiple comparisons of means
##     95% family-wise confidence level
##
## Fit: aov(formula = JA_2)
##
## $`res_2$Genotype`
##                 diff        lwr        upr     p adj
## EV-ETR1     91.85235  -93.20178 276.906485 0.5007365
## LOX3-ETR1 -176.27907 -361.33320   8.775063 0.0644842
## MYB8-ETR1  167.65033  -17.40381 352.704461 0.0824230
## LOX3-EV   -268.13142 -442.60213 -93.660711 0.0024508
## MYB8-EV     75.79798  -98.67273 250.268687 0.6051936
## MYB8-LOX3  343.92940  169.45869 518.400108 0.0002286
summary(aov(JA.Ile_2))
##                Df Sum Sq Mean Sq F value  Pr(>F)
## res_2$Genotype  3  165.9   55.29   7.351 0.00294 **
## Residuals      15  112.8    7.52
## ---
## Signif. codes:  0 '***' 0.001 '**' 0.01 '*' 0.05 '.' 0.1 ' ' 1
TukeyHSD(aov(JA.Ile_2))
##   Tukey multiple comparisons of means
##     95% family-wise confidence level
##
## Fit: aov(formula = JA.Ile_2)
##
## $`res_2$Genotype`
##                diff        lwr        upr     p adj
## EV-ETR1    1.350066  -3.952090  6.6522222 0.8819714
## LOX3-ETR1 -4.892930 -10.195087  0.4092259 0.0754655
## MYB8-ETR1  2.751854  -2.550302  8.0540101 0.4639783
## LOX3-EV   -6.242996 -11.241917 -1.2440755 0.0125167
## MYB8-EV    1.401788  -3.597133  6.4007087 0.8495960
## MYB8-LOX3  7.644784   2.645863 12.6437050 0.0025570
summary(aov(OH.JA.Ile_2))
##                Df Sum Sq Mean Sq F value   Pr(>F)
## res_2$Genotype  3   4065  1355.1   15.35 7.69e-05 ***
## Residuals      15   1324    88.3
## ---
## Signif. codes:  0 '***' 0.001 '**' 0.01 '*' 0.05 '.' 0.1 ' ' 1
TukeyHSD(aov(OH.JA.Ile_2))
##   Tukey multiple comparisons of means
##     95% family-wise confidence level
##
## Fit: aov(formula = OH.JA.Ile_2)
##
## $`res_2$Genotype`
##                 diff        lwr       upr     p adj
## EV-ETR1     9.851469  -8.315115 28.018052 0.4274544
## LOX3-ETR1 -14.739314 -32.905897  3.427269 0.1332127
## MYB8-ETR1  24.492440   6.325857 42.659023 0.0071245
## LOX3-EV   -24.590783 -41.718401 -7.463164 0.0043365
## MYB8-EV    14.640971  -2.486647 31.768590 0.1071959
## MYB8-LOX3  39.231754  22.104135 56.359373 0.0000449
kruskal.test(ABA_2)
##
##  Kruskal-Wallis rank sum test
##
## data:  res_2$ABA by res_2$Genotype
## Kruskal-Wallis chi-squared = 14.116, df = 3, p-value = 0.002752
pairwise.wilcox.test(res_2$ABA, res_2$Genotype)
##
##  Pairwise comparisons using Wilcoxon rank sum test
##
## data:  res_2$ABA and res_2$Genotype
##
##      ETR1  EV    LOX3
## EV   0.381 -     -
## LOX3 0.730 0.048 -
## MYB8 0.048 0.048 0.048
##
## P value adjustment method: holm
summary(aov(SA_2))
##                Df Sum Sq Mean Sq F value Pr(>F)
## res_2$Genotype  3   9663    3221   0.855  0.486
## Residuals      15  56518    3768

data = read.table("R_data.csv", header=T, sep=",", dec=".")

## Normality analysis

shapiro.test(subset(data$ET, data$Genotype=="EV"))
##
##  Shapiro-Wilk normality test
##
## data:  subset(data$ET, data$Genotype == "EV")
## W = 0.86379, p-value = 0.2422
shapiro.test(subset(data$ET, data$Genotype=="ETR1"))
##
##  Shapiro-Wilk normality test
##
## data:  subset(data$ET, data$Genotype == "ETR1")
## W = 0.81379, p-value = 0.1045
shapiro.test(subset(data$ET, data$Genotype=="LOX3"))
##
##  Shapiro-Wilk normality test
##
## data:  subset(data$ET, data$Genotype == "LOX3")
## W = 0.92168, p-value = 0.5408
shapiro.test(subset(data$ET, data$Genotype=="MYB8"))
##
##  Shapiro-Wilk normality test
##
## data:  subset(data$ET, data$Genotype == "MYB8")
## W = 0.81488, p-value = 0.1065
shapiro.test(subset(data$ET, data$Genotype=="ACO"))
##
##  Shapiro-Wilk normality test
##
## data:  subset(data$ET, data$Genotype == "ACO")
## W = 0.97644, p-value = 0.9147
## Variance analysis

ET = formula(data$ET~data$Genotype)
fligner.test(ET)
##
##  Fligner-Killeen test of homogeneity of variances
##
## data:  data$ET by data$Genotype
## Fligner-Killeen:med chi-squared = 8.0973, df = 4, p-value =
## 0.08808
## ANOVA

summary(aov(ET))
##               Df Sum Sq Mean Sq F value   Pr(>F)
## data$Genotype  4  96616   24154   12.89 2.38e-05 ***
## Residuals     20  37474    1874
## ---
## Signif. codes:  0 '***' 0.001 '**' 0.01 '*' 0.05 '.' 0.1 ' ' 1
TukeyHSD(aov(ET))
##   Tukey multiple comparisons of means
##     95% family-wise confidence level
##
## Fit: aov(formula = ET)
##
## $`data$Genotype`
##                  diff         lwr       upr     p adj
## ETR1-ACO   193.247353  111.326083 275.16862 0.0000069
## EV-ACO      72.661999   -9.259271 154.58327 0.0977160
## LOX3-ACO    69.985604  -11.935666 151.90687 0.1174930
## MYB8-ACO    86.988661    5.067391 168.90993 0.0340508
## EV-ETR1   -120.585354 -202.506624 -38.66408 0.0022625
## LOX3-ETR1 -123.261749 -205.183019 -41.34048 0.0018136
## MYB8-ETR1 -106.258692 -188.179961 -24.33742 0.0073485
## LOX3-EV     -2.676395  -84.597665  79.24487 0.9999774
## MYB8-EV     14.326662  -67.594608  96.24793 0.9839194
## MYB8-LOX3   17.003057  -64.918212  98.92433 0.9699496
```

### Statistical analysis of phenolamide levels by HPLC-UV

#### Local leaves

```
data = read.table("R_data_local_sansMYB8.csv", header=T, sep=",", dec=".")

res_0 = subset(data, data$Time==0)
res_24 = subset(data, data$Time==24)
res_48 = subset(data, data$Time==48)
res_72 = subset(data, data$Time==72)

### Tests de normalite

shapiro.test(subset(res_0$CP, res_0$Genotype=="EV")) # OK
##
##  Shapiro-Wilk normality test
##
## data:  subset(res_0$CP, res_0$Genotype == "EV")
## W = 0.93344, p-value = 0.62
shapiro.test(subset(res_0$CP, res_0$Genotype=="ETR1")) # OK
##
##  Shapiro-Wilk normality test
##
## data:  subset(res_0$CP, res_0$Genotype == "ETR1")
## W = 0.90608, p-value = 0.4444
shapiro.test(subset(res_0$CP, res_0$Genotype=="LOX3")) # OK
##
##  Shapiro-Wilk normality test
##
## data:  subset(res_0$CP, res_0$Genotype == "LOX3")
## W = 0.97535, p-value = 0.9083
shapiro.test(subset(res_24$CP, res_24$Genotype=="EV")) # OK
##
##  Shapiro-Wilk normality test
##
## data:  subset(res_24$CP, res_24$Genotype == "EV")
## W = 0.82692, p-value = 0.1319
shapiro.test(subset(res_24$CP, res_24$Genotype=="ETR1")) # OK
##
##  Shapiro-Wilk normality test
##
## data:  subset(res_24$CP, res_24$Genotype == "ETR1")
## W = 0.83982, p-value = 0.1644
shapiro.test(subset(res_48$CP, res_48$Genotype=="EV")) # OK
##
##  Shapiro-Wilk normality test
##
## data:  subset(res_48$CP, res_48$Genotype == "EV")
## W = 0.86109, p-value = 0.2322
shapiro.test(subset(res_48$CP, res_48$Genotype=="ETR1")) # OK
##
##  Shapiro-Wilk normality test
##
## data:  subset(res_48$CP, res_48$Genotype == "ETR1")
## W = 0.83439, p-value = 0.15
shapiro.test(subset(res_48$CP, res_48$Genotype=="LOX3")) # NO
##
##  Shapiro-Wilk normality test
##
## data:  subset(res_48$CP, res_48$Genotype == "LOX3")
## W = 0.55218, p-value = 0.000131
shapiro.test(subset(res_72$CP, res_72$Genotype=="EV")) # OK
##
##  Shapiro-Wilk normality test
##
## data:  subset(res_72$CP, res_72$Genotype == "EV")
## W = 0.85896, p-value = 0.2245
shapiro.test(subset(res_72$CP, res_72$Genotype=="ETR1")) # OK
##
##  Shapiro-Wilk normality test
##
## data:  subset(res_72$CP, res_72$Genotype == "ETR1")
## W = 0.89077, p-value = 0.361
shapiro.test(subset(res_72$CP, res_72$Genotype=="LOX3")) # NO
##
##  Shapiro-Wilk normality test
##
## data:  subset(res_72$CP, res_72$Genotype == "LOX3")
## W = 0.70291, p-value = 0.01027
shapiro.test(subset(res_0$DCSs, res_0$Genotype=="EV"))  # OK
##
##  Shapiro-Wilk normality test
##
## data:  subset(res_0$DCSs, res_0$Genotype == "EV")
## W = 0.92548, p-value = 0.5659
shapiro.test(subset(res_0$DCSs, res_0$Genotype=="ETR1")) # OK
##
##  Shapiro-Wilk normality test
##
## data:  subset(res_0$DCSs, res_0$Genotype == "ETR1")
## W = 0.93465, p-value = 0.6284
shapiro.test(subset(res_0$DCSs, res_0$Genotype=="LOX3")) # OK
##
##  Shapiro-Wilk normality test
##
## data:  subset(res_0$DCSs, res_0$Genotype == "LOX3")
## W = 0.89071, p-value = 0.3607
shapiro.test(subset(res_24$DCSs, res_24$Genotype=="EV")) # OK
##
##  Shapiro-Wilk normality test
##
## data:  subset(res_24$DCSs, res_24$Genotype == "EV")
## W = 0.86849, p-value = 0.2604
shapiro.test(subset(res_24$DCSs, res_24$Genotype=="ETR1")) # OK
##
##  Shapiro-Wilk normality test
##
## data:  subset(res_24$DCSs, res_24$Genotype == "ETR1")
## W = 0.893, p-value = 0.3724
shapiro.test(subset(res_24$DCSs, res_24$Genotype=="LOX3")) # OK
##
##  Shapiro-Wilk normality test
##
## data:  subset(res_24$DCSs, res_24$Genotype == "LOX3")
## W = 0.97037, p-value = 0.8776
shapiro.test(subset(res_48$DCSs, res_48$Genotype=="EV")) # OK
##
##  Shapiro-Wilk normality test
##
## data:  subset(res_48$DCSs, res_48$Genotype == "EV")
## W = 0.94811, p-value = 0.7237
shapiro.test(subset(res_48$DCSs, res_48$Genotype=="ETR1")) # OK
##
##  Shapiro-Wilk normality test
##
## data:  subset(res_48$DCSs, res_48$Genotype == "ETR1")
## W = 0.7772, p-value = 0.05215
shapiro.test(subset(res_48$DCSs, res_48$Genotype=="LOX3")) # NO
##
##  Shapiro-Wilk normality test
##
## data:  subset(res_48$DCSs, res_48$Genotype == "LOX3")
## W = 0.7251, p-value = 0.01722
shapiro.test(subset(res_72$DCSs, res_72$Genotype=="EV")) # OK
##
##  Shapiro-Wilk normality test
##
## data:  subset(res_72$DCSs, res_72$Genotype == "EV")
## W = 0.95834, p-value = 0.7964
shapiro.test(subset(res_72$DCSs, res_72$Genotype=="ETR1")) # NO
##
##  Shapiro-Wilk normality test
##
## data:  subset(res_72$DCSs, res_72$Genotype == "ETR1")
## W = 0.94584, p-value = 0.7075
shapiro.test(subset(res_72$DCSs, res_72$Genotype=="LOX3")) # NO
##
##  Shapiro-Wilk normality test
##
## data:  subset(res_72$DCSs, res_72$Genotype == "LOX3")
## W = 0.65366, p-value = 0.00294
### Tests de variances

CP_0 = formula(res_0$CP~res_0$Genotype)
DCSs_0 = formula(res_0$DCSs~res_0$Genotype)
CP_24 = formula(res_24$CP~res_24$Genotype)
DCSs_24 = formula(res_24$DCSs~res_24$Genotype)
CP_48 = formula(res_48$CP~res_48$Genotype)
DCSs_48 = formula(res_48$DCSs~res_48$Genotype)
CP_72 = formula(res_72$CP~res_72$Genotype)
DCSs_72 = formula(res_72$DCSs~res_72$Genotype)

fligner.test(CP_0) # OK
##
##  Fligner-Killeen test of homogeneity of variances
##
## data:  res_0$CP by res_0$Genotype
## Fligner-Killeen:med chi-squared = 4.5942, df = 2, p-value = 0.1005
fligner.test(CP_24) # OK
##
##  Fligner-Killeen test of homogeneity of variances
##
## data:  res_24$CP by res_24$Genotype
## Fligner-Killeen:med chi-squared = 5.8465, df = 2, p-value =
## 0.05376
fligner.test(CP_48) # OK
##
##  Fligner-Killeen test of homogeneity of variances
##
## data:  res_48$CP by res_48$Genotype
## Fligner-Killeen:med chi-squared = 4.6345, df = 2, p-value =
## 0.09854
fligner.test(CP_72) # OK
##
##  Fligner-Killeen test of homogeneity of variances
##
## data:  res_72$CP by res_72$Genotype
## Fligner-Killeen:med chi-squared = 3.5309, df = 2, p-value = 0.1711
fligner.test(DCSs_0) # OK
##
##  Fligner-Killeen test of homogeneity of variances
##
## data:  res_0$DCSs by res_0$Genotype
## Fligner-Killeen:med chi-squared = 2.161, df = 2, p-value = 0.3394
fligner.test(DCSs_24) # OK
##
##  Fligner-Killeen test of homogeneity of variances
##
## data:  res_24$DCSs by res_24$Genotype
## Fligner-Killeen:med chi-squared = 2.7347, df = 2, p-value = 0.2548
fligner.test(DCSs_48) # OK
##
##  Fligner-Killeen test of homogeneity of variances
##
## data:  res_48$DCSs by res_48$Genotype
## Fligner-Killeen:med chi-squared = 0.11508, df = 2, p-value =
## 0.9441
fligner.test(DCSs_72) # OK
##
##  Fligner-Killeen test of homogeneity of variances
##
## data:  res_72$DCSs by res_72$Genotype
## Fligner-Killeen:med chi-squared = 1.1076, df = 2, p-value = 0.5747
### ANOVAs

summary(aov(CP_0))
##                Df Sum Sq Mean Sq F value Pr(>F)
## res_0$Genotype  2  62.21  31.105   6.438 0.0126 *
## Residuals      12  57.98   4.832
## ---
## Signif. codes:  0 '***' 0.001 '**' 0.01 '*' 0.05 '.' 0.1 ' ' 1
TukeyHSD(aov(CP_0))
##   Tukey multiple comparisons of means
##     95% family-wise confidence level
##
## Fit: aov(formula = CP_0)
##
## $`res_0$Genotype`
##                diff       lwr        upr     p adj
## EV-ETR1    1.227502 -2.481329  4.9363330 0.6607156
## LOX3-ETR1 -3.573510 -7.282341  0.1353204 0.0592910
## LOX3-EV   -4.801013 -8.509843 -1.0921819 0.0122807
summary(aov(DCSs_0))
##                Df Sum Sq Mean Sq F value Pr(>F)
## res_0$Genotype  2  12986    6493   0.662  0.534
## Residuals      12 117771    9814
summary(aov(CP_24))
##                 Df Sum Sq Mean Sq F value   Pr(>F)
## res_24$Genotype  2  15767    7883   15.11 0.000528 ***
## Residuals       12   6262     522
## ---
## Signif. codes:  0 '***' 0.001 '**' 0.01 '*' 0.05 '.' 0.1 ' ' 1
TukeyHSD(aov(CP_24))
##   Tukey multiple comparisons of means
##     95% family-wise confidence level
##
## Fit: aov(formula = CP_24)
##
## $`res_24$Genotype`
##            diff         lwr         upr     p adj
## EV-ETR1    41.0    2.454525  79.5454753 0.0370263
## LOX3-ETR1 -38.4  -76.945475   0.1454753 0.0508926
## LOX3-EV   -79.4 -117.945475 -40.8545247 0.0003731
summary(aov(DCSs_24))
##                 Df Sum Sq Mean Sq F value   Pr(>F)
## res_24$Genotype  2  35066   17533   25.27 4.99e-05 ***
## Residuals       12   8326     694
## ---
## Signif. codes:  0 '***' 0.001 '**' 0.01 '*' 0.05 '.' 0.1 ' ' 1
TukeyHSD(aov(DCSs_24))
##   Tukey multiple comparisons of means
##     95% family-wise confidence level
##
## Fit: aov(formula = DCSs_24)
##
## $`res_24$Genotype`
##             diff        lwr       upr     p adj
## EV-ETR1     61.6   17.15517 106.04483 0.0079301
## LOX3-ETR1  -56.8 -101.24483 -12.35517 0.0132905
## LOX3-EV   -118.4 -162.84483 -73.95517 0.0000341
kruskal.test(CP_48)
##
##  Kruskal-Wallis rank sum test
##
## data:  res_48$CP by res_48$Genotype
## Kruskal-Wallis chi-squared = 9.9578, df = 2, p-value = 0.006882
pairwise.wilcox.test(res_48$CP, res_48$Genotype)
## Warning in wilcox.test.default(xi, xj, paired = paired, ...): cannot
## compute exact p-value with ties

## Warning in wilcox.test.default(xi, xj, paired = paired, ...): cannot
## compute exact p-value with ties
##
##  Pairwise comparisons using Wilcoxon rank sum test
##
## data:  res_48$CP and res_48$Genotype
##
##      ETR1  EV
## EV   0.421 -
## LOX3 0.029 0.029
##
## P value adjustment method: holm
kruskal.test(DCSs_48)
##
##  Kruskal-Wallis rank sum test
##
## data:  res_48$DCSs by res_48$Genotype
## Kruskal-Wallis chi-squared = 11.58, df = 2, p-value = 0.003058
pairwise.wilcox.test(res_48$DCSs, res_48$Genotype)
##
##  Pairwise comparisons using Wilcoxon rank sum test
##
## data:  res_48$DCSs and res_48$Genotype
##
##      ETR1  EV
## EV   0.024 -
## LOX3 0.032 0.024
##
## P value adjustment method: holm
kruskal.test(CP_72)
##
##  Kruskal-Wallis rank sum test
##
## data:  res_72$CP by res_72$Genotype
## Kruskal-Wallis chi-squared = 11.58, df = 2, p-value = 0.003058
pairwise.wilcox.test(res_72$CP, res_72$Genotype)
##
##  Pairwise comparisons using Wilcoxon rank sum test
##
## data:  res_72$CP and res_72$Genotype
##
##      ETR1  EV
## EV   0.032 -
## LOX3 0.024 0.024
##
## P value adjustment method: holm
kruskal.test(DCSs_72)
##
##  Kruskal-Wallis rank sum test
##
## data:  res_72$DCSs by res_72$Genotype
## Kruskal-Wallis chi-squared = 9.62, df = 2, p-value = 0.008148
pairwise.wilcox.test(res_72$DCSs, res_72$Genotype)
##
##  Pairwise comparisons using Wilcoxon rank sum test
##
## data:  res_72$DCSs and res_72$Genotype
##
##      ETR1  EV
## EV   0.024 -
## LOX3 0.151 0.032
##
## P value adjustment method: holm
```

#### Systemic leaves

```
data = read.table("R_data_systemic_sansMYB8.csv", header=T, sep=",", dec=".")

res_24 = subset(data, data$Time==24)
res_48 = subset(data, data$Time==48)
res_72 = subset(data, data$Time==72)

### Tests de normalite


shapiro.test(subset(res_24$CP, res_24$Genotype=="EV")) # OK
##
##  Shapiro-Wilk normality test
##
## data:  subset(res_24$CP, res_24$Genotype == "EV")
## W = 0.89786, p-value = 0.3982
shapiro.test(subset(res_24$CP, res_24$Genotype=="ETR1")) # NO
##
##  Shapiro-Wilk normality test
##
## data:  subset(res_24$CP, res_24$Genotype == "ETR1")
## W = 0.59576, p-value = 0.0005497
shapiro.test(subset(res_24$CP, res_24$Genotype=="LOX3")) # OK
##
##  Shapiro-Wilk normality test
##
## data:  subset(res_24$CP, res_24$Genotype == "LOX3")
## W = 0.8707, p-value = 0.2693
shapiro.test(subset(res_48$CP, res_48$Genotype=="EV")) # OK
##
##  Shapiro-Wilk normality test
##
## data:  subset(res_48$CP, res_48$Genotype == "EV")
## W = 0.92857, p-value = 0.5867
shapiro.test(subset(res_48$CP, res_48$Genotype=="ETR1")) # NO
##
##  Shapiro-Wilk normality test
##
## data:  subset(res_48$CP, res_48$Genotype == "ETR1")
## W = 0.75073, p-value = 0.03023
shapiro.test(subset(res_48$CP, res_48$Genotype=="LOX3")) # NO
##
##  Shapiro-Wilk normality test
##
## data:  subset(res_48$CP, res_48$Genotype == "LOX3")
## W = 0.76411, p-value = 0.04
shapiro.test(subset(res_72$CP, res_72$Genotype=="EV")) # OK
##
##  Shapiro-Wilk normality test
##
## data:  subset(res_72$CP, res_72$Genotype == "EV")
## W = 0.85137, p-value = 0.1989
shapiro.test(subset(res_72$CP, res_72$Genotype=="ETR1")) # OK
##
##  Shapiro-Wilk normality test
##
## data:  subset(res_72$CP, res_72$Genotype == "ETR1")
## W = 0.93143, p-value = 0.6062
shapiro.test(subset(res_72$CP, res_72$Genotype=="LOX3")) # OK
##
##  Shapiro-Wilk normality test
##
## data:  subset(res_72$CP, res_72$Genotype == "LOX3")
## W = 0.82021, p-value = 0.1172
shapiro.test(subset(res_24$DCSs, res_24$Genotype=="EV")) # OK
##
##  Shapiro-Wilk normality test
##
## data:  subset(res_24$DCSs, res_24$Genotype == "EV")
## W = 0.95063, p-value = 0.7417
shapiro.test(subset(res_24$DCSs, res_24$Genotype=="ETR1")) # OK
##
##  Shapiro-Wilk normality test
##
## data:  subset(res_24$DCSs, res_24$Genotype == "ETR1")
## W = 0.86212, p-value = 0.236
shapiro.test(subset(res_24$DCSs, res_24$Genotype=="LOX3")) # OK
##
##  Shapiro-Wilk normality test
##
## data:  subset(res_24$DCSs, res_24$Genotype == "LOX3")
## W = 0.98517, p-value = 0.9602
shapiro.test(subset(res_48$DCSs, res_48$Genotype=="EV")) # OK
##
##  Shapiro-Wilk normality test
##
## data:  subset(res_48$DCSs, res_48$Genotype == "EV")
## W = 0.98376, p-value = 0.9537
shapiro.test(subset(res_48$DCSs, res_48$Genotype=="ETR1")) # OK
##
##  Shapiro-Wilk normality test
##
## data:  subset(res_48$DCSs, res_48$Genotype == "ETR1")
## W = 0.89154, p-value = 0.3649
shapiro.test(subset(res_48$DCSs, res_48$Genotype=="LOX3")) # OK
##
##  Shapiro-Wilk normality test
##
## data:  subset(res_48$DCSs, res_48$Genotype == "LOX3")
## W = 0.88589, p-value = 0.3369
shapiro.test(subset(res_72$DCSs, res_72$Genotype=="EV")) # OK
##
##  Shapiro-Wilk normality test
##
## data:  subset(res_72$DCSs, res_72$Genotype == "EV")
## W = 0.96205, p-value = 0.8222
shapiro.test(subset(res_72$DCSs, res_72$Genotype=="ETR1")) # OK
##
##  Shapiro-Wilk normality test
##
## data:  subset(res_72$DCSs, res_72$Genotype == "ETR1")
## W = 0.94593, p-value = 0.7081
shapiro.test(subset(res_72$DCSs, res_72$Genotype=="LOX3")) # OK
##
##  Shapiro-Wilk normality test
##
## data:  subset(res_72$DCSs, res_72$Genotype == "LOX3")
## W = 0.90891, p-value = 0.461
### Tests de variances

CP_24 = formula(res_24$CP~res_24$Genotype)
DCSs_24 = formula(res_24$DCSs~res_24$Genotype)
CP_48 = formula(res_48$CP~res_48$Genotype)
DCSs_48 = formula(res_48$DCSs~res_48$Genotype)
CP_72 = formula(res_72$CP~res_72$Genotype)
DCSs_72 = formula(res_72$DCSs~res_72$Genotype)


fligner.test(CP_24) # OK
##
##  Fligner-Killeen test of homogeneity of variances
##
## data:  res_24$CP by res_24$Genotype
## Fligner-Killeen:med chi-squared = 3.0143, df = 2, p-value = 0.2215
fligner.test(CP_48) # OK
##
##  Fligner-Killeen test of homogeneity of variances
##
## data:  res_48$CP by res_48$Genotype
## Fligner-Killeen:med chi-squared = 3.3728, df = 2, p-value = 0.1852
fligner.test(CP_72) # OK
##
##  Fligner-Killeen test of homogeneity of variances
##
## data:  res_72$CP by res_72$Genotype
## Fligner-Killeen:med chi-squared = 2.6345, df = 2, p-value = 0.2679
fligner.test(DCSs_24) # OK
##
##  Fligner-Killeen test of homogeneity of variances
##
## data:  res_24$DCSs by res_24$Genotype
## Fligner-Killeen:med chi-squared = 0.0022842, df = 2, p-value =
## 0.9989
fligner.test(DCSs_48) # OK
##
##  Fligner-Killeen test of homogeneity of variances
##
## data:  res_48$DCSs by res_48$Genotype
## Fligner-Killeen:med chi-squared = 2.914, df = 2, p-value = 0.2329
fligner.test(DCSs_72) # OK
##
##  Fligner-Killeen test of homogeneity of variances
##
## data:  res_72$DCSs by res_72$Genotype
## Fligner-Killeen:med chi-squared = 0.7843, df = 2, p-value = 0.6756
### ANOVAs

kruskal.test(CP_24)
##
##  Kruskal-Wallis rank sum test
##
## data:  res_24$CP by res_24$Genotype
## Kruskal-Wallis chi-squared = 7.5253, df = 2, p-value = 0.02322
pairwise.wilcox.test(res_24$CP, res_24$Genotype)
## Warning in wilcox.test.default(xi, xj, paired = paired, ...): cannot
## compute exact p-value with ties

## Warning in wilcox.test.default(xi, xj, paired = paired, ...): cannot
## compute exact p-value with ties

## Warning in wilcox.test.default(xi, xj, paired = paired, ...): cannot
## compute exact p-value with ties
##
##  Pairwise comparisons using Wilcoxon rank sum test
##
## data:  res_24$CP and res_24$Genotype
##
##      ETR1  EV
## EV   0.530 -
## LOX3 0.035 0.148
##
## P value adjustment method: holm
summary(aov(DCSs_24))
##                 Df Sum Sq Mean Sq F value   Pr(>F)
## res_24$Genotype  2  33701   16851   37.86 6.55e-06 ***
## Residuals       12   5340     445
## ---
## Signif. codes:  0 '***' 0.001 '**' 0.01 '*' 0.05 '.' 0.1 ' ' 1
TukeyHSD(aov(DCSs_24))
##   Tukey multiple comparisons of means
##     95% family-wise confidence level
##
## Fit: aov(formula = DCSs_24)
##
## $`res_24$Genotype`
##            diff       lwr       upr     p adj
## EV-ETR1    31.0 -4.595074  66.59507 0.0906916
## LOX3-ETR1 112.4 76.804926 147.99507 0.0000061
## LOX3-EV    81.4 45.804926 116.99507 0.0001459
kruskal.test(CP_48)
##
##  Kruskal-Wallis rank sum test
##
## data:  res_48$CP by res_48$Genotype
## Kruskal-Wallis chi-squared = 9.8504, df = 2, p-value = 0.007261
pairwise.wilcox.test(res_48$CP, res_48$Genotype)
## Warning in wilcox.test.default(xi, xj, paired = paired, ...): cannot
## compute exact p-value with ties

## Warning in wilcox.test.default(xi, xj, paired = paired, ...): cannot
## compute exact p-value with ties
##
##  Pairwise comparisons using Wilcoxon rank sum test
##
## data:  res_48$CP and res_48$Genotype
##
##      ETR1  EV
## EV   0.421 -
## LOX3 0.033 0.033
##
## P value adjustment method: holm
summary(aov(DCSs_48))
##                 Df Sum Sq Mean Sq F value Pr(>F)
## res_48$Genotype  2  12386    6193   3.941 0.0483 *
## Residuals       12  18858    1572
## ---
## Signif. codes:  0 '***' 0.001 '**' 0.01 '*' 0.05 '.' 0.1 ' ' 1
TukeyHSD(aov(DCSs_48))
##   Tukey multiple comparisons of means
##     95% family-wise confidence level
##
## Fit: aov(formula = DCSs_48)
##
## $`res_48$Genotype`
##           diff        lwr      upr     p adj
## EV-ETR1   64.8  -2.088402 131.6884 0.0578561
## LOX3-ETR1 56.2 -10.688402 123.0884 0.1041010
## LOX3-EV   -8.6 -75.488402  58.2884 0.9375344
summary(aov(CP_72))
##                 Df Sum Sq Mean Sq F value   Pr(>F)
## res_72$Genotype  2 795501  397750   16.32 0.000377 ***
## Residuals       12 292376   24365
## ---
## Signif. codes:  0 '***' 0.001 '**' 0.01 '*' 0.05 '.' 0.1 ' ' 1
TukeyHSD(aov(CP_72))
##   Tukey multiple comparisons of means
##     95% family-wise confidence level
##
## Fit: aov(formula = CP_72)
##
## $`res_72$Genotype`
##                 diff       lwr       upr     p adj
## EV-ETR1     37.77047 -225.6040  301.1449 0.9229891
## LOX3-ETR1 -468.53688 -731.9113 -205.1625 0.0012755
## LOX3-EV   -506.30735 -769.6818 -242.9329 0.0006748
summary(aov(DCSs_72))
##                 Df Sum Sq Mean Sq F value Pr(>F)
## res_72$Genotype  2    991   495.6   0.235  0.794
## Residuals       12  25263  2105.2
```

### Statistical analysis of polyamine levels by HPLC-UV

```
data = read.table("R_data.csv", header=T, sep=";", dec=".")

res_0 = subset(data, data$Time==0)
res_0.5 = subset(data, data$Time==0.5)
res_1 = subset(data, data$Time==1)
res_2 = subset(data, data$Time==2)

res_6l = subset(data, data$Time==6 & data$Local==T)
res_6l = subset(data, data$Time==24 & data$Local==T)
res_24l = subset(data, data$Time==24 & data$Local==T)
res_48l = subset(data, data$Time==48 & data$Local==T)
res_72l = subset(data, data$Time==72 & data$Local==T)

res_6s = subset(data, data$Time==6 & data$Local==F)
res_24s = subset(data, data$Time==24 & data$Local==F)
res_24s = subset(data, data$Time==24 & data$Local==F)
res_48s = subset(data, data$Time==48 & data$Local==F)
res_72s = subset(data, data$Time==72 & data$Local==F)

### Tests de normalite

shapiro.test(subset(res_0$Tyramine, res_0$Genotype=="EV"))
##
##  Shapiro-Wilk normality test
##
## data:  subset(res_0$Tyramine, res_0$Genotype == "EV")
## W = 0.9189, p-value = 0.5229
shapiro.test(subset(res_0$Tyramine, res_0$Genotype=="ETR1"))
##
##  Shapiro-Wilk normality test
##
## data:  subset(res_0$Tyramine, res_0$Genotype == "ETR1")
## W = 0.93442, p-value = 0.6268
shapiro.test(subset(res_0$Tyramine, res_0$Genotype=="LOX3"))
##
##  Shapiro-Wilk normality test
##
## data:  subset(res_0$Tyramine, res_0$Genotype == "LOX3")
## W = 0.90217, p-value = 0.422
shapiro.test(subset(res_0$Tyramine, res_0$Genotype=="MYB8"))
##
##  Shapiro-Wilk normality test
##
## data:  subset(res_0$Tyramine, res_0$Genotype == "MYB8")
## W = 0.79682, p-value = 0.07632
shapiro.test(subset(res_0$Putrescine, res_0$Genotype=="EV"))
##
##  Shapiro-Wilk normality test
##
## data:  subset(res_0$Putrescine, res_0$Genotype == "EV")
## W = 0.78465, p-value = 0.0604
shapiro.test(subset(res_0$Putrescine, res_0$Genotype=="ETR1"))
##
##  Shapiro-Wilk normality test
##
## data:  subset(res_0$Putrescine, res_0$Genotype == "ETR1")
## W = 0.95425, p-value = 0.7675
shapiro.test(subset(res_0$Putrescine, res_0$Genotype=="LOX3"))
##
##  Shapiro-Wilk normality test
##
## data:  subset(res_0$Putrescine, res_0$Genotype == "LOX3")
## W = 0.90172, p-value = 0.4195
shapiro.test(subset(res_0$Putrescine, res_0$Genotype=="MYB8"))
##
##  Shapiro-Wilk normality test
##
## data:  subset(res_0$Putrescine, res_0$Genotype == "MYB8")
## W = 0.84071, p-value = 0.1669
shapiro.test(subset(res_0$Spermidine, res_0$Genotype=="EV"))
##
##  Shapiro-Wilk normality test
##
## data:  subset(res_0$Spermidine, res_0$Genotype == "EV")
## W = 0.97355, p-value = 0.8975
shapiro.test(subset(res_0$Spermidine, res_0$Genotype=="ETR1"))
##
##  Shapiro-Wilk normality test
##
## data:  subset(res_0$Spermidine, res_0$Genotype == "ETR1")
## W = 0.79388, p-value = 0.07218
shapiro.test(subset(res_0$Spermidine, res_0$Genotype=="LOX3"))
##
##  Shapiro-Wilk normality test
##
## data:  subset(res_0$Spermidine, res_0$Genotype == "LOX3")
## W = 0.89457, p-value = 0.3806
shapiro.test(subset(res_0$Spermidine, res_0$Genotype=="MYB8")) # NO
##
##  Shapiro-Wilk normality test
##
## data:  subset(res_0$Spermidine, res_0$Genotype == "MYB8")
## W = 0.75302, p-value = 0.03173
shapiro.test(subset(res_0.5$Tyramine, res_0.5$Genotype=="EV"))
##
##  Shapiro-Wilk normality test
##
## data:  subset(res_0.5$Tyramine, res_0.5$Genotype == "EV")
## W = 0.96017, p-value = 0.8092
shapiro.test(subset(res_0.5$Tyramine, res_0.5$Genotype=="ETR1"))
##
##  Shapiro-Wilk normality test
##
## data:  subset(res_0.5$Tyramine, res_0.5$Genotype == "ETR1")
## W = 0.9493, p-value = 0.7322
shapiro.test(subset(res_0.5$Tyramine, res_0.5$Genotype=="LOX3"))
##
##  Shapiro-Wilk normality test
##
## data:  subset(res_0.5$Tyramine, res_0.5$Genotype == "LOX3")
## W = 0.83026, p-value = 0.1397
shapiro.test(subset(res_0.5$Tyramine, res_0.5$Genotype=="MYB8")) # NO
##
##  Shapiro-Wilk normality test
##
## data:  subset(res_0.5$Tyramine, res_0.5$Genotype == "MYB8")
## W = 0.71001, p-value = 0.01215
shapiro.test(subset(res_0.5$Putrescine, res_0.5$Genotype=="EV"))
##
##  Shapiro-Wilk normality test
##
## data:  subset(res_0.5$Putrescine, res_0.5$Genotype == "EV")
## W = 0.91993, p-value = 0.5295
shapiro.test(subset(res_0.5$Putrescine, res_0.5$Genotype=="ETR1"))
##
##  Shapiro-Wilk normality test
##
## data:  subset(res_0.5$Putrescine, res_0.5$Genotype == "ETR1")
## W = 0.88966, p-value = 0.3554
shapiro.test(subset(res_0.5$Putrescine, res_0.5$Genotype=="LOX3"))
##
##  Shapiro-Wilk normality test
##
## data:  subset(res_0.5$Putrescine, res_0.5$Genotype == "LOX3")
## W = 0.90709, p-value = 0.4503
shapiro.test(subset(res_0.5$Putrescine, res_0.5$Genotype=="MYB8")) # NO
##
##  Shapiro-Wilk normality test
##
## data:  subset(res_0.5$Putrescine, res_0.5$Genotype == "MYB8")
## W = 0.7211, p-value = 0.01572
shapiro.test(subset(res_0.5$Spermidine, res_0.5$Genotype=="EV"))
##
##  Shapiro-Wilk normality test
##
## data:  subset(res_0.5$Spermidine, res_0.5$Genotype == "EV")
## W = 0.94641, p-value = 0.7115
shapiro.test(subset(res_0.5$Spermidine, res_0.5$Genotype=="ETR1"))
##
##  Shapiro-Wilk normality test
##
## data:  subset(res_0.5$Spermidine, res_0.5$Genotype == "ETR1")
## W = 0.88315, p-value = 0.3238
shapiro.test(subset(res_0.5$Spermidine, res_0.5$Genotype=="LOX3"))
##
##  Shapiro-Wilk normality test
##
## data:  subset(res_0.5$Spermidine, res_0.5$Genotype == "LOX3")
## W = 0.97368, p-value = 0.8983
shapiro.test(subset(res_0.5$Spermidine, res_0.5$Genotype=="MYB8"))
##
##  Shapiro-Wilk normality test
##
## data:  subset(res_0.5$Spermidine, res_0.5$Genotype == "MYB8")
## W = 0.92689, p-value = 0.5753
shapiro.test(subset(res_1$Tyramine, res_1$Genotype=="EV"))
##
##  Shapiro-Wilk normality test
##
## data:  subset(res_1$Tyramine, res_1$Genotype == "EV")
## W = 0.9718, p-value = 0.8867
shapiro.test(subset(res_1$Tyramine, res_1$Genotype=="ETR1"))
##
##  Shapiro-Wilk normality test
##
## data:  subset(res_1$Tyramine, res_1$Genotype == "ETR1")
## W = 0.92419, p-value = 0.5573
shapiro.test(subset(res_1$Tyramine, res_1$Genotype=="LOX3"))
##
##  Shapiro-Wilk normality test
##
## data:  subset(res_1$Tyramine, res_1$Genotype == "LOX3")
## W = 0.82557, p-value = 0.1288
shapiro.test(subset(res_1$Tyramine, res_1$Genotype=="MYB8")) # NO
##
##  Shapiro-Wilk normality test
##
## data:  subset(res_1$Tyramine, res_1$Genotype == "MYB8")
## W = 0.75918, p-value = 0.03612
shapiro.test(subset(res_1$Putrescine, res_1$Genotype=="EV")) # NO
##
##  Shapiro-Wilk normality test
##
## data:  subset(res_1$Putrescine, res_1$Genotype == "EV")
## W = 0.70882, p-value = 0.01182
shapiro.test(subset(res_1$Putrescine, res_1$Genotype=="ETR1"))
##
##  Shapiro-Wilk normality test
##
## data:  subset(res_1$Putrescine, res_1$Genotype == "ETR1")
## W = 0.83027, p-value = 0.1398
shapiro.test(subset(res_1$Putrescine, res_1$Genotype=="LOX3"))
##
##  Shapiro-Wilk normality test
##
## data:  subset(res_1$Putrescine, res_1$Genotype == "LOX3")
## W = 0.80919, p-value = 0.09609
shapiro.test(subset(res_1$Putrescine, res_1$Genotype=="MYB8"))
##
##  Shapiro-Wilk normality test
##
## data:  subset(res_1$Putrescine, res_1$Genotype == "MYB8")
## W = 0.90109, p-value = 0.416
shapiro.test(subset(res_1$Spermidine, res_1$Genotype=="EV"))
##
##  Shapiro-Wilk normality test
##
## data:  subset(res_1$Spermidine, res_1$Genotype == "EV")
## W = 0.88584, p-value = 0.3366
shapiro.test(subset(res_1$Spermidine, res_1$Genotype=="ETR1"))
##
##  Shapiro-Wilk normality test
##
## data:  subset(res_1$Spermidine, res_1$Genotype == "ETR1")
## W = 0.9974, p-value = 0.9981
shapiro.test(subset(res_1$Spermidine, res_1$Genotype=="LOX3"))
##
##  Shapiro-Wilk normality test
##
## data:  subset(res_1$Spermidine, res_1$Genotype == "LOX3")
## W = 0.94812, p-value = 0.7238
shapiro.test(subset(res_1$Spermidine, res_1$Genotype=="MYB8"))
##
##  Shapiro-Wilk normality test
##
## data:  subset(res_1$Spermidine, res_1$Genotype == "MYB8")
## W = 0.95602, p-value = 0.78
shapiro.test(subset(res_2$Tyramine, res_2$Genotype=="EV"))
##
##  Shapiro-Wilk normality test
##
## data:  subset(res_2$Tyramine, res_2$Genotype == "EV")
## W = 0.89992, p-value = 0.4094
shapiro.test(subset(res_2$Tyramine, res_2$Genotype=="ETR1"))
##
##  Shapiro-Wilk normality test
##
## data:  subset(res_2$Tyramine, res_2$Genotype == "ETR1")
## W = 0.91458, p-value = 0.4956
shapiro.test(subset(res_2$Tyramine, res_2$Genotype=="LOX3"))
##
##  Shapiro-Wilk normality test
##
## data:  subset(res_2$Tyramine, res_2$Genotype == "LOX3")
## W = 0.94101, p-value = 0.6731
shapiro.test(subset(res_2$Tyramine, res_2$Genotype=="MYB8"))
##
##  Shapiro-Wilk normality test
##
## data:  subset(res_2$Tyramine, res_2$Genotype == "MYB8")
## W = 0.9721, p-value = 0.8886
shapiro.test(subset(res_2$Putrescine, res_2$Genotype=="EV"))
##
##  Shapiro-Wilk normality test
##
## data:  subset(res_2$Putrescine, res_2$Genotype == "EV")
## W = 0.81557, p-value = 0.1079
shapiro.test(subset(res_2$Putrescine, res_2$Genotype=="ETR1"))
##
##  Shapiro-Wilk normality test
##
## data:  subset(res_2$Putrescine, res_2$Genotype == "ETR1")
## W = 0.971, p-value = 0.8817
shapiro.test(subset(res_2$Putrescine, res_2$Genotype=="LOX3"))
##
##  Shapiro-Wilk normality test
##
## data:  subset(res_2$Putrescine, res_2$Genotype == "LOX3")
## W = 0.82165, p-value = 0.1202
shapiro.test(subset(res_2$Putrescine, res_2$Genotype=="MYB8"))
##
##  Shapiro-Wilk normality test
##
## data:  subset(res_2$Putrescine, res_2$Genotype == "MYB8")
## W = 0.95884, p-value = 0.7999
shapiro.test(subset(res_2$Spermidine, res_2$Genotype=="EV"))
##
##  Shapiro-Wilk normality test
##
## data:  subset(res_2$Spermidine, res_2$Genotype == "EV")
## W = 0.83212, p-value = 0.1443
shapiro.test(subset(res_2$Spermidine, res_2$Genotype=="ETR1"))
##
##  Shapiro-Wilk normality test
##
## data:  subset(res_2$Spermidine, res_2$Genotype == "ETR1")
## W = 0.82885, p-value = 0.1364
shapiro.test(subset(res_2$Spermidine, res_2$Genotype=="LOX3"))
##
##  Shapiro-Wilk normality test
##
## data:  subset(res_2$Spermidine, res_2$Genotype == "LOX3")
## W = 0.95625, p-value = 0.7816
shapiro.test(subset(res_2$Spermidine, res_2$Genotype=="MYB8"))
##
##  Shapiro-Wilk normality test
##
## data:  subset(res_2$Spermidine, res_2$Genotype == "MYB8")
## W = 0.94077, p-value = 0.6714
shapiro.test(subset(res_6l$Tyramine, res_6l$Genotype=="EV"))
##
##  Shapiro-Wilk normality test
##
## data:  subset(res_6l$Tyramine, res_6l$Genotype == "EV")
## W = 0.9888, p-value = 0.9753
shapiro.test(subset(res_6l$Tyramine, res_6l$Genotype=="ETR1"))
##
##  Shapiro-Wilk normality test
##
## data:  subset(res_6l$Tyramine, res_6l$Genotype == "ETR1")
## W = 0.89361, p-value = 0.3756
shapiro.test(subset(res_6l$Tyramine, res_6l$Genotype=="LOX3"))
##
##  Shapiro-Wilk normality test
##
## data:  subset(res_6l$Tyramine, res_6l$Genotype == "LOX3")
## W = 0.89733, p-value = 0.3953
shapiro.test(subset(res_6l$Tyramine, res_6l$Genotype=="MYB8"))
##
##  Shapiro-Wilk normality test
##
## data:  subset(res_6l$Tyramine, res_6l$Genotype == "MYB8")
## W = 0.96172, p-value = 0.8199
shapiro.test(subset(res_6l$Putrescine, res_6l$Genotype=="EV"))
##
##  Shapiro-Wilk normality test
##
## data:  subset(res_6l$Putrescine, res_6l$Genotype == "EV")
## W = 0.85868, p-value = 0.2236
shapiro.test(subset(res_6l$Putrescine, res_6l$Genotype=="ETR1"))
##
##  Shapiro-Wilk normality test
##
## data:  subset(res_6l$Putrescine, res_6l$Genotype == "ETR1")
## W = 0.904, p-value = 0.4324
shapiro.test(subset(res_6l$Putrescine, res_6l$Genotype=="LOX3"))
##
##  Shapiro-Wilk normality test
##
## data:  subset(res_6l$Putrescine, res_6l$Genotype == "LOX3")
## W = 0.99149, p-value = 0.9846
shapiro.test(subset(res_6l$Putrescine, res_6l$Genotype=="MYB8"))
##
##  Shapiro-Wilk normality test
##
## data:  subset(res_6l$Putrescine, res_6l$Genotype == "MYB8")
## W = 0.87864, p-value = 0.3032
shapiro.test(subset(res_6l$Spermidine, res_6l$Genotype=="EV"))
##
##  Shapiro-Wilk normality test
##
## data:  subset(res_6l$Spermidine, res_6l$Genotype == "EV")
## W = 0.97756, p-value = 0.9212
shapiro.test(subset(res_6l$Spermidine, res_6l$Genotype=="ETR1")) # NO
##
##  Shapiro-Wilk normality test
##
## data:  subset(res_6l$Spermidine, res_6l$Genotype == "ETR1")
## W = 0.77004, p-value = 0.04516
shapiro.test(subset(res_6l$Spermidine, res_6l$Genotype=="LOX3"))
##
##  Shapiro-Wilk normality test
##
## data:  subset(res_6l$Spermidine, res_6l$Genotype == "LOX3")
## W = 0.88249, p-value = 0.3208
shapiro.test(subset(res_6l$Spermidine, res_6l$Genotype=="MYB8")) # NO
##
##  Shapiro-Wilk normality test
##
## data:  subset(res_6l$Spermidine, res_6l$Genotype == "MYB8")
## W = 0.73252, p-value = 0.02034
shapiro.test(subset(res_24l$Tyramine, res_24l$Genotype=="EV"))
##
##  Shapiro-Wilk normality test
##
## data:  subset(res_24l$Tyramine, res_24l$Genotype == "EV")
## W = 0.9888, p-value = 0.9753
shapiro.test(subset(res_24l$Tyramine, res_24l$Genotype=="ETR1"))
##
##  Shapiro-Wilk normality test
##
## data:  subset(res_24l$Tyramine, res_24l$Genotype == "ETR1")
## W = 0.89361, p-value = 0.3756
shapiro.test(subset(res_24l$Tyramine, res_24l$Genotype=="LOX3"))
##
##  Shapiro-Wilk normality test
##
## data:  subset(res_24l$Tyramine, res_24l$Genotype == "LOX3")
## W = 0.89733, p-value = 0.3953
shapiro.test(subset(res_24l$Tyramine, res_24l$Genotype=="MYB8"))
##
##  Shapiro-Wilk normality test
##
## data:  subset(res_24l$Tyramine, res_24l$Genotype == "MYB8")
## W = 0.96172, p-value = 0.8199
shapiro.test(subset(res_24l$Putrescine, res_24l$Genotype=="EV"))
##
##  Shapiro-Wilk normality test
##
## data:  subset(res_24l$Putrescine, res_24l$Genotype == "EV")
## W = 0.85868, p-value = 0.2236
shapiro.test(subset(res_24l$Putrescine, res_24l$Genotype=="ETR1"))
##
##  Shapiro-Wilk normality test
##
## data:  subset(res_24l$Putrescine, res_24l$Genotype == "ETR1")
## W = 0.904, p-value = 0.4324
shapiro.test(subset(res_24l$Putrescine, res_24l$Genotype=="LOX3"))
##
##  Shapiro-Wilk normality test
##
## data:  subset(res_24l$Putrescine, res_24l$Genotype == "LOX3")
## W = 0.99149, p-value = 0.9846
shapiro.test(subset(res_24l$Putrescine, res_24l$Genotype=="MYB8"))
##
##  Shapiro-Wilk normality test
##
## data:  subset(res_24l$Putrescine, res_24l$Genotype == "MYB8")
## W = 0.87864, p-value = 0.3032
shapiro.test(subset(res_24l$Spermidine, res_24l$Genotype=="EV"))
##
##  Shapiro-Wilk normality test
##
## data:  subset(res_24l$Spermidine, res_24l$Genotype == "EV")
## W = 0.97756, p-value = 0.9212
shapiro.test(subset(res_24l$Spermidine, res_24l$Genotype=="ETR1"))
##
##  Shapiro-Wilk normality test
##
## data:  subset(res_24l$Spermidine, res_24l$Genotype == "ETR1")
## W = 0.77004, p-value = 0.04516
shapiro.test(subset(res_24l$Spermidine, res_24l$Genotype=="LOX3"))
##
##  Shapiro-Wilk normality test
##
## data:  subset(res_24l$Spermidine, res_24l$Genotype == "LOX3")
## W = 0.88249, p-value = 0.3208
shapiro.test(subset(res_24l$Spermidine, res_24l$Genotype=="MYB8"))
##
##  Shapiro-Wilk normality test
##
## data:  subset(res_24l$Spermidine, res_24l$Genotype == "MYB8")
## W = 0.73252, p-value = 0.02034
##
##  Shapiro-Wilk normality test
##
## data:  subset(res_24l$Spermine, res_24l$Genotype == "MYB8")
## W = 0.69182, p-value = 0.007849
shapiro.test(subset(res_48l$Tyramine, res_48l$Genotype=="EV"))
##
##  Shapiro-Wilk normality test
##
## data:  subset(res_48l$Tyramine, res_48l$Genotype == "EV")
## W = 0.86863, p-value = 0.2609
shapiro.test(subset(res_48l$Tyramine, res_48l$Genotype=="ETR1"))
##
##  Shapiro-Wilk normality test
##
## data:  subset(res_48l$Tyramine, res_48l$Genotype == "ETR1")
## W = 0.85569, p-value = 0.2132
shapiro.test(subset(res_48l$Tyramine, res_48l$Genotype=="LOX3"))
##
##  Shapiro-Wilk normality test
##
## data:  subset(res_48l$Tyramine, res_48l$Genotype == "LOX3")
## W = 0.92696, p-value = 0.5758
shapiro.test(subset(res_48l$Tyramine, res_48l$Genotype=="MYB8")) # NO
##
##  Shapiro-Wilk normality test
##
## data:  subset(res_48l$Tyramine, res_48l$Genotype == "MYB8")
## W = 0.67058, p-value = 0.004596
shapiro.test(subset(res_48l$Putrescine, res_48l$Genotype=="EV"))
##
##  Shapiro-Wilk normality test
##
## data:  subset(res_48l$Putrescine, res_48l$Genotype == "EV")
## W = 0.9314, p-value = 0.606
shapiro.test(subset(res_48l$Putrescine, res_48l$Genotype=="ETR1"))
##
##  Shapiro-Wilk normality test
##
## data:  subset(res_48l$Putrescine, res_48l$Genotype == "ETR1")
## W = 0.89883, p-value = 0.4035
shapiro.test(subset(res_48l$Putrescine, res_48l$Genotype=="LOX3"))
##
##  Shapiro-Wilk normality test
##
## data:  subset(res_48l$Putrescine, res_48l$Genotype == "LOX3")
## W = 0.84273, p-value = 0.1726
shapiro.test(subset(res_48l$Putrescine, res_48l$Genotype=="MYB8"))
##
##  Shapiro-Wilk normality test
##
## data:  subset(res_48l$Putrescine, res_48l$Genotype == "MYB8")
## W = 0.9593, p-value = 0.8031
shapiro.test(subset(res_48l$Spermidine, res_48l$Genotype=="EV"))
##
##  Shapiro-Wilk normality test
##
## data:  subset(res_48l$Spermidine, res_48l$Genotype == "EV")
## W = 0.91765, p-value = 0.5149
shapiro.test(subset(res_48l$Spermidine, res_48l$Genotype=="ETR1"))
##
##  Shapiro-Wilk normality test
##
## data:  subset(res_48l$Spermidine, res_48l$Genotype == "ETR1")
## W = 0.85615, p-value = 0.2148
shapiro.test(subset(res_48l$Spermidine, res_48l$Genotype=="LOX3"))
##
##  Shapiro-Wilk normality test
##
## data:  subset(res_48l$Spermidine, res_48l$Genotype == "LOX3")
## W = 0.81628, p-value = 0.1093
shapiro.test(subset(res_48l$Spermidine, res_48l$Genotype=="MYB8"))
##
##  Shapiro-Wilk normality test
##
## data:  subset(res_48l$Spermidine, res_48l$Genotype == "MYB8")
## W = 0.95344, p-value = 0.7617
shapiro.test(subset(res_72l$Tyramine, res_72l$Genotype=="EV"))
##
##  Shapiro-Wilk normality test
##
## data:  subset(res_72l$Tyramine, res_72l$Genotype == "EV")
## W = 0.96108, p-value = 0.8155
shapiro.test(subset(res_72l$Tyramine, res_72l$Genotype=="ETR1"))
##
##  Shapiro-Wilk normality test
##
## data:  subset(res_72l$Tyramine, res_72l$Genotype == "ETR1")
## W = 0.93125, p-value = 0.6049
shapiro.test(subset(res_72l$Tyramine, res_72l$Genotype=="LOX3"))
##
##  Shapiro-Wilk normality test
##
## data:  subset(res_72l$Tyramine, res_72l$Genotype == "LOX3")
## W = 0.93285, p-value = 0.6159
shapiro.test(subset(res_72l$Tyramine, res_72l$Genotype=="MYB8"))
##
##  Shapiro-Wilk normality test
##
## data:  subset(res_72l$Tyramine, res_72l$Genotype == "MYB8")
## W = 0.81092, p-value = 0.09916
shapiro.test(subset(res_72l$Putrescine, res_72l$Genotype=="EV"))
##
##  Shapiro-Wilk normality test
##
## data:  subset(res_72l$Putrescine, res_72l$Genotype == "EV")
## W = 0.8723, p-value = 0.2758
shapiro.test(subset(res_72l$Putrescine, res_72l$Genotype=="ETR1"))
##
##  Shapiro-Wilk normality test
##
## data:  subset(res_72l$Putrescine, res_72l$Genotype == "ETR1")
## W = 0.87118, p-value = 0.2712
shapiro.test(subset(res_72l$Putrescine, res_72l$Genotype=="LOX3"))
##
##  Shapiro-Wilk normality test
##
## data:  subset(res_72l$Putrescine, res_72l$Genotype == "LOX3")
## W = 0.97351, p-value = 0.8973
shapiro.test(subset(res_72l$Putrescine, res_72l$Genotype=="MYB8"))
##
##  Shapiro-Wilk normality test
##
## data:  subset(res_72l$Putrescine, res_72l$Genotype == "MYB8")
## W = 0.79386, p-value = 0.07215
shapiro.test(subset(res_72l$Spermidine, res_72l$Genotype=="EV"))
##
##  Shapiro-Wilk normality test
##
## data:  subset(res_72l$Spermidine, res_72l$Genotype == "EV")
## W = 0.90664, p-value = 0.4477
shapiro.test(subset(res_72l$Spermidine, res_72l$Genotype=="ETR1"))
##
##  Shapiro-Wilk normality test
##
## data:  subset(res_72l$Spermidine, res_72l$Genotype == "ETR1")
## W = 0.8649, p-value = 0.2464
shapiro.test(subset(res_72l$Spermidine, res_72l$Genotype=="LOX3"))
##
##  Shapiro-Wilk normality test
##
## data:  subset(res_72l$Spermidine, res_72l$Genotype == "LOX3")
## W = 0.92859, p-value = 0.5868
shapiro.test(subset(res_72l$Spermidine, res_72l$Genotype=="MYB8"))
##
##  Shapiro-Wilk normality test
##
## data:  subset(res_72l$Spermidine, res_72l$Genotype == "MYB8")
## W = 0.81512, p-value = 0.107
shapiro.test(subset(res_6s$Tyramine, res_6s$Genotype=="EV")) # NO
##
##  Shapiro-Wilk normality test
##
## data:  subset(res_6s$Tyramine, res_6s$Genotype == "EV")
## W = 0.76955, p-value = 0.04471
shapiro.test(subset(res_6s$Tyramine, res_6s$Genotype=="ETR1"))
##
##  Shapiro-Wilk normality test
##
## data:  subset(res_6s$Tyramine, res_6s$Genotype == "ETR1")
## W = 0.9204, p-value = 0.5325
shapiro.test(subset(res_6s$Tyramine, res_6s$Genotype=="LOX3"))
##
##  Shapiro-Wilk normality test
##
## data:  subset(res_6s$Tyramine, res_6s$Genotype == "LOX3")
## W = 0.95883, p-value = 0.7998
shapiro.test(subset(res_6s$Tyramine, res_6s$Genotype=="MYB8")) # NO
##
##  Shapiro-Wilk normality test
##
## data:  subset(res_6s$Tyramine, res_6s$Genotype == "MYB8")
## W = 0.73604, p-value = 0.02199
shapiro.test(subset(res_6s$Putrescine, res_6s$Genotype=="EV"))
##
##  Shapiro-Wilk normality test
##
## data:  subset(res_6s$Putrescine, res_6s$Genotype == "EV")
## W = 0.90644, p-value = 0.4465
shapiro.test(subset(res_6s$Putrescine, res_6s$Genotype=="ETR1"))
##
##  Shapiro-Wilk normality test
##
## data:  subset(res_6s$Putrescine, res_6s$Genotype == "ETR1")
## W = 0.91497, p-value = 0.498
shapiro.test(subset(res_6s$Putrescine, res_6s$Genotype=="LOX3"))
##
##  Shapiro-Wilk normality test
##
## data:  subset(res_6s$Putrescine, res_6s$Genotype == "LOX3")
## W = 0.89428, p-value = 0.3791
shapiro.test(subset(res_6s$Putrescine, res_6s$Genotype=="MYB8"))
##
##  Shapiro-Wilk normality test
##
## data:  subset(res_6s$Putrescine, res_6s$Genotype == "MYB8")
## W = 0.90872, p-value = 0.46
shapiro.test(subset(res_6s$Spermidine, res_6s$Genotype=="EV"))
##
##  Shapiro-Wilk normality test
##
## data:  subset(res_6s$Spermidine, res_6s$Genotype == "EV")
## W = 0.98462, p-value = 0.9577
shapiro.test(subset(res_6s$Spermidine, res_6s$Genotype=="ETR1"))
##
##  Shapiro-Wilk normality test
##
## data:  subset(res_6s$Spermidine, res_6s$Genotype == "ETR1")
## W = 0.93824, p-value = 0.6536
shapiro.test(subset(res_6s$Spermidine, res_6s$Genotype=="LOX3"))
##
##  Shapiro-Wilk normality test
##
## data:  subset(res_6s$Spermidine, res_6s$Genotype == "LOX3")
## W = 0.91644, p-value = 0.5072
shapiro.test(subset(res_6s$Spermidine, res_6s$Genotype=="MYB8"))
##
##  Shapiro-Wilk normality test
##
## data:  subset(res_6s$Spermidine, res_6s$Genotype == "MYB8")
## W = 0.8194, p-value = 0.1155
shapiro.test(subset(res_24s$Tyramine, res_24s$Genotype=="EV"))
##
##  Shapiro-Wilk normality test
##
## data:  subset(res_24s$Tyramine, res_24s$Genotype == "EV")
## W = 0.94072, p-value = 0.6711
shapiro.test(subset(res_24s$Tyramine, res_24s$Genotype=="ETR1"))
##
##  Shapiro-Wilk normality test
##
## data:  subset(res_24s$Tyramine, res_24s$Genotype == "ETR1")
## W = 0.83233, p-value = 0.1448
shapiro.test(subset(res_24s$Tyramine, res_24s$Genotype=="LOX3"))
##
##  Shapiro-Wilk normality test
##
## data:  subset(res_24s$Tyramine, res_24s$Genotype == "LOX3")
## W = 0.89098, p-value = 0.3621
shapiro.test(subset(res_24s$Tyramine, res_24s$Genotype=="MYB8"))
##
##  Shapiro-Wilk normality test
##
## data:  subset(res_24s$Tyramine, res_24s$Genotype == "MYB8")
## W = 0.87303, p-value = 0.2789
shapiro.test(subset(res_24s$Putrescine, res_24s$Genotype=="EV"))
##
##  Shapiro-Wilk normality test
##
## data:  subset(res_24s$Putrescine, res_24s$Genotype == "EV")
## W = 0.97563, p-value = 0.91
shapiro.test(subset(res_24s$Putrescine, res_24s$Genotype=="ETR1"))
##
##  Shapiro-Wilk normality test
##
## data:  subset(res_24s$Putrescine, res_24s$Genotype == "ETR1")
## W = 0.96881, p-value = 0.8675
shapiro.test(subset(res_24s$Putrescine, res_24s$Genotype=="LOX3"))
##
##  Shapiro-Wilk normality test
##
## data:  subset(res_24s$Putrescine, res_24s$Genotype == "LOX3")
## W = 0.97744, p-value = 0.9205
shapiro.test(subset(res_24s$Putrescine, res_24s$Genotype=="MYB8"))
##
##  Shapiro-Wilk normality test
##
## data:  subset(res_24s$Putrescine, res_24s$Genotype == "MYB8")
## W = 0.96888, p-value = 0.868
shapiro.test(subset(res_24s$Spermidine, res_24s$Genotype=="EV"))
##
##  Shapiro-Wilk normality test
##
## data:  subset(res_24s$Spermidine, res_24s$Genotype == "EV")
## W = 0.96597, p-value = 0.8488
shapiro.test(subset(res_24s$Spermidine, res_24s$Genotype=="ETR1"))
##
##  Shapiro-Wilk normality test
##
## data:  subset(res_24s$Spermidine, res_24s$Genotype == "ETR1")
## W = 0.91094, p-value = 0.4732
shapiro.test(subset(res_24s$Spermidine, res_24s$Genotype=="LOX3"))
##
##  Shapiro-Wilk normality test
##
## data:  subset(res_24s$Spermidine, res_24s$Genotype == "LOX3")
## W = 0.98645, p-value = 0.9658
shapiro.test(subset(res_24s$Spermidine, res_24s$Genotype=="MYB8"))
##
##  Shapiro-Wilk normality test
##
## data:  subset(res_24s$Spermidine, res_24s$Genotype == "MYB8")
## W = 0.98931, p-value = 0.9772
shapiro.test(subset(res_48s$Tyramine, res_48s$Genotype=="EV"))
##
##  Shapiro-Wilk normality test
##
## data:  subset(res_48s$Tyramine, res_48s$Genotype == "EV")
## W = 0.96575, p-value = 0.8474
shapiro.test(subset(res_48s$Tyramine, res_48s$Genotype=="ETR1"))
##
##  Shapiro-Wilk normality test
##
## data:  subset(res_48s$Tyramine, res_48s$Genotype == "ETR1")
## W = 0.94047, p-value = 0.6692
shapiro.test(subset(res_48s$Tyramine, res_48s$Genotype=="LOX3"))
##
##  Shapiro-Wilk normality test
##
## data:  subset(res_48s$Tyramine, res_48s$Genotype == "LOX3")
## W = 0.98524, p-value = 0.9606
shapiro.test(subset(res_48s$Tyramine, res_48s$Genotype=="MYB8"))
##
##  Shapiro-Wilk normality test
##
## data:  subset(res_48s$Tyramine, res_48s$Genotype == "MYB8")
## W = 0.83077, p-value = 0.141
shapiro.test(subset(res_48s$Putrescine, res_48s$Genotype=="EV"))
##
##  Shapiro-Wilk normality test
##
## data:  subset(res_48s$Putrescine, res_48s$Genotype == "EV")
## W = 0.94733, p-value = 0.7181
shapiro.test(subset(res_48s$Putrescine, res_48s$Genotype=="ETR1"))
##
##  Shapiro-Wilk normality test
##
## data:  subset(res_48s$Putrescine, res_48s$Genotype == "ETR1")
## W = 0.83145, p-value = 0.1426
shapiro.test(subset(res_48s$Putrescine, res_48s$Genotype=="LOX3"))
##
##  Shapiro-Wilk normality test
##
## data:  subset(res_48s$Putrescine, res_48s$Genotype == "LOX3")
## W = 0.94131, p-value = 0.6752
shapiro.test(subset(res_48s$Putrescine, res_48s$Genotype=="MYB8"))
##
##  Shapiro-Wilk normality test
##
## data:  subset(res_48s$Putrescine, res_48s$Genotype == "MYB8")
## W = 0.97092, p-value = 0.8811
shapiro.test(subset(res_48s$Spermidine, res_48s$Genotype=="EV"))
##
##  Shapiro-Wilk normality test
##
## data:  subset(res_48s$Spermidine, res_48s$Genotype == "EV")
## W = 0.95738, p-value = 0.7896
shapiro.test(subset(res_48s$Spermidine, res_48s$Genotype=="ETR1"))
##
##  Shapiro-Wilk normality test
##
## data:  subset(res_48s$Spermidine, res_48s$Genotype == "ETR1")
## W = 0.87069, p-value = 0.2692
shapiro.test(subset(res_48s$Spermidine, res_48s$Genotype=="LOX3"))
##
##  Shapiro-Wilk normality test
##
## data:  subset(res_48s$Spermidine, res_48s$Genotype == "LOX3")
## W = 0.89894, p-value = 0.404
shapiro.test(subset(res_48s$Spermidine, res_48s$Genotype=="MYB8"))
##
##  Shapiro-Wilk normality test
##
## data:  subset(res_48s$Spermidine, res_48s$Genotype == "MYB8")
## W = 0.80395, p-value = 0.08723
shapiro.test(subset(res_72s$Tyramine, res_72s$Genotype=="EV"))
##
##  Shapiro-Wilk normality test
##
## data:  subset(res_72s$Tyramine, res_72s$Genotype == "EV")
## W = 0.90796, p-value = 0.4554
shapiro.test(subset(res_72s$Tyramine, res_72s$Genotype=="ETR1")) # NO
##
##  Shapiro-Wilk normality test
##
## data:  subset(res_72s$Tyramine, res_72s$Genotype == "ETR1")
## W = 0.71816, p-value = 0.01469
shapiro.test(subset(res_72s$Tyramine, res_72s$Genotype=="LOX3"))
##
##  Shapiro-Wilk normality test
##
## data:  subset(res_72s$Tyramine, res_72s$Genotype == "LOX3")
## W = 0.85878, p-value = 0.2239
shapiro.test(subset(res_72s$Tyramine, res_72s$Genotype=="MYB8"))
##
##  Shapiro-Wilk normality test
##
## data:  subset(res_72s$Tyramine, res_72s$Genotype == "MYB8")
## W = 0.92624, p-value = 0.571
shapiro.test(subset(res_72s$Putrescine, res_72s$Genotype=="EV"))
##
##  Shapiro-Wilk normality test
##
## data:  subset(res_72s$Putrescine, res_72s$Genotype == "EV")
## W = 0.93634, p-value = 0.6402
shapiro.test(subset(res_72s$Putrescine, res_72s$Genotype=="ETR1"))
##
##  Shapiro-Wilk normality test
##
## data:  subset(res_72s$Putrescine, res_72s$Genotype == "ETR1")
## W = 0.9803, p-value = 0.9362
shapiro.test(subset(res_72s$Putrescine, res_72s$Genotype=="LOX3"))
##
##  Shapiro-Wilk normality test
##
## data:  subset(res_72s$Putrescine, res_72s$Genotype == "LOX3")
## W = 0.89815, p-value = 0.3997
shapiro.test(subset(res_72s$Putrescine, res_72s$Genotype=="MYB8"))
##
##  Shapiro-Wilk normality test
##
## data:  subset(res_72s$Putrescine, res_72s$Genotype == "MYB8")
## W = 0.89756, p-value = 0.3966
shapiro.test(subset(res_72s$Spermidine, res_72s$Genotype=="EV"))
##
##  Shapiro-Wilk normality test
##
## data:  subset(res_72s$Spermidine, res_72s$Genotype == "EV")
## W = 0.93192, p-value = 0.6095
shapiro.test(subset(res_72s$Spermidine, res_72s$Genotype=="ETR1"))
##
##  Shapiro-Wilk normality test
##
## data:  subset(res_72s$Spermidine, res_72s$Genotype == "ETR1")
## W = 0.78888, p-value = 0.06557
shapiro.test(subset(res_72s$Spermidine, res_72s$Genotype=="LOX3"))
##
##  Shapiro-Wilk normality test
##
## data:  subset(res_72s$Spermidine, res_72s$Genotype == "LOX3")
## W = 0.96126, p-value = 0.8167
shapiro.test(subset(res_72s$Spermidine, res_72s$Genotype=="MYB8"))
##
##  Shapiro-Wilk normality test
##
## data:  subset(res_72s$Spermidine, res_72s$Genotype == "MYB8")
## W = 0.92058, p-value = 0.5337

Tyramine_0 = formula(res_0$Tyramine~res_0$Genotype)
fligner.test(Tyramine_0)
##
##  Fligner-Killeen test of homogeneity of variances
##
## data:  res_0$Tyramine by res_0$Genotype
## Fligner-Killeen:med chi-squared = 2.6281, df = 3, p-value = 0.4526
Putrescine_0 = formula(res_0$Putrescine~res_0$Genotype)
fligner.test(Putrescine_0)
##
##  Fligner-Killeen test of homogeneity of variances
##
## data:  res_0$Putrescine by res_0$Genotype
## Fligner-Killeen:med chi-squared = 1.1534, df = 3, p-value = 0.7642
Spermidine_0 = formula(res_0$Spermidine~res_0$Genotype)
fligner.test(Spermidine_0)
##
##  Fligner-Killeen test of homogeneity of variances
##
## data:  res_0$Spermidine by res_0$Genotype
## Fligner-Killeen:med chi-squared = 1.5713, df = 3, p-value = 0.6659
Tyramine_0.5 = formula(res_0.5$Tyramine~res_0.5$Genotype)
fligner.test(Tyramine_0.5)
##
##  Fligner-Killeen test of homogeneity of variances
##
## data:  res_0.5$Tyramine by res_0.5$Genotype
## Fligner-Killeen:med chi-squared = 0.358, df = 3, p-value = 0.9488
Putrescine_0.5 = formula(res_0.5$Putrescine~res_0.5$Genotype)
fligner.test(Putrescine_0.5)
##
##  Fligner-Killeen test of homogeneity of variances
##
## data:  res_0.5$Putrescine by res_0.5$Genotype
## Fligner-Killeen:med chi-squared = 0.38269, df = 3, p-value =
## 0.9438
Spermidine_0.5 = formula(res_0.5$Spermidine~res_0.5$Genotype)
fligner.test(Spermidine_0.5)
##
##  Fligner-Killeen test of homogeneity of variances
##
## data:  res_0.5$Spermidine by res_0.5$Genotype
## Fligner-Killeen:med chi-squared = 1.6064, df = 3, p-value = 0.6579
Tyramine_1 = formula(res_1$Tyramine~res_1$Genotype)
fligner.test(Tyramine_1)
##
##  Fligner-Killeen test of homogeneity of variances
##
## data:  res_1$Tyramine by res_1$Genotype
## Fligner-Killeen:med chi-squared = 0.74762, df = 3, p-value = 0.862
Putrescine_1 = formula(res_1$Putrescine~res_1$Genotype)
fligner.test(Putrescine_1)
##
##  Fligner-Killeen test of homogeneity of variances
##
## data:  res_1$Putrescine by res_1$Genotype
## Fligner-Killeen:med chi-squared = 4.1223, df = 3, p-value = 0.2486
Spermidine_1 = formula(res_1$Spermidine~res_1$Genotype)
fligner.test(Spermidine_1)
##
##  Fligner-Killeen test of homogeneity of variances
##
## data:  res_1$Spermidine by res_1$Genotype
## Fligner-Killeen:med chi-squared = 3.4316, df = 3, p-value = 0.3297
Tyramine_2 = formula(res_2$Tyramine~res_2$Genotype)
fligner.test(Tyramine_2)
##
##  Fligner-Killeen test of homogeneity of variances
##
## data:  res_2$Tyramine by res_2$Genotype
## Fligner-Killeen:med chi-squared = 3.758, df = 3, p-value = 0.2888
Putrescine_2 = formula(res_2$Putrescine~res_2$Genotype)
fligner.test(Putrescine_2)
##
##  Fligner-Killeen test of homogeneity of variances
##
## data:  res_2$Putrescine by res_2$Genotype
## Fligner-Killeen:med chi-squared = 1.1904, df = 3, p-value = 0.7553
Spermidine_2 = formula(res_2$Spermidine~res_2$Genotype)
fligner.test(Spermidine_2)
##
##  Fligner-Killeen test of homogeneity of variances
##
## data:  res_2$Spermidine by res_2$Genotype
## Fligner-Killeen:med chi-squared = 2.941, df = 3, p-value = 0.4008
Tyramine_6l = formula(res_6l$Tyramine~res_6l$Genotype)
fligner.test(Tyramine_6l)
##
##  Fligner-Killeen test of homogeneity of variances
##
## data:  res_6l$Tyramine by res_6l$Genotype
## Fligner-Killeen:med chi-squared = 6.9937, df = 3, p-value = 0.0721
Putrescine_6l = formula(res_6l$Putrescine~res_6l$Genotype)
fligner.test(Putrescine_6l)
##
##  Fligner-Killeen test of homogeneity of variances
##
## data:  res_6l$Putrescine by res_6l$Genotype
## Fligner-Killeen:med chi-squared = 2.426, df = 3, p-value = 0.4888
Spermidine_6l = formula(res_6l$Spermidine~res_6l$Genotype)
fligner.test(Spermidine_6l)
##
##  Fligner-Killeen test of homogeneity of variances
##
## data:  res_6l$Spermidine by res_6l$Genotype
## Fligner-Killeen:med chi-squared = 1.0443, df = 3, p-value = 0.7905
Tyramine_24l = formula(res_24l$Tyramine~res_24l$Genotype)
fligner.test(Tyramine_24l)
##
##  Fligner-Killeen test of homogeneity of variances
##
## data:  res_24l$Tyramine by res_24l$Genotype
## Fligner-Killeen:med chi-squared = 6.9937, df = 3, p-value = 0.0721
Putrescine_24l = formula(res_24l$Putrescine~res_24l$Genotype)
fligner.test(Putrescine_24l)
##
##  Fligner-Killeen test of homogeneity of variances
##
## data:  res_24l$Putrescine by res_24l$Genotype
## Fligner-Killeen:med chi-squared = 2.426, df = 3, p-value = 0.4888
Spermidine_24l = formula(res_24l$Spermidine~res_24l$Genotype)
fligner.test(Spermidine_24l)
##
##  Fligner-Killeen test of homogeneity of variances
##
## data:  res_24l$Spermidine by res_24l$Genotype
## Fligner-Killeen:med chi-squared = 1.0443, df = 3, p-value = 0.7905
Tyramine_48l = formula(res_48l$Tyramine~res_48l$Genotype)
fligner.test(Tyramine_48l)
##
##  Fligner-Killeen test of homogeneity of variances
##
## data:  res_48l$Tyramine by res_48l$Genotype
## Fligner-Killeen:med chi-squared = 4.7932, df = 3, p-value = 0.1876
Putrescine_48l = formula(res_48l$Putrescine~res_48l$Genotype)
fligner.test(Putrescine_48l)
##
##  Fligner-Killeen test of homogeneity of variances
##
## data:  res_48l$Putrescine by res_48l$Genotype
## Fligner-Killeen:med chi-squared = 3.1232, df = 3, p-value = 0.373
Spermidine_48l = formula(res_48l$Spermidine~res_48l$Genotype)
fligner.test(Spermidine_48l)
##
##  Fligner-Killeen test of homogeneity of variances
##
## data:  res_48l$Spermidine by res_48l$Genotype
## Fligner-Killeen:med chi-squared = 0.1222, df = 3, p-value = 0.989
Tyramine_72l = formula(res_72l$Tyramine~res_72l$Genotype)
fligner.test(Tyramine_72l)
##
##  Fligner-Killeen test of homogeneity of variances
##
## data:  res_72l$Tyramine by res_72l$Genotype
## Fligner-Killeen:med chi-squared = 5.1143, df = 3, p-value = 0.1636
Putrescine_72l = formula(res_72l$Putrescine~res_72l$Genotype)
fligner.test(Putrescine_72l)
##
##  Fligner-Killeen test of homogeneity of variances
##
## data:  res_72l$Putrescine by res_72l$Genotype
## Fligner-Killeen:med chi-squared = 2.2089, df = 3, p-value = 0.5302
Spermidine_72l = formula(res_72l$Spermidine~res_72l$Genotype)
fligner.test(Spermidine_72l)
##
##  Fligner-Killeen test of homogeneity of variances
##
## data:  res_72l$Spermidine by res_72l$Genotype
## Fligner-Killeen:med chi-squared = 1.6964, df = 3, p-value = 0.6377
Tyramine_6s = formula(res_6s$Tyramine~res_6s$Genotype)
fligner.test(Tyramine_6s)
##
##  Fligner-Killeen test of homogeneity of variances
##
## data:  res_6s$Tyramine by res_6s$Genotype
## Fligner-Killeen:med chi-squared = 4.5841, df = 3, p-value = 0.2049
Putrescine_6s = formula(res_6s$Putrescine~res_6s$Genotype)
fligner.test(Putrescine_6s)
##
##  Fligner-Killeen test of homogeneity of variances
##
## data:  res_6s$Putrescine by res_6s$Genotype
## Fligner-Killeen:med chi-squared = 2.0528, df = 3, p-value = 0.5615
Spermidine_6s = formula(res_6s$Spermidine~res_6s$Genotype)
fligner.test(Spermidine_6s)
##
##  Fligner-Killeen test of homogeneity of variances
##
## data:  res_6s$Spermidine by res_6s$Genotype
## Fligner-Killeen:med chi-squared = 0.97066, df = 3, p-value =
## 0.8084
Tyramine_24s = formula(res_24s$Tyramine~res_24s$Genotype)
fligner.test(Tyramine_24s)
##
##  Fligner-Killeen test of homogeneity of variances
##
## data:  res_24s$Tyramine by res_24s$Genotype
## Fligner-Killeen:med chi-squared = 0.54358, df = 3, p-value =
## 0.9092
Putrescine_24s = formula(res_24s$Putrescine~res_24s$Genotype)
fligner.test(Putrescine_24s)
##
##  Fligner-Killeen test of homogeneity of variances
##
## data:  res_24s$Putrescine by res_24s$Genotype
## Fligner-Killeen:med chi-squared = 0.73426, df = 3, p-value =
## 0.8651
Spermidine_24s = formula(res_24s$Spermidine~res_24s$Genotype)
fligner.test(Spermidine_24s)
##
##  Fligner-Killeen test of homogeneity of variances
##
## data:  res_24s$Spermidine by res_24s$Genotype
## Fligner-Killeen:med chi-squared = 0.87083, df = 3, p-value =
## 0.8325
Tyramine_48s = formula(res_48s$Tyramine~res_48s$Genotype)
fligner.test(Tyramine_48s)
##
##  Fligner-Killeen test of homogeneity of variances
##
## data:  res_48s$Tyramine by res_48s$Genotype
## Fligner-Killeen:med chi-squared = 0.22961, df = 3, p-value =
## 0.9727
Putrescine_48s = formula(res_48s$Putrescine~res_48s$Genotype)
fligner.test(Putrescine_48s)
##
##  Fligner-Killeen test of homogeneity of variances
##
## data:  res_48s$Putrescine by res_48s$Genotype
## Fligner-Killeen:med chi-squared = 1.526, df = 3, p-value = 0.6763
Spermidine_48s = formula(res_48s$Spermidine~res_48s$Genotype)
fligner.test(Spermidine_48s)
##
##  Fligner-Killeen test of homogeneity of variances
##
## data:  res_48s$Spermidine by res_48s$Genotype
## Fligner-Killeen:med chi-squared = 2.2635, df = 3, p-value = 0.5196
Tyramine_72s = formula(res_72s$Tyramine~res_72s$Genotype)
fligner.test(Tyramine_72s)
##
##  Fligner-Killeen test of homogeneity of variances
##
## data:  res_72s$Tyramine by res_72s$Genotype
## Fligner-Killeen:med chi-squared = 3.1576, df = 3, p-value = 0.368
Putrescine_72s = formula(res_72s$Putrescine~res_72s$Genotype)
fligner.test(Putrescine_72s)
##
##  Fligner-Killeen test of homogeneity of variances
##
## data:  res_72s$Putrescine by res_72s$Genotype
## Fligner-Killeen:med chi-squared = 0.72235, df = 3, p-value =
## 0.8679
Spermidine_72s = formula(res_72s$Spermidine~res_72s$Genotype)
fligner.test(Spermidine_72s)
##
##  Fligner-Killeen test of homogeneity of variances
##
## data:  res_72s$Spermidine by res_72s$Genotype
## Fligner-Killeen:med chi-squared = 3.5223, df = 3, p-value = 0.3179
### ANOVA

summary(aov(Tyramine_0))
##                Df Sum Sq Mean Sq F value Pr(>F)
## res_0$Genotype  3  1.068  0.3561   2.722 0.0788 .
## Residuals      16  2.093  0.1308
## ---
## Signif. codes:  0 '***' 0.001 '**' 0.01 '*' 0.05 '.' 0.1 ' ' 1
summary(aov(Putrescine_0))
##                Df Sum Sq Mean Sq F value Pr(>F)
## res_0$Genotype  3  83.64   27.88   1.988  0.156
## Residuals      16 224.33   14.02
kruskal.test(Spermidine_0)
##
##  Kruskal-Wallis rank sum test
##
## data:  res_0$Spermidine by res_0$Genotype
## Kruskal-Wallis chi-squared = 2.4514, df = 3, p-value = 0.4841
kruskal.test(Spermine_0)
##
##  Kruskal-Wallis rank sum test
##
## data:  res_0$Spermine by res_0$Genotype
## Kruskal-Wallis chi-squared = 1.6763, df = 3, p-value = 0.6422
kruskal.test(Tyramine_0.5)
##
##  Kruskal-Wallis rank sum test
##
## data:  res_0.5$Tyramine by res_0.5$Genotype
## Kruskal-Wallis chi-squared = 5.64, df = 3, p-value = 0.1305
kruskal.test(Putrescine_0.5)
##
##  Kruskal-Wallis rank sum test
##
## data:  res_0.5$Putrescine by res_0.5$Genotype
## Kruskal-Wallis chi-squared = 7.6171, df = 3, p-value = 0.05462
summary(aov(Spermidine_0.5))
##                  Df Sum Sq Mean Sq F value Pr(>F)
## res_0.5$Genotype  3   5386    1796   1.068   0.39
## Residuals        16  26887    1680
kruskal.test(Tyramine_1)
##
##  Kruskal-Wallis rank sum test
##
## data:  res_1$Tyramine by res_1$Genotype
## Kruskal-Wallis chi-squared = 4.8286, df = 3, p-value = 0.1848
summary(aov(Putrescine_1))
##                Df Sum Sq Mean Sq F value Pr(>F)
## res_1$Genotype  3  10010    3337   1.031  0.405
## Residuals      16  51792    3237
kruskal.test(Spermidine_1)
##
##  Kruskal-Wallis rank sum test
##
## data:  res_1$Spermidine by res_1$Genotype
## Kruskal-Wallis chi-squared = 1.0114, df = 3, p-value = 0.7985
summary(aov(Tyramine_2))
##                Df Sum Sq Mean Sq F value  Pr(>F)
## res_2$Genotype  3   2653   884.3   8.281 0.00149 **
## Residuals      16   1709   106.8
## ---
## Signif. codes:  0 '***' 0.001 '**' 0.01 '*' 0.05 '.' 0.1 ' ' 1
TukeyHSD(aov(Tyramine_2))
##   Tukey multiple comparisons of means
##     95% family-wise confidence level
##
## Fit: aov(formula = Tyramine_2)
##
## $`res_2$Genotype`
##                 diff       lwr        upr     p adj
## EV-ETR1    -4.499390 -23.19820 14.1994171 0.9000047
## LOX3-ETR1 -27.036866 -45.73567 -8.3380595 0.0038776
## MYB8-ETR1 -22.671018 -41.36982 -3.9722111 0.0150612
## LOX3-EV   -22.537477 -41.23628 -3.8386699 0.0156941
## MYB8-EV   -18.171628 -36.87043  0.5271785 0.0583132
## MYB8-LOX3   4.365848 -14.33296 23.0646551 0.9075883
summary(aov(Putrescine_2))
##                Df Sum Sq Mean Sq F value Pr(>F)
## res_2$Genotype  3   3816    1272    0.62  0.612
## Residuals      16  32804    2050
summary(aov(Spermidine_2))
##                Df Sum Sq Mean Sq F value Pr(>F)
## res_2$Genotype  3  11766    3922   2.436  0.102
## Residuals      16  25755    1610
summary(aov(Tyramine_6l))
##                 Df Sum Sq Mean Sq F value   Pr(>F)
## res_6l$Genotype  3  18423    6141   11.62 0.000272 ***
## Residuals       16   8457     529
## ---
## Signif. codes:  0 '***' 0.001 '**' 0.01 '*' 0.05 '.' 0.1 ' ' 1
TukeyHSD(aov(Tyramine_6l))
##   Tukey multiple comparisons of means
##     95% family-wise confidence level
##
## Fit: aov(formula = Tyramine_6l)
##
## $`res_6l$Genotype`
##                 diff        lwr        upr     p adj
## EV-ETR1    30.863857  -10.73703  72.464745 0.1880536
## LOX3-ETR1 -38.668829  -80.26972   2.932059 0.0731962
## MYB8-ETR1 -43.542735  -85.14362  -1.941847 0.0386261
## LOX3-EV   -69.532686 -111.13357 -27.931798 0.0010524
## MYB8-EV   -74.406592 -116.00748 -32.805704 0.0005408
## MYB8-LOX3  -4.873906  -46.47479  36.726982 0.9865354
summary(aov(Putrescine_6l))
##                 Df Sum Sq Mean Sq F value Pr(>F)
## res_6l$Genotype  3   3044    1015   0.277  0.841
## Residuals       16  58684    3668
kruskal.test(Spermidine_6l)
##
##  Kruskal-Wallis rank sum test
##
## data:  res_6l$Spermidine by res_6l$Genotype
## Kruskal-Wallis chi-squared = 2.4514, df = 3, p-value = 0.4841
summary(aov(Tyramine_24l))
##                  Df Sum Sq Mean Sq F value   Pr(>F)
## res_24l$Genotype  3  18423    6141   11.62 0.000272 ***
## Residuals        16   8457     529
## ---
## Signif. codes:  0 '***' 0.001 '**' 0.01 '*' 0.05 '.' 0.1 ' ' 1
TukeyHSD(aov(Tyramine_24l))
##   Tukey multiple comparisons of means
##     95% family-wise confidence level
##
## Fit: aov(formula = Tyramine_24l)
##
## $`res_24l$Genotype`
##                 diff        lwr        upr     p adj
## EV-ETR1    30.863857  -10.73703  72.464745 0.1880536
## LOX3-ETR1 -38.668829  -80.26972   2.932059 0.0731962
## MYB8-ETR1 -43.542735  -85.14362  -1.941847 0.0386261
## LOX3-EV   -69.532686 -111.13357 -27.931798 0.0010524
## MYB8-EV   -74.406592 -116.00748 -32.805704 0.0005408
## MYB8-LOX3  -4.873906  -46.47479  36.726982 0.9865354
summary(aov(Putrescine_24l))
##                  Df Sum Sq Mean Sq F value Pr(>F)
## res_24l$Genotype  3   3044    1015   0.277  0.841
## Residuals        16  58684    3668
summary(aov(Spermidine_24l))
##                  Df Sum Sq Mean Sq F value Pr(>F)
## res_24l$Genotype  3   1562   520.7   0.569  0.643
## Residuals        16  14634   914.7
kruskal.test(Tyramine_48l)
##
##  Kruskal-Wallis rank sum test
##
## data:  res_48l$Tyramine by res_48l$Genotype
## Kruskal-Wallis chi-squared = 12.531, df = 3, p-value = 0.005768
pairwise.wilcox.test(res_48l$Tyramine, res_48l$Genotype)
##
##  Pairwise comparisons using Wilcoxon rank sum test
##
## data:  res_48l$Tyramine and res_48l$Genotype
##
##      ETR1  EV    LOX3
## EV   0.444 -     -
## LOX3 0.127 0.048 -
## MYB8 0.286 0.048 0.444
##
## P value adjustment method: holm
summary(aov(Putrescine_48l))
##                  Df Sum Sq Mean Sq F value Pr(>F)
## res_48l$Genotype  3  12866    4289   2.366  0.109
## Residuals        16  29005    1813
summary(aov(Spermidine_48l))
##                  Df Sum Sq Mean Sq F value Pr(>F)
## res_48l$Genotype  3    823   274.3   0.262  0.851
## Residuals        16  16721  1045.1
summary(aov(Tyramine_72l))
##                  Df Sum Sq Mean Sq F value   Pr(>F)
## res_72l$Genotype  3  479.7  159.90   15.64 5.13e-05 ***
## Residuals        16  163.5   10.22
## ---
## Signif. codes:  0 '***' 0.001 '**' 0.01 '*' 0.05 '.' 0.1 ' ' 1
TukeyHSD(aov(Tyramine_72l))
##   Tukey multiple comparisons of means
##     95% family-wise confidence level
##
## Fit: aov(formula = Tyramine_72l)
##
## $`res_72l$Genotype`
##                  diff        lwr       upr     p adj
## EV-ETR1     2.7154393  -3.069544  8.500423 0.5505733
## LOX3-ETR1  -8.3912282 -14.176212 -2.606244 0.0037750
## MYB8-ETR1  -8.0983435 -13.883327 -2.313360 0.0050699
## LOX3-EV   -11.1066675 -16.891651 -5.321684 0.0002598
## MYB8-EV   -10.8137829 -16.598767 -5.028799 0.0003440
## MYB8-LOX3   0.2928847  -5.492099  6.077868 0.9988688
summary(aov(Putrescine_72l))
##                  Df Sum Sq Mean Sq F value   Pr(>F)
## res_72l$Genotype  3  235.6   78.52   11.57 0.000278 ***
## Residuals        16  108.6    6.79
## ---
## Signif. codes:  0 '***' 0.001 '**' 0.01 '*' 0.05 '.' 0.1 ' ' 1
TukeyHSD(aov(Putrescine_72l))
##   Tukey multiple comparisons of means
##     95% family-wise confidence level
##
## Fit: aov(formula = Putrescine_72l)
##
## $`res_72l$Genotype`
##                 diff        lwr       upr     p adj
## EV-ETR1    0.5965000  -4.117447  5.310447 0.9831734
## LOX3-ETR1 -6.2227878 -10.936734 -1.508841 0.0080700
## MYB8-ETR1 -6.8534041 -11.567351 -2.139457 0.0037024
## LOX3-EV   -6.8192877 -11.533234 -2.105341 0.0038617
## MYB8-EV   -7.4499040 -12.163851 -2.735957 0.0017767
## MYB8-LOX3 -0.6306163  -5.344563  4.083330 0.9802512
summary(aov(Spermidine_72l))
##                  Df Sum Sq Mean Sq F value Pr(>F)
## res_72l$Genotype  3   8.95   2.982   1.243  0.327
## Residuals        16  38.39   2.399
kruskal.test(Tyramine_6s)
##
##  Kruskal-Wallis rank sum test
##
## data:  res_6s$Tyramine by res_6s$Genotype
## Kruskal-Wallis chi-squared = 4.8286, df = 3, p-value = 0.1848
summary(aov(Putrescine_6s))
##                 Df Sum Sq Mean Sq F value Pr(>F)
## res_6s$Genotype  3  19388    6463   3.074 0.0577 .
## Residuals       16  33641    2103
## ---
## Signif. codes:  0 '***' 0.001 '**' 0.01 '*' 0.05 '.' 0.1 ' ' 1
summary(aov(Spermidine_6s))
##                 Df Sum Sq Mean Sq F value Pr(>F)
## res_6s$Genotype  3   8785    2928    0.39  0.762
## Residuals       16 120042    7503
summary(aov(Tyramine_24s))
##                  Df Sum Sq Mean Sq F value Pr(>F)
## res_24s$Genotype  3  146.6   48.88   2.173  0.131
## Residuals        16  360.0   22.50
summary(aov(Putrescine_24s))
##                  Df Sum Sq Mean Sq F value Pr(>F)
## res_24s$Genotype  3  42111   14037   3.744 0.0327 *
## Residuals        16  59994    3750
## ---
## Signif. codes:  0 '***' 0.001 '**' 0.01 '*' 0.05 '.' 0.1 ' ' 1
TukeyHSD(aov(Putrescine_24s))
##   Tukey multiple comparisons of means
##     95% family-wise confidence level
##
## Fit: aov(formula = Putrescine_24s)
##
## $`res_24s$Genotype`
##                diff         lwr       upr     p adj
## EV-ETR1   -21.95264 -132.753919  88.84864 0.9404901
## LOX3-ETR1 -40.73789 -151.539165  70.06339 0.7223482
## MYB8-ETR1  79.70636  -31.094920 190.50764 0.2087323
## LOX3-EV   -18.78525 -129.586523  92.01603 0.9613159
## MYB8-EV   101.65900   -9.142278 212.46028 0.0780225
## MYB8-LOX3 120.44424    9.642967 231.24552 0.0308096
summary(aov(Spermidine_24s))
##                  Df Sum Sq Mean Sq F value Pr(>F)
## res_24s$Genotype  3  16857    5619   1.375  0.286
## Residuals        16  65387    4087
summary(aov(Tyramine_48s))
##                  Df Sum Sq Mean Sq F value   Pr(>F)
## res_48s$Genotype  3 170.90   56.97   10.83 0.000395 ***
## Residuals        16  84.18    5.26
## ---
## Signif. codes:  0 '***' 0.001 '**' 0.01 '*' 0.05 '.' 0.1 ' ' 1
TukeyHSD(aov(Tyramine_48s))
##   Tukey multiple comparisons of means
##     95% family-wise confidence level
##
## Fit: aov(formula = Tyramine_48s)
##
## $`res_48s$Genotype`
##                diff        lwr        upr     p adj
## EV-ETR1   -1.525879 -5.6763282  2.6245697 0.7223878
## LOX3-ETR1 -3.426428 -7.5768769  0.7240209 0.1253437
## MYB8-ETR1  4.490428  0.3399793  8.6408772 0.0317098
## LOX3-EV   -1.900549 -6.0509977  2.2499002 0.5700030
## MYB8-EV    6.016308  1.8658586 10.1667564 0.0037963
## MYB8-LOX3  7.916856  3.7664073 12.0673052 0.0002783
summary(aov(Putrescine_48s))
##                  Df Sum Sq Mean Sq F value Pr(>F)
## res_48s$Genotype  3   4051    1350   0.625  0.609
## Residuals        16  34548    2159
summary(aov(Spermidine_48s))
##                  Df Sum Sq Mean Sq F value Pr(>F)
## res_48s$Genotype  3   6477    2159    1.15  0.359
## Residuals        16  30032    1877
kruskal.test(Tyramine_72s)
##
##  Kruskal-Wallis rank sum test
##
## data:  res_72s$Tyramine by res_72s$Genotype
## Kruskal-Wallis chi-squared = 12.577, df = 3, p-value = 0.005646
pairwise.wilcox.test(res_72s$Tyramine, res_72s$Genotype)
##
##  Pairwise comparisons using Wilcoxon rank sum test
##
## data:  res_72s$Tyramine and res_72s$Genotype
##
##      ETR1  EV    LOX3
## EV   0.452 -     -
## LOX3 0.048 0.048 -
## MYB8 0.548 0.452 0.048
##
## P value adjustment method: holm
summary(aov(Putrescine_72s))
##                  Df Sum Sq Mean Sq F value Pr(>F)
## res_72s$Genotype  3  94.07   31.36   2.918 0.0662 .
## Residuals        16 171.92   10.74
## ---
## Signif. codes:  0 '***' 0.001 '**' 0.01 '*' 0.05 '.' 0.1 ' ' 1
summary(aov(Spermidine_72s))
##                  Df Sum Sq Mean Sq F value  Pr(>F)
## res_72s$Genotype  3  247.6   82.54   8.121 0.00164 **
## Residuals        16  162.6   10.16
## ---
## Signif. codes:  0 '***' 0.001 '**' 0.01 '*' 0.05 '.' 0.1 ' ' 1
TukeyHSD(aov(Spermidine_72s))
##   Tukey multiple comparisons of means
##     95% family-wise confidence level
##
## Fit: aov(formula = Spermidine_72s)
##
## $`res_72s$Genotype`
##                diff        lwr       upr     p adj
## EV-ETR1   -1.955088 -7.7238289  3.813654 0.7682348
## LOX3-ETR1  6.776972  1.0082312 12.545714 0.0187033
## MYB8-ETR1  4.774650 -0.9940908 10.543392 0.1240187
## LOX3-EV    8.732060  2.9633188 14.500801 0.0026147
## MYB8-EV    6.729738  0.9609968 12.498479 0.0196020
## MYB8-LOX3 -2.002322 -7.7710632  3.766419 0.7554091
```

### Statistical analysis of transcript levels by RT-qPCR

```
data = read.table("R_data.csv", header=T, sep=",", dec=".")

## Normality analysis

shapiro.test(subset(data$MYB8, data$Genotype=="EV"))
##
##  Shapiro-Wilk normality test
##
## data:  subset(data$MYB8, data$Genotype == "EV")
## W = 0.89797, p-value = 0.3987
shapiro.test(subset(data$MYB8, data$Genotype=="ETR1")) # NO
##
##  Shapiro-Wilk normality test
##
## data:  subset(data$MYB8, data$Genotype == "ETR1")
## W = 0.75797, p-value = 0.03522
shapiro.test(subset(data$MYB8, data$Genotype=="LOX3"))
##
##  Shapiro-Wilk normality test
##
## data:  subset(data$MYB8, data$Genotype == "LOX3")
## W = 0.98502, p-value = 0.9595
shapiro.test(subset(data$MYB8, data$Genotype=="MYB8"))
##
##  Shapiro-Wilk normality test
##
## data:  subset(data$MYB8, data$Genotype == "MYB8")
## W = 0.77738, p-value = 0.05233
shapiro.test(subset(data$AT1, data$Genotype=="EV"))
##
##  Shapiro-Wilk normality test
##
## data:  subset(data$AT1, data$Genotype == "EV")
## W = 0.92224, p-value = 0.5496
shapiro.test(subset(data$AT1, data$Genotype=="ETR1"))
##
##  Shapiro-Wilk normality test
##
## data:  subset(data$AT1, data$Genotype == "ETR1")
## W = 0.91799, p-value = 0.5171
shapiro.test(subset(data$AT1, data$Genotype=="LOX3"))
##
##  Shapiro-Wilk normality test
##
## data:  subset(data$AT1, data$Genotype == "LOX3")
## W = 0.89121, p-value = 0.3632
shapiro.test(subset(data$AT1, data$Genotype=="MYB8"))
##
##  Shapiro-Wilk normality test
##
## data:  subset(data$AT1, data$Genotype == "MYB8")
## W = 0.89744, p-value = 0.3959
shapiro.test(subset(data$DH29, data$Genotype=="EV"))
##
##  Shapiro-Wilk normality test
##
## data:  subset(data$DH29, data$Genotype == "EV")
## W = 0.84189, p-value = 0.1702
shapiro.test(subset(data$DH29, data$Genotype=="ETR1")) # NO
##
##  Shapiro-Wilk normality test
##
## data:  subset(data$DH29, data$Genotype == "ETR1")
## W = 0.73332, p-value = 0.0207
shapiro.test(subset(data$DH29, data$Genotype=="LOX3"))
##
##  Shapiro-Wilk normality test
##
## data:  subset(data$DH29, data$Genotype == "LOX3")
## W = 0.91975, p-value = 0.5283
shapiro.test(subset(data$DH29, data$Genotype=="MYB8"))
##
##  Shapiro-Wilk normality test
##
## data:  subset(data$DH29, data$Genotype == "MYB8")
## W = 0.94899, p-value = 0.73
shapiro.test(subset(data$CV86, data$Genotype=="EV"))
##
##  Shapiro-Wilk normality test
##
## data:  subset(data$CV86, data$Genotype == "EV")
## W = 0.90868, p-value = 0.4597
shapiro.test(subset(data$CV86, data$Genotype=="ETR1")) # NO
##
##  Shapiro-Wilk normality test
##
## data:  subset(data$CV86, data$Genotype == "ETR1")
## W = 0.6962, p-value = 0.008736
shapiro.test(subset(data$CV86, data$Genotype=="LOX3"))
##
##  Shapiro-Wilk normality test
##
## data:  subset(data$CV86, data$Genotype == "LOX3")
## W = 0.99458, p-value = 0.993
shapiro.test(subset(data$CV86, data$Genotype=="MYB8"))
##
##  Shapiro-Wilk normality test
##
## data:  subset(data$CV86, data$Genotype == "MYB8")
## W = 0.83714, p-value = 0.1572
shapiro.test(subset(data$PAL2, data$Genotype=="EV"))
##
##  Shapiro-Wilk normality test
##
## data:  subset(data$PAL2, data$Genotype == "EV")
## W = 0.87934, p-value = 0.3063
shapiro.test(subset(data$PAL2, data$Genotype=="ETR1"))
##
##  Shapiro-Wilk normality test
##
## data:  subset(data$PAL2, data$Genotype == "ETR1")
## W = 0.94851, p-value = 0.7265
shapiro.test(subset(data$PAL2, data$Genotype=="LOX3"))
##
##  Shapiro-Wilk normality test
##
## data:  subset(data$PAL2, data$Genotype == "LOX3")
## W = 0.94037, p-value = 0.6685
shapiro.test(subset(data$PAL2, data$Genotype=="MYB8"))
##
##  Shapiro-Wilk normality test
##
## data:  subset(data$PAL2, data$Genotype == "MYB8")
## W = 0.93261, p-value = 0.6143
## Variance analysis

MYB8_2 = formula(data$MYB8~data$Genotype)
fligner.test(MYB8_2)
##
##  Fligner-Killeen test of homogeneity of variances
##
## data:  data$MYB8 by data$Genotype
## Fligner-Killeen:med chi-squared = 4.6199, df = 3, p-value = 0.2018
AT1_2 = formula(data$AT1~data$Genotype)
fligner.test(AT1_2)
##
##  Fligner-Killeen test of homogeneity of variances
##
## data:  data$AT1 by data$Genotype
## Fligner-Killeen:med chi-squared = 6.511, df = 3, p-value = 0.08923
DH29_2 = formula(data$DH29~data$Genotype)
fligner.test(DH29_2)
##
##  Fligner-Killeen test of homogeneity of variances
##
## data:  data$DH29 by data$Genotype
## Fligner-Killeen:med chi-squared = 5.3233, df = 3, p-value = 0.1496
CV86_2 = formula(data$CV86~data$Genotype)
fligner.test(CV86_2)
##
##  Fligner-Killeen test of homogeneity of variances
##
## data:  data$CV86 by data$Genotype
## Fligner-Killeen:med chi-squared = 5.3128, df = 3, p-value = 0.1503
PAL2_2 = formula(data$PAL2~data$Genotype)
fligner.test(PAL2_2)
##
##  Fligner-Killeen test of homogeneity of variances
##
## data:  data$PAL2 by data$Genotype
## Fligner-Killeen:med chi-squared = 2.949, df = 3, p-value = 0.3996
## ANOVA

kruskal.test(MYB8_2)
##
##  Kruskal-Wallis rank sum test
##
## data:  data$MYB8 by data$Genotype
## Kruskal-Wallis chi-squared = 14.703, df = 3, p-value = 0.002089
pairwise.wilcox.test(data$MYB8, data$Genotype)
##
##  Pairwise comparisons using Wilcoxon rank sum test
##
## data:  data$MYB8 and data$Genotype
##
##      ETR1  EV    LOX3
## EV   0.095 -     -
## LOX3 0.048 0.048 -
## MYB8 0.095 0.048 1.000
##
## P value adjustment method: holm
summary(aov(AT1_2))
##               Df Sum Sq Mean Sq F value  Pr(>F)
## data$Genotype  3  3.456  1.1522   36.82 3.7e-07 ***
## Residuals     15  0.469  0.0313
## ---
## Signif. codes:  0 '***' 0.001 '**' 0.01 '*' 0.05 '.' 0.1 ' ' 1
## 1 observation deleted due to missingness
TukeyHSD(aov(AT1_2))
##   Tukey multiple comparisons of means
##     95% family-wise confidence level
##
## Fit: aov(formula = AT1_2)
##
## $`data$Genotype`
##                  diff         lwr        upr     p adj
## EV-ETR1    0.29593637 -0.04606235  0.6379351 0.1016269
## LOX3-ETR1 -0.67273939 -0.99517887 -0.3502999 0.0001256
## MYB8-ETR1 -0.72229682 -1.04473630 -0.3998573 0.0000577
## LOX3-EV   -0.96867576 -1.31067448 -0.6266770 0.0000036
## MYB8-EV   -1.01823319 -1.36023191 -0.6762345 0.0000020
## MYB8-LOX3 -0.04955743 -0.37199692  0.2728821 0.9699761
kruskal.test(DH29_2)
##
##  Kruskal-Wallis rank sum test
##
## data:  data$DH29 by data$Genotype
## Kruskal-Wallis chi-squared = 15.754, df = 3, p-value = 0.001273
pairwise.wilcox.test(data$DH29, data$Genotype)
##
##  Pairwise comparisons using Wilcoxon rank sum test
##
## data:  data$DH29 and data$Genotype
##
##      ETR1  EV    LOX3
## EV   0.190 -     -
## LOX3 0.048 0.048 -
## MYB8 0.048 0.048 0.190
##
## P value adjustment method: holm
kruskal.test(CV86_2)
##
##  Kruskal-Wallis rank sum test
##
## data:  data$CV86 by data$Genotype
## Kruskal-Wallis chi-squared = 16.577, df = 3, p-value = 0.0008633
pairwise.wilcox.test(data$CV86, data$Genotype)
##
##  Pairwise comparisons using Wilcoxon rank sum test
##
## data:  data$CV86 and data$Genotype
##
##      ETR1  EV    LOX3
## EV   0.063 -     -
## LOX3 0.048 0.048 -
## MYB8 0.048 0.048 0.063
##
## P value adjustment method: holm
summary(aov(PAL2_2))
##               Df Sum Sq Mean Sq F value   Pr(>F)
## data$Genotype  3  91.65  30.551   40.46 1.07e-07 ***
## Residuals     16  12.08   0.755
## ---
## Signif. codes:  0 '***' 0.001 '**' 0.01 '*' 0.05 '.' 0.1 ' ' 1
TukeyHSD(aov(PAL2_2))
##   Tukey multiple comparisons of means
##     95% family-wise confidence level
##
## Fit: aov(formula = PAL2_2)
##
## $`data$Genotype`
##                diff        lwr        upr     p adj
## EV-ETR1    1.104038 -0.4683829  2.6764583 0.2257102
## LOX3-ETR1 -2.166949 -3.7393692 -0.5945280 0.0057563
## MYB8-ETR1 -4.484287 -6.0567071 -2.9118659 0.0000024
## LOX3-EV   -3.270986 -4.8434069 -1.6985657 0.0001085
## MYB8-EV   -5.588324 -7.1607448 -4.0159036 0.0000001
## MYB8-LOX3 -2.317338 -3.8897586 -0.7449173 0.0032979
```
