## Supplemental Table S1 for "Ethylene is a local modulator of jasmonate-dependent phenolamide accumulation during *Manduca sexta* herbivory in *Nicotiana attenuata*"

|  |  |  | Fold change in systemic leaves 72h after simulated herbivory (unpaired *t*-test) | | | |
| --- | --- | --- | --- | --- | --- | --- |
| Compound | **Precursor *m/z*** | **Retention time** | **W+OS inducibility** (W+OS WT 72h > WT 0h) | **JA regulation** (W+OS asLOX3 72h > W+OS WT 72h) | **ET regulation** (W+OS sETR1 72h > W+OS WT 72h) | **MYB8 regulation** (W+OS irMYB8 72h > W+OS WT 72h) |
| *N*-Coumaroylputrescine | 235.14 | 161s | **364.32**  (**) | *0.01*  (**) | 1.30 | *0.00*  (**) |
|  |  | 201s | **129.98**  (***) | *0.02*  (***) | 1.00 | *0.00*  (***) |
| *N*-Caffeoylputrescine | 251.14 | 114s | **9.99**  (**) | 1.00 | *0.60*  (*) | *0.01*  (**) |
|  |  | 132s | **42.51**  (**) | *0.07*  (**) | 0.86 | *0.00*  (**) |
| *N*-Feruloylputrescine | 265.15 | 195s | **22.47**  (***) | *0.10*  (***) | 0.80 | *0.00*  (***) |
|  |  | 248s | **20.30**  (***) | *0.08*  (***) | 1.01 | *0.00*  (***) |
| Putative acylated  *N*-Caffeoylputrescine | 347.19 | 542s | **333.05**  (***) | *0.12*  (***) | *0.74*  (*) | *0.00*  (***) |
|  |  | 627s | **1078.62**  (***) | *0.02*  (**) | 0.71 | *0.00*  (***) |
| *N*-Caffeoylspermidine | 308.20 | 366s | *0.01*  (*) | **224.45**  (*) | 2.20 | 2.86 |
|  |  | 414s | *0.23*  (*) | 3.09 | 0.61 | *0.03*  (**) |
|  |  | 450s | 0.54 | 0.59 | 0.63 | *0.00*  (**) |
| *N'*-*N"*-Dicoumaroylspermidine | 438.20 | 240s | **112.91**  (*) | 0.00 | *0.54*  (*) | *0.00*  (*) |
| *N'*-*N"*-Coumaroylcaffeoylspermidine | 454.23 | 462s | *0.21*  (*) | **8.40**  (**) | 0.60 | *0.22*  (*) |
|  |  | 537s | **3.71**  (**) | *0.16*  (***) | 0.82 | *0.00*  (**) |
| *N'*-*N"*-Dicaffeoylspermidine | 470.23 | 365s | *0.09*  (*) | **17.41**  (*) | 1.73 | *0.24*  (**) |
|  |  | 412s | *0.30*  (*) | 2.63 | 0.46 | 0.03  (**) |
|  |  | 449s | 0.55 | 0.70 | 0.62 | 0.01  (**) |
| *N'*-*N"*-Feruloylcaffeoylspermidine | 484.24 | 492s | *0.16*  (**) | **6.67**  (**) | *0.27*  (**) | 0.96 |
|  |  | 526s | 0.68 | 1.07 | 0.74 | 0.06  (***) |
|  |  | 550s | *0.09*  (*) | **9.79**  (**) | 0.43 | 0.82 |
|  |  | 573s | 0.34 | 1.14 | 0.55 | 0.05  (**) |
| *N'*-*N"*-Diferuloylspermidine | 498.26 | 623s | 1.00 | 1.15 | *0.67*  (*) | *0.07*  (***) |
|  |  | 664s | 1.11 | 1.03 | *0.70*  (**) | *0.06*  (***) |
|  |  | 696s | 0.88 | *0.74*  (*) | *0.58*  (***) | *0.03*  (***) |
| Unknown  *N'*-*N"*-Diacylatedspermidine | 566.28 | 759s | 0.53 | 1.60 | *0.48*  (*) | *0.02*  (**) |
|  |  | 807s | 0.23 | **4.84**  (**) | 0.54 | *0.02*  (**) |
| Phenylalanine | 166.09 | 126s | 0.92 | *0.81*  (*) | 1.36 | 0.77 |
| Tyrosine | 182.08 | 96s | *0.53*  (*) | **1.36**  (*) | 1.16 | 1.09 |
| Cinnamic acid | 148.06 | 71s | *0.30*  (**) | **1.65**  (**) | 1.11 | 1.08 |
| Coumaric acid | 147.04 | 528s | *0.42*  (**) | **2.01**  (*) | 0.77 | 5.22 |
| Coumaroyl quinate | 339.11 | 420s | *0.41*  (**) | **1.98**  (*) | 0.49 | 4.24 |
|  |  | 528s | *0.13*  (*) | 6.30 | 1.11 | 18.71 |
| Chlorogenic acid  (Caffeoyl quinate) | 355.10 | 294s | *0.60*  (*) | 1.13 | 0.84 | *0.58*  (**) |
|  |  | 396s | *0.44*  (**) | 1.65 | *0.83*  (*) | 1.01 |
| Rutin | 611.16 | 636s | 0.97 | 1.10 | *0.76*  (*) | 1.08 |
| Nicotine | 163.12 | 72s | **2.52**  (***) | *0.53*  (***) | *0.75*  (*) | 0.96 |
|  |  | 120s | **3.46**  (**) | *0.57*  (**) | 0.57 | 0.88 |
| Attenoside | 956.50 | 1338s | 0.70 | 1.60 | 0.69 | **1.67**  (*) |
| Nicotianoside I | 885.41 | 1362s | **16.54**  (**) | *0.11*  (**) | 1.21 | *0.12*  (**) |
| Nicotianoside VII | 271.24 | 1290s | **2.31**  (*) | 0.62 | *0.05*  (**) | 0.87 |
|  |  | 1356s | 1.08 | 1.10 | 1.10 | **1.35**  (*) |
| Lyciumoside I | 625.32 | 1368s | 0.43 | 3.44 | 1.04 | **2.92**  (*) |
| Lyciumoside II | 810.44 | 1134s | 0.95 | 4.34 | 0.25 | **9.97**  (**) |
|  |  | 1152s | 0.75 | 3.94 | 0.05 | **7.53**  (**) |
|  |  | 1200s | 1.02 | 2.36 | 0.25 | **5.06**  (**) |
|  |  | 1218s | 1.19 | 1.77 | 0.44 | **4.15**  (**) |
|  |  | 1266s | 1.97 | 1.54 | 0.44 | **4.98**  (**) |
|  |  | 1344s | *0.29*  (*) | **2.51**  (**) | 0.78 | **2.63**  (***) |
| Lyciumoside IV | 471.17 | 1356s | 1.00 | 1.23 | 0.89 | **1.40**  (*) |
